# Discovery of First-in-Class PROTAC Degraders of SARS-CoV-2 Main Protease

**DOI:** 10.1101/2023.09.29.560163

**Authors:** Yugendar R. Alugubelli, Jing Xiao, Kaustav Khatua, Sathish Kumar, Yuying Ma, Xinyu R. Ma, Veerabhadra R. Vulupala, Sandeep R. Atla, Lauren Blankenship, Demonta Coleman, Benjamin W. Neuman, Wenshe Ray Liu, Shiqing Xu

## Abstract

We have witnessed three coronavirus (CoV) outbreaks in the past two decades, including the COVID-19 pandemic caused by SARS-CoV-2. Main protease (M^Pro^) is a highly conserved and essential protease that plays key roles in viral replication and pathogenesis among various CoVs, representing one of the most attractive drug targets for antiviral drug development. Traditional antiviral drug development strategies focus on the pursuit of high-affinity binding inhibitors against M^Pro^. However, this approach often suffers from issues such as toxicity, drug resistance, and a lack of broad-spectrum efficacy. Targeted protein degradation represents a promising strategy for developing next-generation antiviral drugs to combat infectious diseases. Here we leverage the proteolysis targeting chimera (PROTAC) technology to develop a new class of small-molecule antivirals that induce the degradation of SARS-CoV-2 M^Pro^. Our previously developed M^Pro^ inhibitors MPI8 and MPI29 were used as M^Pro^ ligands to conjugate a CRBN E3 ligand, leading to compounds that can both inhibit and degrade SARS-CoV-2 M^Pro^. Among them, MDP2 was demonstrated to effectively reduce M^Pro^ protein levels in 293T cells (DC_50_ = 296 nM), relying on a time-dependent, CRBN-mediated, and proteasome-driven mechanism. Furthermore, MPD2 exhibited remarkable efficacy in diminishing M^Pro^ protein levels in SARS-CoV-2-infected A549-ACE2 cells, concurrently demonstrating potent anti-SARS-CoV-2 activity (EC_50_ = 492 nM). This proof-of-concept study highlights the potential of PROTAC-mediated targeted protein degradation of M^Pro^ as an innovative and promising approach for COVID-19 drug discovery.

## INTRODUCTION

Coronaviruses (CoVs) are a family of single-stranded, positive-sense RNA viruses with a wide-ranging ability to infect mammals including humans and birds. They are responsible for a spectrum of diseases, including respiratory, hepatic, enteric, and neurologic diseases.^1-5^ Prior to 2003, only a few CoVs such as HCoV-229E and HCoV-OC43^6-7^ were recognized as human pathogens. However, within a span of just two decades, the world witnessed the emergence of three major severe respiratory disease outbreaks: the severe acute respiratory syndrome coronavirus (SARS-CoV) in 2003,^8^ Middle East respiratory syndrome coronavirus (MERS-CoV) in 2012,^9^ and SARS-CoV-2 in 2019.^10-13^ These outbreaks swiftly evolved into significant public health crises. In particular, the ongoing coronavirus disease 2019 (COVID-19) pandemic caused by SARS-CoV-2 has evolved into one of the most formidable public health challenges in human history. According to statistics released from the World Health Organization (WHO) on August 23, 2023, the numbers of confirmed cases and deaths of COVID-19 worldwide have exceeded 770 million and 6.9 million, respectively. The persistent recurrence of CoV pandemics over the past two decades has led scientists to believe that new CoV strains will periodically emerge in human populations. This phenomenon is attributed to various factors, including the high prevalence and widespread distribution of CoVs, their substantial genetic diversity, and the ever-increasing close human-animal interactions in modern society.^14-15^ Given the ongoing COVID-19 pandemic and the potential for future CoV outbreaks, it is imperative to prioritize the development of orally available small-molecule drugs that can be easily distributed as effective CoV antivirals for both treatment and prevention.

The SARS-CoV-2 genome encodes conserved replicase and structural proteins, plus several small open reading frames (ORFs) that encode accessory proteins dispensable for virus growth. ORF1ab, the largest ORF, contains overlapping open reading frames that encode polyproteins pp1ab and pp1a.^16^ These polyproteins are cleaved to yield 16 nonstructural proteins (nsps), nsp1-16, through their two viral proteases: papain-like proteinase protein (PL^Pro^, nsp3) and 3C-like (3CL^Pro^, nsp5) that is also called main protease (M^Pro^, nsp5).^17^ These nsps play a pivotal role in viral transcription, replication, proteolytic processing, suppression of host immune responses and host gene expression. Notably, M^Pro^ is responsible for processing 13 out of the 16 critical nsps that are indispensable for viral replication and packaging. Furthermore, M^Pro^ exhibits a high degree of conservation among various CoVs, making it one of the most attractive drug targets for developing antivirals against COVID-19.^18-20^ M^Pro^ is a cysteine protease with three domains, and its active form is a homodimer comprising two protomers. Each protomer contains a Cys145-His41 catalytic dyad, where cysteine serves as the nucleophile in the proteolytic process.^21^ Consequently, the primary strategy for the development of antivirals has centered on high-affinity ligands that can bind directly to M^Pro^, thereby inhibiting its enzymatic activities and functions. A significant milestone in M^Pro^ inhibitors was the development of nirmatrelvir, an orally available reversible covalent SARS-CoV-2 M^Pro^ inhibitor developed at Pfizer.^22^ In December 2021, the US FDA granted emergency use authorization for Paxlovid, a combination therapy comprising nirmatrelvir and ritonavir, for the treatment of COVID-19. By May 2023, Paxlovid received full FDA approval for use in high-risk adults, and has been granted conditional or emergency use authorization in more than 70 countries worldwide for combating COVID-19. While Paxlovid has generated significant excitement, it is essential to acknowledge several associated issues. These include but are not limited to the potential for serious side effects, drug interactions, the COVID-19 rebound likely due to inadequate drug exposure, and the emergence of drug resistance associated with naturally occurring mutations of SARS-CoV-2 M^Pro^.^23-25^ Given these challenges, there remains a critical need for novel antiviral therapies, particularly those with alternative mechanisms of action that can effectively address these concerns, in the ongoing battle against SARS-CoV-2 and newly emerging CoV pathogens.

Proteolysis targeting chimera (PROTAC) represents an innovative technology for targeted protein degradation in precision medicine.^26-31^ These rationally designed small-molecule PROTACs are heterobifunctional compounds composed of two active ligands interconnected by a chemical linker. One of these ligands is tailored to bind specifically to a protein of interest (POI) while the other selectively engages an E3 ubiquitin ligase. The recruitment of the E3 ligase to the target protein facilitates the formation of a ternary complex, leading to the ubiquitination and ultimate degradation of the target protein.^26-31^ PROTACs offer numerous advantages over traditional occupancy-based inhibitors, including: (i) the ability to degrade multi-domain proteins, eliminating both enzymatic and non-enzymatic/scaffolding functions;^30^ (ii) flexibility in recruiting targets via various binding sites, even in cases where sustained inhibition is unnecessary. This enables the degradation of “undruggable” targets, using allosteric sites if needed;^32-33^ (iii) the potential to transform weak binders into potent degraders; (iv) a catalytic nature allowing for sub-stoichiometric activity and improved efficacy;^34-36^ (v) enhanced target selectivity achieved through protein–protein interactions between the targeted protein and the recruited E3 ligase;^35-40^ (vi) capability to overcome drug resistance stemming from target protein overexpression by degrading the full-length protein;^41-43^ and (vii) enhanced intracellular accumulation and target engagement.^44^ These distinctive features position PROTAC as a potential game-changing technology in drug discovery that has been widely explored for degrading various disease-causing proteins. Harnessing these advantages, antiviral PROTACs via targeted protein degradation has been considered as a promising strategy for developing nextgeneration antiviral drugs to combat infectious diseases.^45-48^ The first antiviral PROTAC has been reported for degrading hepatitis C virus (HCV) NS3/4A protease with superior antiviral activity and resistance profiles to those of traditional drug telaprevir.^49^ However, to the best of our knowledge, there has been no report of a small-molecule PROTAC degrader targeting M^Pro^ for combating COVID-19 to date. In our ongoing pursuit of COVID-19 drug discovery targeting M^Pro^, we present the design, synthesis, and evaluation of the first series of small-molecule PROTAC degraders targeting SARS-CoV-2 M^Pro^. Our work has led to the identification of MPD2 as a potent degrader of M^Pro^ with a DC_50_ value of 296 nM, demonstrating promising antiviral activity against SARS-CoV-2 with an EC_50_ value of 492 nM.

## RESULTS and DISCUSSION

### Design of M^Pro^ PROTAC degraders (MPDs) based on reversible covalent inhibitors MPI8 and MPI29

M^Pro^, a protein consisting of 306 amino acids, features an active site comprising four small pockets responsible for binding to the P1, P2, P4, and P1’-3’ residues within a protein substrate.^50^ Crucially, M^Pro^ specifically recognizes a protein substrate containing a strictly P1 glutamine residue. Two active site residues Cys145 and His41 form a catalytic dyad.^51^ An effective strategy for developing M^Pro^ inhibitors involves the conversion of the hydrolytic peptide bond within a substrate into an electrophilic warhead (e.g. aldehyde) to facilitates the formation of a covalent adduct with the catalytic Cys145 of M^Pro^.^20, 52^ In 2020, building upon insights from prior medicinal chemistry studies on SARS-CoV M^Pro^ inhibitors,^20, 52^ we embarked on the design and synthesis of new reversible covalent inhibitors for SARS-CoV-2 M^Pro^ by: (i) employing a more potent β-(S-2-oxopyrrolidin-3-yl)-alanine as the fixed P1 residue and enhancing binding due to reduced entropy loss when M^Pro^ engages with the more rigid lactam; (ii) utilizing aldehyde group as an electrophilic warhead to form a reversible covalent bond with active site Cys145; (iii) optimizing the P2, P3 and P4 positions for improved potency.^50^ We characterized their enzymatic inhibition and antiviral potency, and determined the crystal structures of M^Pro^ bound with these inhibitors. Among these inhibitors, MPI8 emerged with the highest cellular potency against M^Pro^ and highest antiviral activity against SARS-CoV-2.^50, 53^ X-ray crystallography analysis of the M^Pro^-MPI8 complex (PDB ID: 7JQ5) revealed that MPI8 fits precisely in the P1- and P2-binding pockets at the M^Pro^ active site(Figure 1a).^50^ Strong van der Waals interactions at the P1- and P2-binding pockets, a number of hydrogen bonds with activesite residues, and the covalent interaction with C145 to form a hemithioacetal adduct contribute to high affinity of MPI8 binding to M^Pro^. The *N*-terminal phenyl group of MPI8 and other inhibitors are not well defined in the crystal structures, indicating an unfitting size or relatively loosely bound pattern within P4-binding pocket (Figure 1a).

**Figure 1.**
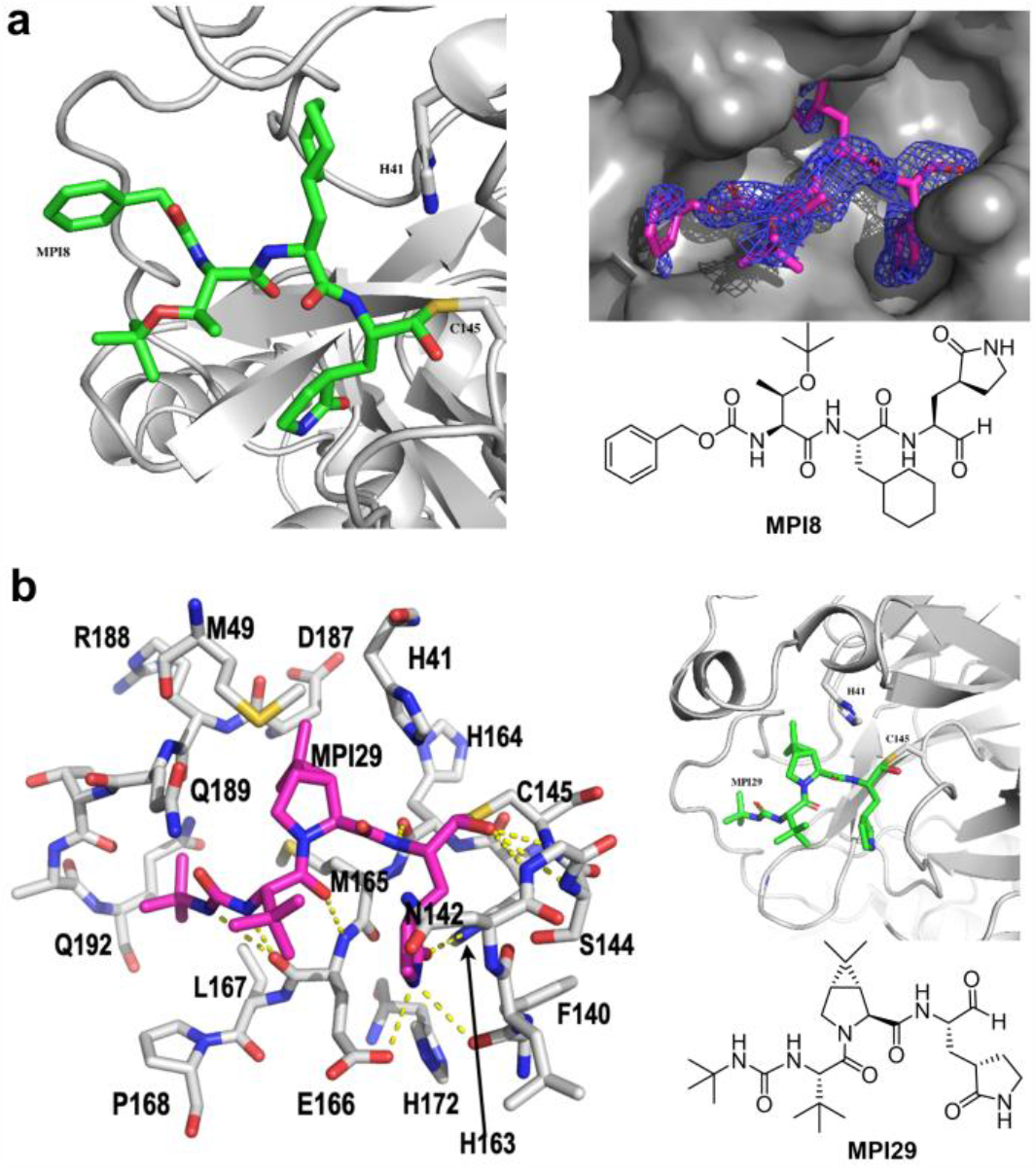
Representative reversible covalent inhibitors MPI8 and MPI29 of SARS-CoV-2 M^Pro^. **a**, M^Pro^-MPI8 structure (PDB: 7JQ5).^50^ **b**, M^Pro^-MPI29 structure (PDB: 7S6W).^57^

Boceprevir has been previously explored as a repurposed drug for COVID-19.^54-56^ It serves as an M^Pro^ inhibitor and contains an α-ketoamide warhead, a P1 β-cyclobutylalanyl moiety, a P2 dimethylcyclopropylproline, a P3 *tert*-butylglycine, and an *N*-terminal *tert*-butylcarbamide. In 2020, we designed and synthesized MPI29 by replacing the P1 site of Boceprevir with a 5-oxopyrrolidine-containing residue and altering the warhead to an aldehyde, resulting in an incredibly potent enzymatic inhibitor (IC_50_ = 9.3 nM).^57^ In the M^Pro^-MPI29 complex (PDB ID: 7S6W) as depicted in Figure 1b, MPI29 forms a covalent adduct with M^Pro^ C145 and multiple hydrogen bonds with M^Pro^. Notably, the amide of the lactam at P1 side chain of MPI29 engages in three hydrogen bonds with M^Pro^ residues including F140, H163, and E166. Additionally, two hydrogen bonds are generated between two backbone amides of MPI29 and M^Pro^ residues involving H164 and E166. The P4 *N*-terminal urea cap of MPI29 employs its two nitrogen atoms to form hydrogen bonds with the backbone oxygen of M^Pro^ E166 (Figure 1b).^57^ Recent studies revealed that PROTAC degraders based on reversible covalent ligands not only exhibit enhanced selectivity and binding affinity, but also retain the catalytic ability to facilitate multiple rounds of POI degradation.^44, 58^ Hence, we chose to utilize reversible covalent M^Pro^ ligands MPI8 and MPI29 for the development of covalent PROTAC degraders for M^Pro^ by taking the advantages of the strengths of reversible covalent inhibitors and event-driven PROTAC technology.

The crystal structures of M^Pro^-MPI8 complex (PDB: 7JQ5) and M^Pro^-MPI29 complex (PDB: 7S6W) provided a structural basis to rationally choose a link site for the design of PROTAC degraders. We initiated our research by exploring ubiquitin E3 ligase cereblon (CRBN) ligands as the E3 ligase binders. Drawing insights from crystal structures of M^Pro^-MPI8 complex (PDB: 7JQ5), it became apparent that the *N*-terminal phenyl moiety of MPI8 exhibited a less defined binding pattern within the P4-binding pocket (Figure 1a). These observations guided the design of a series of M^Pro^ degraders, achieved through the incorporation of a ligand for the CRBN E3 ligase using a diverse array of linkers (Figure 2a): (i) modification of the *N*-terminal phenyl group of MPI8 with different linkers; and (ii) substitution of *N*-terminal phenyl group of MPI8 with a triazole via Cu-catalyzed azide–alkyne cycloaddition with different linkers. Building on crystal structures of M^Pro^-MPI29 complex (PDB: 7S6W), we identified the *N*-terminal *tert*-butyl moiety of MPI29 at the P4 site as a suitable linker site for introducing CRBN ligand without interrupting its binding with M^Pro^ (Figure 2b).

**Figure 2.**
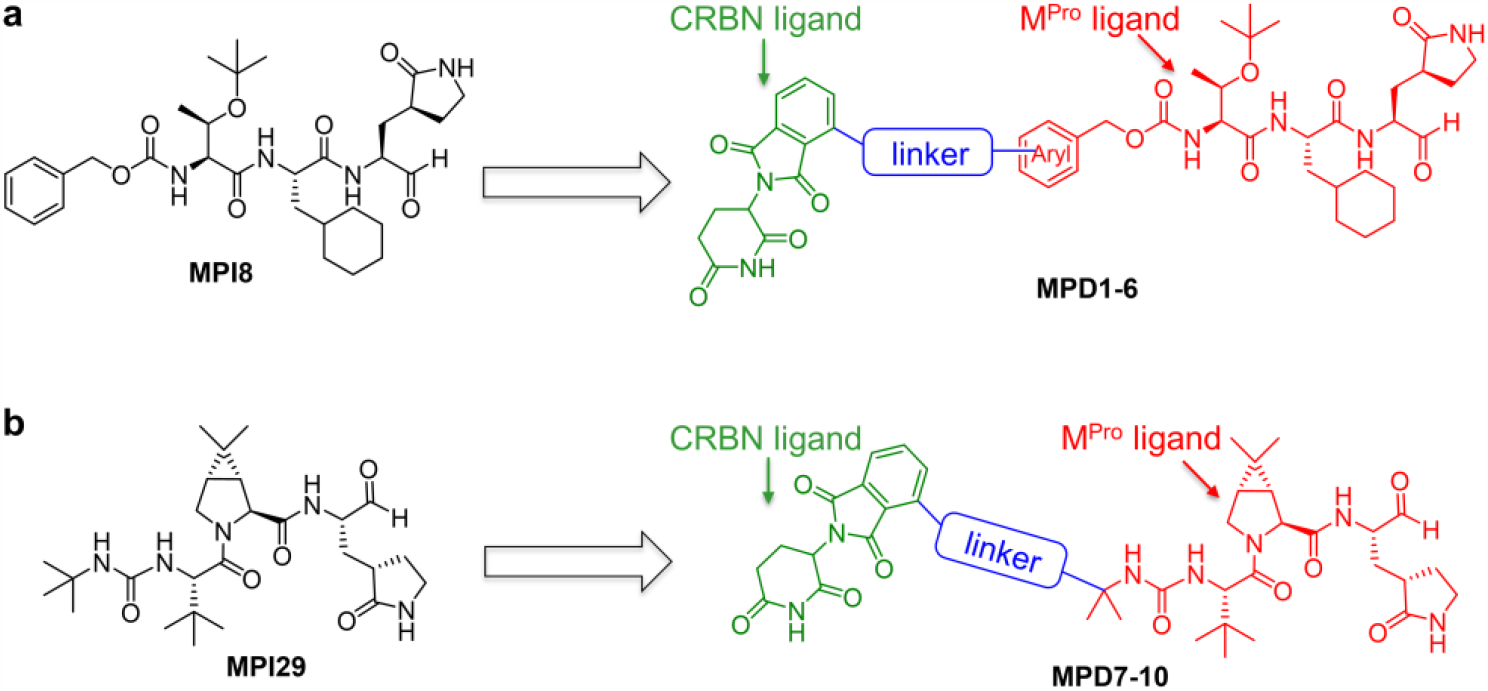
Design of M^Pro^ PROTAC degraders based on reversible covalent inhibitors MPI8 and MPI29. **a**, MPD1-6 via linking a CRBN ligand at *N*-terminal phenyl group at P4 site of MPI8. **b**, MPD7-10 via linking a CRBN ligand at the *N*-terminal *tert-*butyl group of MPI29.

We synthesized a series of bifunctional small-molecule PROTACs, namely MPD1-MPD6 based on MPI8 and MPD7-MPD10 based on MPI29, as shown in Figure 3a and Figure S1. The CRBN-binding moieties of MPD1-MPD3 and MPD4 were derived from thalidomide-4-OH and pomalidomide, respectively, two widely used CRBN binders for the PROTAC development.^59^ For MPD5-MPD6, we replaced *N*-terminal phenyl group of MPI8 with a triazole and introduced pomalidomide CRBN ligand with different linkers via Cu-catalyzed azide–alkyne cycloaddition. For MPI29-based PROTACs MPD7-10, we modified one of methyl group of *N*-terminal *tertbutyl* moiety of MPI29 to introduce thalidomide-4-OH and pomalidomide CRBN ligands (Figure 3a). To assess the inhibitory potency of MPD1-MPD10, we used a previously established protocol that uses Sub3 (Dabcyl-KTSAVLQSGFRKME-Edans) (Figure S2), a fluorogenic peptide substrate of M^Pro^.^60^ In this assay, we preincubated M^Pro^ with a PROTAC molecule for 30 min before Sub3 was added and the fluorescent product formation (Ex: 336 nm/Em: 490 nm) was recorded in a fluorescence plate reader. As illustrated in Figure 3b, all MPD1-MPD10 retained sub-micromolar IC_50_ values for the inhibition of SARS-CoV-2 M^Pro^, which indicates that our structure-based design of linker sites works well as there were no obvious interferences with MPDs binding to M^Pro^.

**Figure 3.**
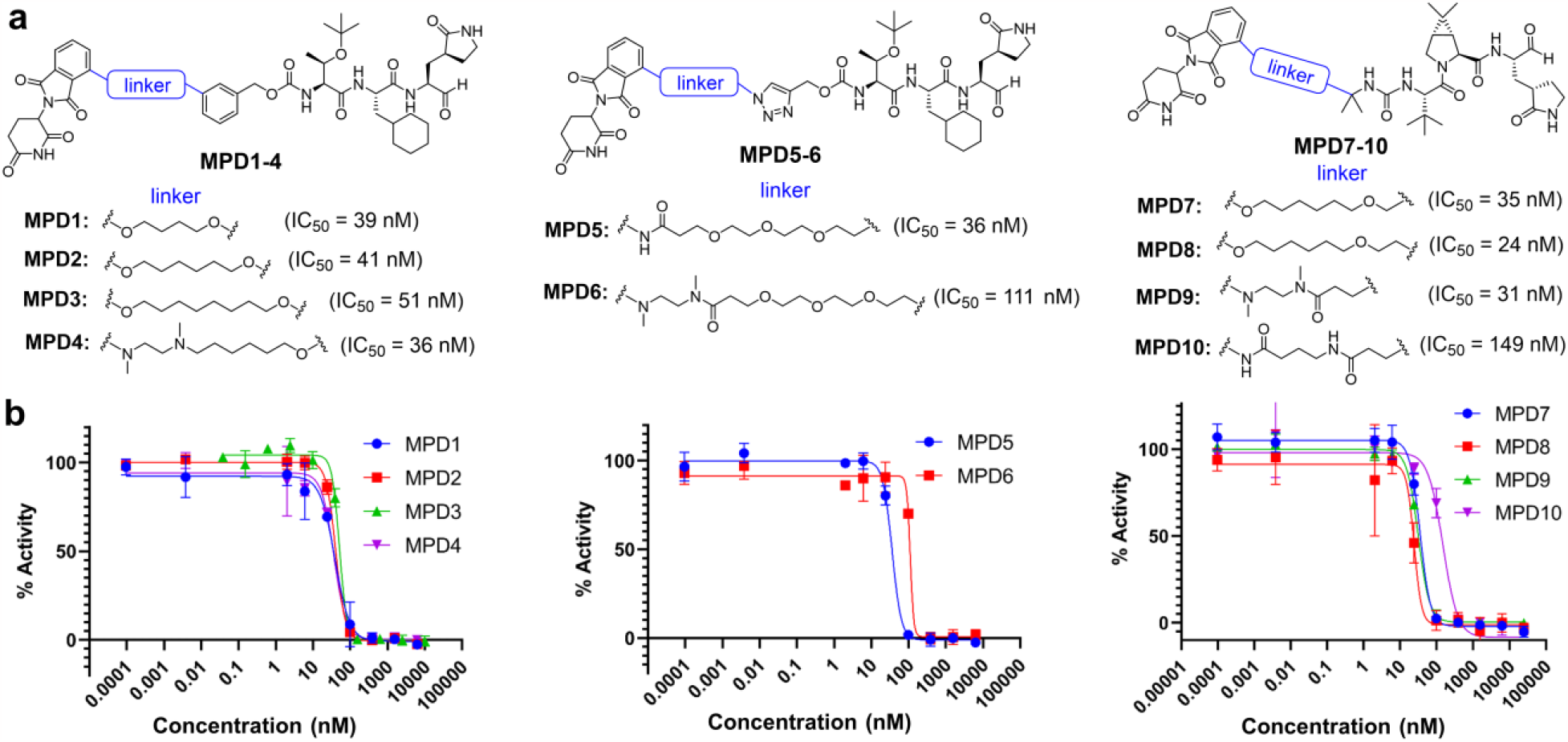
M^Pro^ PROTAC degraders. **a**, Chemical structures of M^Pro^ PROTAC degraders MPD1-MPD10. **b**, Inhibition curves of MPD1-MPD10 on M^Pro^. Triplicate experiments were performed for each compound. For all experiments, 20 nM M^Pro^ was incubated with an inhibitor for 30 min before 10 μM Sub3 was added. The M^Pro^-catalyzed Sub3 hydrolysis rate was determined by measuring linear increase of product fluorescence (Ex: 336 nm/Em: 490 nm) at the initial 5 min reaction time.

### MPD2 is a potent M^Pro^ degrader

To characterize the degradation of M^Pro^ by PROTAC degraders MPD1-MPD10, we established a cellular system with a stable expression of M^Pro^. For this purpose, we constructed a lentivirus-based system capable of expressing an M^Pro^-eGFP fusion protein (Figure 4a). This system allowed us to generate 293T cells that stably express M^Pro^-eGFP. M^Pro^ natively undergoes cleavage at its *N*- and *C*-terminus and requires a free *N*-terminal end for activation. To maintain the activity of M^Pro^, an *N*-terminal MAVLQ self-cleavage tag was introduced to the fusion protein. Additionally, to prevent the cleavage of the *C*-terminal eGFP, the C-terminal residue Q306 of M^Pro^ was mutated to Gly. Importantly, the transfection of 293T cells with pLVX-M^Pro^-eGFP shows much less toxicity after multiple rounds of proliferation than another construct we previously reported.^53^ This helps us to generate stable cell lines with high levels of eGFP fluorescence for evaluating M^Pro^ PROTACs. Our flow cytometry results confirmed the expression of the fusion protein, displaying strong fluorescence (Figure 4c). This was further corroborated with Western blot analysis to determine half-maximal degradation concentration (DC_50_) using an anti-M^Pro^ antibody (Figure 4b). Among these M^Pro^ PROTACs, MPD1, MPD2, and MPD3 demonstrated high potency in inducing M^Pro^ degradation in M^Pro^-eGFP stable cell lines. Their DC_50_ values were 419 nM, 296 nM, 431 nM, respectively (Figure 4c-h). Next, we evaluated the cytotoxicity of MPD1-MPD3 using 293T cells and the MTT assay.^61^ The determined CC_50_ values for MPD1, MPD2 and MPD3 were 25 μM, 120 μM, and 21 μM, respectively (Figure S3). The cytotoxicity curves for these three compounds are presented in Figure S3. Notably, MPD2 exhibited significantly lower toxicity even when compared to the parental MPI8 (CC_50_ = 70 μM). Consequently, MPD2 was selected for further evaluation.

**Figure 4.**
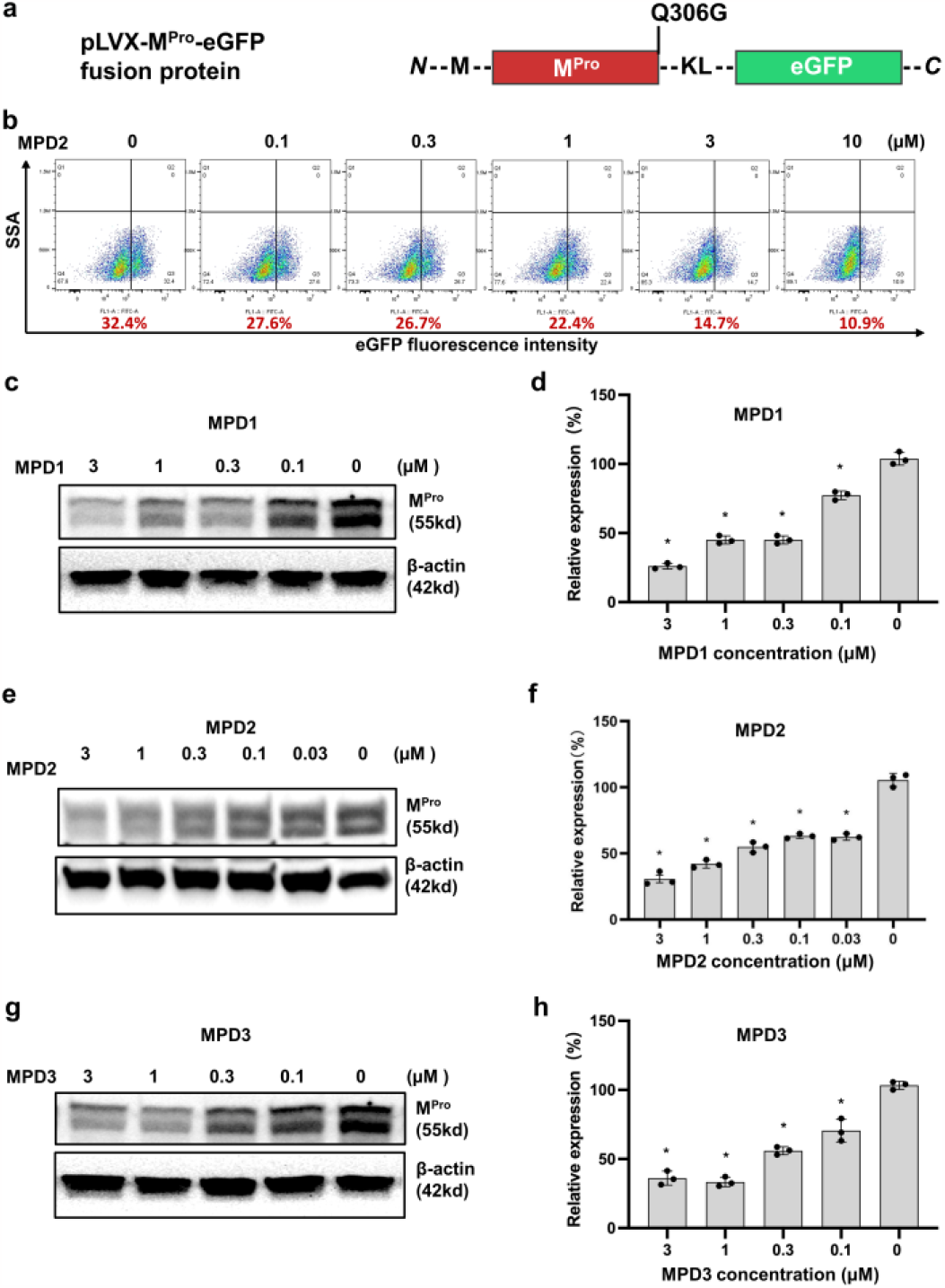
Degradation of M^Pro^ by MPD1-MPD3. **a**, Design of M^Pro^-eGFP fusion. **b**, Representative flow cytometry analysis of the potency of MPD2 in degrading M^Pro^ in M^Pro^-eGFP 293T stable cell line. M^Pro^-eGFP cells were evaluated after being treated with different concentrations of MPD2 for 48 h. The percentage of positively expressed M^Pro^-eGFP fusion protein was displayed in red at the bottom of the graph. **c**,**e**,**g**, The potency of MPD1 (**c**), MPD2 (**e**), and MPD1 (**g**) in degrading M^Pro^ was evaluated in M^Pro^-eGFP 293T stable cell line by immunoblots after the cells were treated with different concentrations of MPD1, MPD2, and MPD3 for 48 h. Representative immunoblots are shown, and β-actin was used as a loading control in all immunoblot analyses. **d**,**f**,**h**, The graph presents the normalized protein content in the immunoblots as mean values ± s.e.m. (n = 3) in the graph.

The induction of M^Pro^ degradation by MPD2 in M^Pro^-eGFP 293T stable cells was rapid and long-lasting. As illustrated in Figure 5a and 5b, significant degradation was already observed after 6 h of incubation with 3 μM MPD2, and nearly maximal degradation was achieved after 12 h of incubation. Furthermore, a one-time incubation of 3 μM MPD2 for 72 h yielded comparable results to the scenario where MPD2 incubation medium was changed every 24 h over the same 72 h period (Figure 5c). To evaluate the *in vitro* metabolic stability of MPD2, we conducted an analysis using human liver microsomes. The determined clearance (CL_int_) value for MPD2 was 36.2 mL/min/kg, resulting in a half-life (t_1/2_) of 48 minutes (Figure S4).

**Figure 5.**
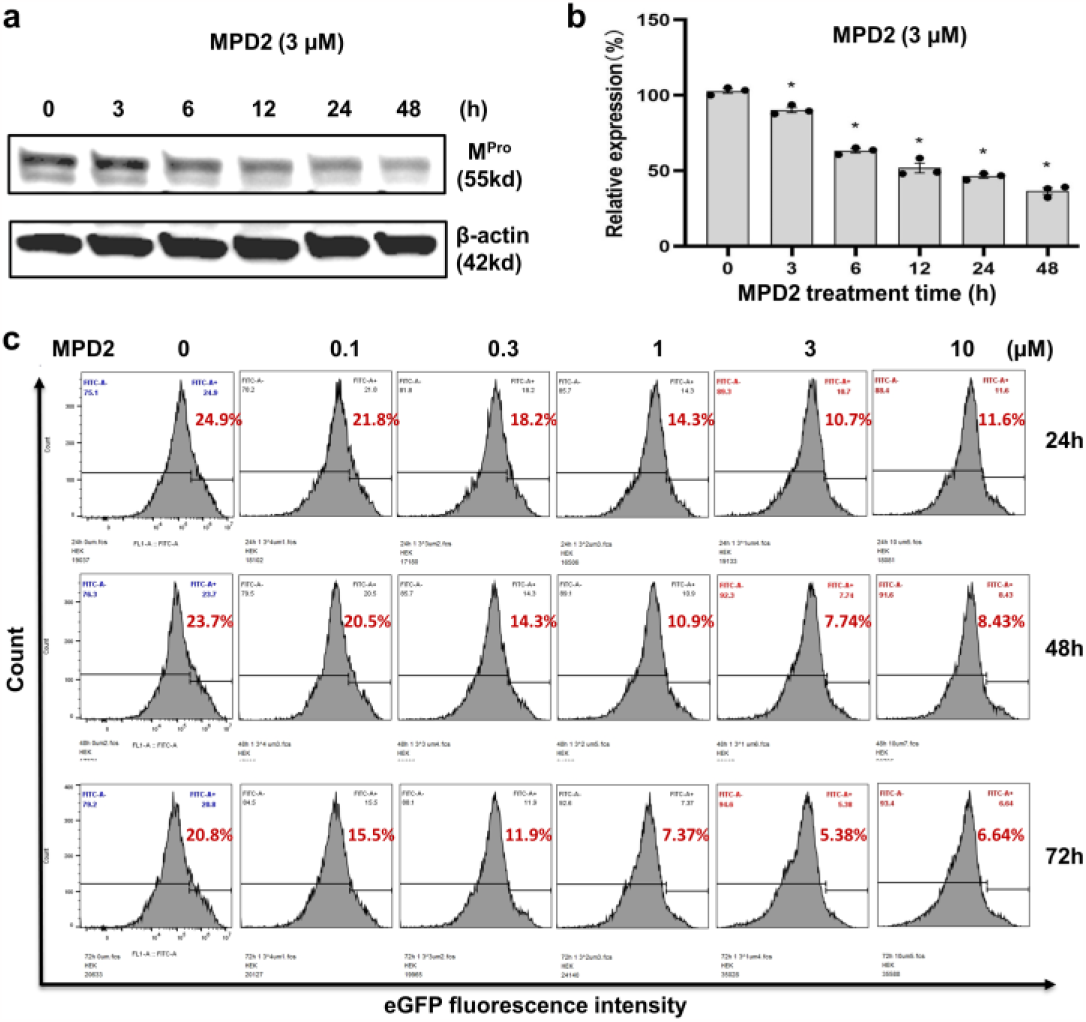
PROTAC degrader MPD2 downregulates the protein levels of M^Pro^ in a time-dependent manner. **a**, The time course of MPD2-mediated M^Pro^ degradation was evaluated in M^Pro^-eGFP 293T stable cell line by immunoblots after the cells were treated with 3 μM MPD2 for various time points as indicated. **b**, The graph presents the normalized protein content in the immunoblots as mean values ± s.e.m. (n = 3 biologically independent experiments) in the graph. **c**, Incubation of MPD2 with different concentrations for 72 h with changing MPD2 incubation medium every 24 h. The percentage of positively expressed M^Pro^-eGFP fusion protein was displayed on the upright top panel in every graph.

### Mechanism of M^Pro^ degradation by MPD2

To validate the mechanism underlying M^Pro^ degradation depending on the concurrent presence of both M^Pro^ and CRBN binding, we conducted competition assay experiments. Initially, the pretreatment of the cells with 3 μM MPI8 (an M^Pro^ ligand) completely blocked the degradation of M^Pro^ by MPD2 (Figure 6a,b and Figure S5). Similarly, the pretreatment of the cells with 3 μM Pomalidomide (a CRBN ligand) resulted in the suppression of the M^Pro^ degradation by MPD2 (Figure 6a,b and Figure S5). These studies implied that the degradation of M^Pro^ induced by MPD2 depends on M^Pro^ binding and CRBN binding. To further establish the CRBN dependance of the degradation mechanism of MPD2, we generated CRBN knockout (CRBN-KO) M^Pro^-eGFP 293T cells using CRISPR editing technique. When comparing the MPD2-induced degradation of M^Pro^ in wildtype CRBN (CN) and CRBN (KO) cells, we observed a significant rescue of intracellular M^Pro^ in cells lacking CRBN (Figure 6c,d and Figure S5). The results of the competition assays with MPI8 and pomalidomide, alongside the CRBN knockout study, collectively affirm that MPD2-induced M^Pro^ degradation relies on both M^Pro^ and CRBN engagement, elucidating a CRBN-mediated mode of action.

**Figure 6.**
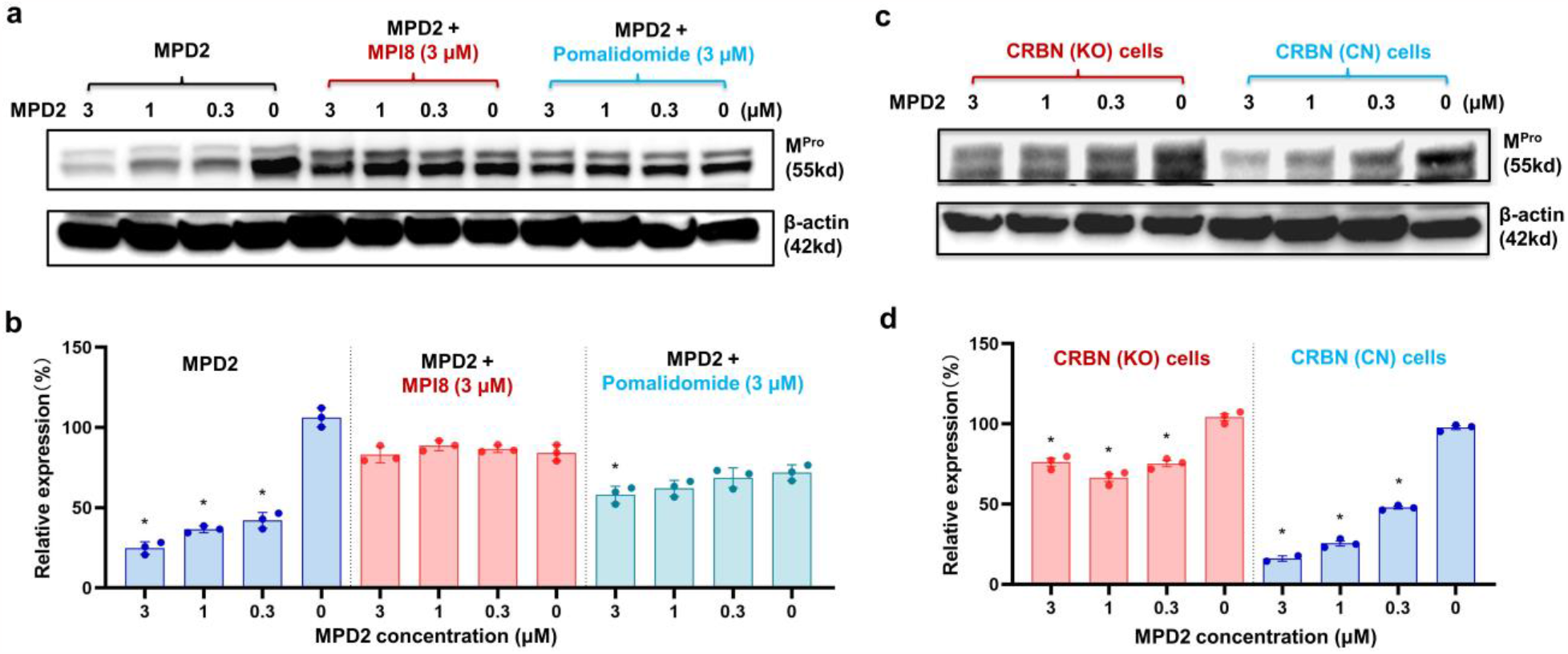
MPD2 degrades M^Pro^ in a ligand- and CRBN-dependent manner. **a**, Pretreatment with M^Pro^ ligand MPI8 or CRBN ligand Pomalidomide blocks the M^Pro^ degradation induced by MPD2. The potency of MPD2 in degrading M^Pro^ was evaluated in M^Pro^-eGFP 293T stable cell line by immunoblots after the cells were treated with different concentrations of MPD2 for 48 h. **b**, Representative immunoblots are shown and β-actin was used as a loading control in all immunoblot analyses. The normalized protein content in the immunoblots is presented as mean values ± s.e.m. (n = 3 biologically independent experiments) in the bar graph. **c**, CRISPR knockout of CRBN blocks MPD2-induced M^Pro^ degradation as shown in control (CRBN-CN) and CRBN knockout (CRBN-KO) M^Pro^-eGFP 293T cells. Immunoblots evaluated the potency of MPD2 in degrading M^Pro^ after the cells were treated with different concentrations of MPD2 for 48 h. **d**, Representative immunoblots are shown and β-actin was used as a loading control in all immunoblot analyses. The normalized protein content in the immunoblots is presented as mean values ± s.e.m. (n = 3 biologically independent experiments) in the bar graph.

Moreover, we observed that the proteasome inhibitor MG132 (1 μM) effectively rescued MPD2-induced M^Pro^ degradation. The notable increase in M^Pro^ levels resulting from the cotreatment of cells with MPD2 and MG132 unequivocally validates the indispensability of proteasome activity in the mechanistic degradation of MPD2 (Figure 7). It is essential to note that the presence of MG132 inhibits proteasome activity, thereby impeding the natural ubiquitination process of M^Pro^, which subsequently leads to elevated M^Pro^ levels. Collectively, our studies utilizing CRBN knockout cells, in combination with MPI8, pomalidomide, MG132 and MPD2 treatments, provide compelling evidence that the degradation activity of MPD2 relies on the engagement of M^Pro^, CRBN, and proteasome-mediated degradation. These findings underscore the multifaceted nature of M^Pro^ degradation induced by MPD2, requiring M^Pro^ binding, CRBN binding, and proteasome-mediated processes.

**Figure 7.**
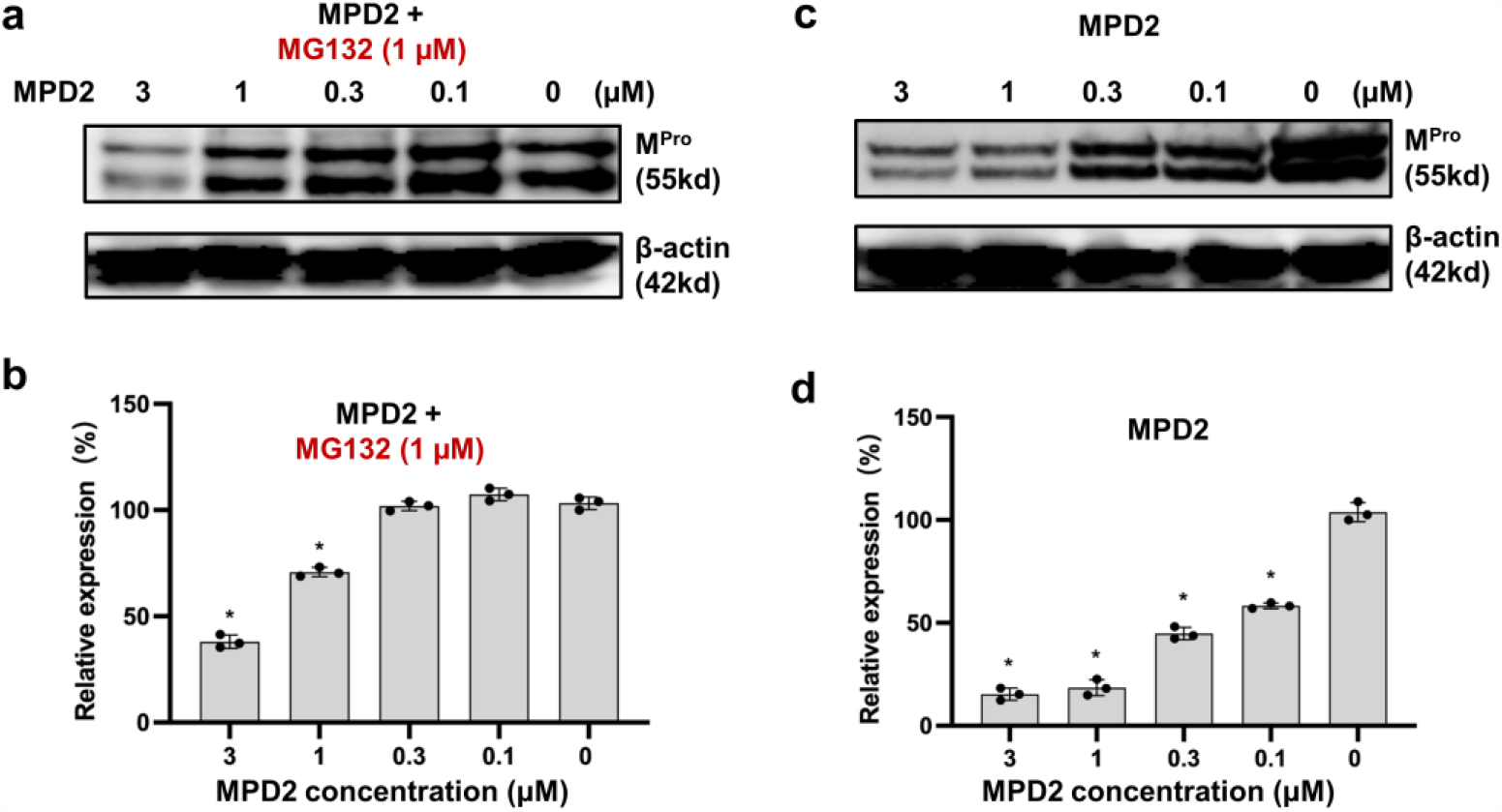
MPD2 degrades M^Pro^ in a proteasome-dependent manner and proteasome inhibition blocks the M^Pro^ degradation by MPD2. **a**,**c**, A representative of 3 immunoblot analyses of M^Pro^ in M^Pro^-eGFP 293T stable cell line after they were either pretreated with the proteasome inhibitor MG132 (1 μM) or pretreated with vehicle for 1 h, and then were treated with different concentration of MPD2 for 48 h. **b**,**d**, Representative immunoblots are shown and β-actin was used as a loading control in all immunoblot analyses. The normalized protein content in the immunoblots is presented as mean values ± s.e.m. (n = 3 biologically independent experiments) in the bar graph.

### Antiviral Potency Characterizations

After successfully validating the degradation of M^Pro^ and understanding the mechanism of degradation induced by the PROTAC degraders, we proceeded to evaluate their antiviral efficacy against SARS-CoV-2 in ACE2^+^ A549 cells infected with hCoV-19/USA/HP05647/2021, an early Delta variant known for its robust growth in cell culture and strong cytopathic effects. MPD1-3 were administered at varying concentrations, and the cells were cultured for 72 h before quantifying SARS-CoV-2 mRNA levels using RT-PCR to determine the antiviral EC_50_ values. MPD1, MPD2, and MPD3 exhibited antiviral EC_50_ values of 1780 nM, 492 nM, and 1160 nM, respectively (Figure 8a). Furthermore, we investigated the impact of PROTAC degrader treatment on M^Pro^ accumulation during SARS-CoV-2 infection. A549-ACE2 cells were inoculated at an approximate multiplicity of 0.1 and subsequently treated with MPD1-3, starting 1 h after inoculation. After 48 h post-inoculation, cell lysates were prepared and subjected to blotting using a monoclonal antibody against the 34 kDa M^Pro^. All three degraders MPD1-3 demonstrated the ability to induce M^Pro^ degradation. Among them, MPD2 emerged as the most potent degrader, which correlates with its highest antiviral activity (Figure 8b).

**Figure 8.**
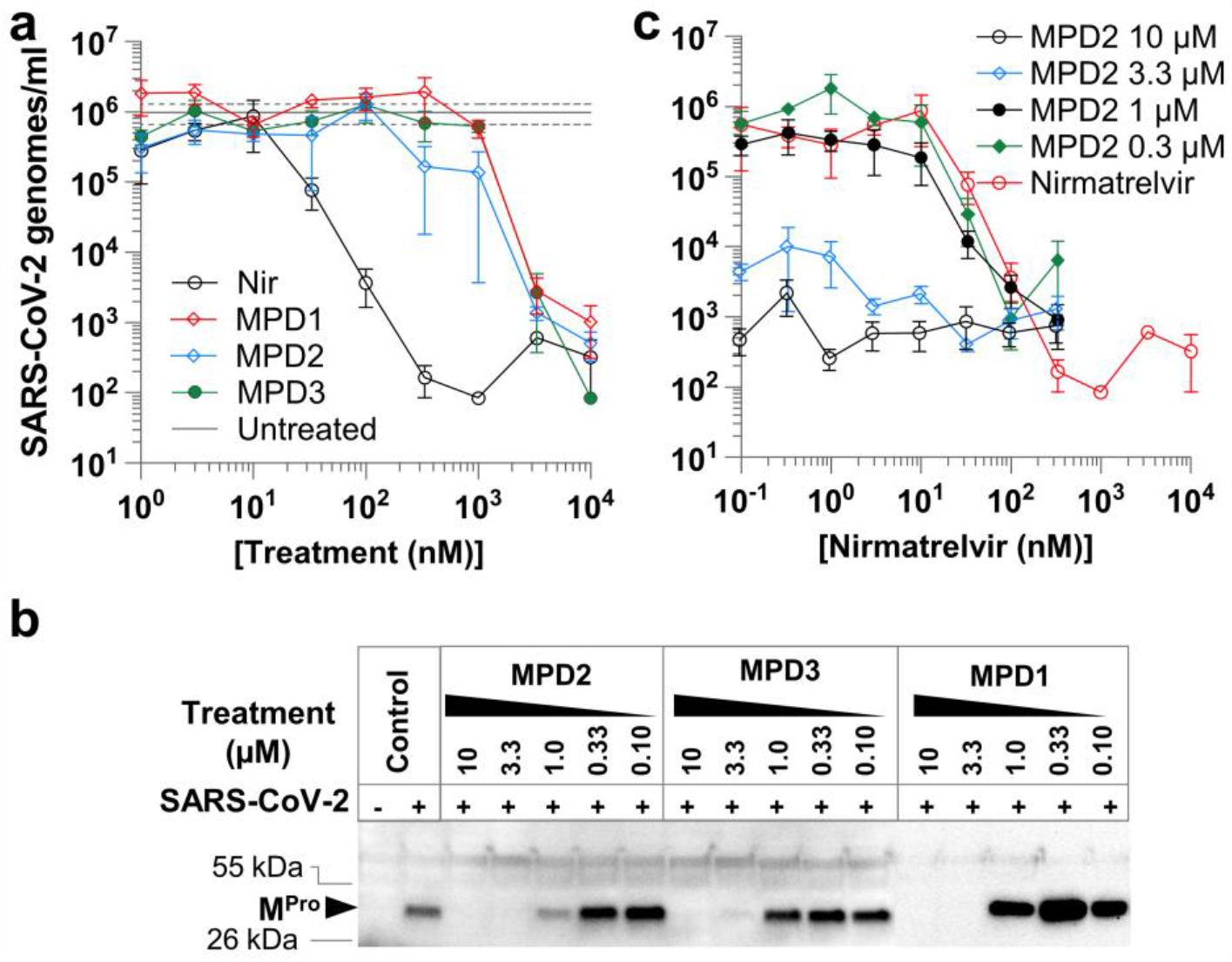
Antiviral effectiveness of PROTACs. A549-ACE2 cells were inoculated with 0.01 TCID_50_ per cell (A, C), or 0.1 TCID_50_ per cell (B) SARS-CoV-2 for 1 h, then treated with antiviral. (A) Effect of PROTACs or nirmatrelvir (Nir) treatment on virus growth 48 h after inoculation. (B) Effect of PROTACs treatment on M^Pro^ accumulation 48 h after inoculation with SARS-CoV-2. M^Pro^ in cell lysates was detected with monoclonal antibody against the 34 kDa M^Pro^. (C) Antiviral synergy testing with fixed concentrations of PROTAC MPD2 and variable concentrations of nirmatrelvir; virus growth was measured 48 h after inoculation. Error bars represent standard error.

We observed synergistic effects between the PROTAC degrader MPD2 and nirmaltrevir (the active antiviral component of the COVID-19 drug Paxlovid) during SARS-CoV-2 infection (Figure 8c). ACE2^+^ A549 cells infected with SARS-CoV-2 were treated with fixed concentrations of PROTAC MPD2 in combination with varying concentrations of nirmaltrevir. Notably, the presence of 1 μM MPD2 shifted the dose-response curve of nirmaltrevir to the left in a dose-dependent manner (Figure 8c). These findings underscore the potential synergistic benefits of combining an M^Pro^ inhibitor and an M^Pro^ degrader, enhancing the overall antiviral efficacy against SARS-CoV-2.

## DISCUSSION

PROTAC is a new technology for targeted therapy in drug discovery. PROTACs are event-driven bifunctional small molecules that simultaneously engage an E3 ubiquitin ligase and a target protein to facilitate the formation of a ternary complex, leading to the ubiquitination and ultimate degradation of the target protein. The distinct mechanism of action offers several compelling advantages over traditional inhibitors, including: (i) catalytic nature to allow for substoichiometric activity, (ii) recruiting target via any binding site where functional and sustained inhibition is not required; (iii) enhanced target selectivity controlled by protein–protein interactions, (iv) high barrier to resistance; and (iv) abrogating all functions of the target protein and its downstream proteins. These inherent advantages have propelled PROTAC technology into rapid development and wide-ranging applications across various diseases. Numerous smallmolecule PROTACs have advanced into clinical trials, particularly in the realm of targeted cancer therapy. While the utilization of PROTAC technology in antiviral research remains relatively limited, and the pool of developed antiviral PROTAC degraders is small, the concept of PROTAC-induced targeted protein degradation is gaining traction as a novel strategy for the development of next-generation antiviral drugs to combat infectious diseases caused by various viruses.^45-48^

In this study, we worked on the design, synthesis, and evaluation of bifunctional M^Pro^ PROTAC degraders MPD1-MPD10 based on the M^Pro^ ligands MPI8 and MPI29 we previously developed. The CRBN-binding moieties of MPD1-MPD10 were derived from the well-established ligands thalidomide-4-OH and pomalidomide. Notably, all MPD1-MPD10 compounds exhibited sub-micromolar IC_50_ values, underscoring the absence of any discernible interference from the linkers and CRBN E3 ligands with M^Pro^ binding (Figure 3). The M^Pro^ degradation by PROTAC degraders MPD1-MPD3 was determined by a cellular system utilizing the M^Pro^-eGFP fusion protein and Western blot analysis (Figure 4). Our studies revealed that MPD2 effectively facilitated M^Pro^ degradation in M^Pro^-eGFP 293T stable cells, characterized by rapid and enduring effects. To substantiate the degradation mechanism of the M^Pro^ degrader MPD2, we conducted a series of control experiments. Notably, pretreatment with an M^Pro^ inhibitor MPI8 (as a competitor for M^Pro^ binding), or a CRBN E3 ligand Pomalidomide (as a competitor of CRBN ligase binding), led to the suppression of M^Pro^ degradation induced by MPD2. These findings strongly implied that MPD2-induced M^Pro^ degradation hinges on both M^Pro^ and CRBN binding. Furthermore, studies involving CRBN knockout cells demonstrated a significant reduction in the degradation potency of MPD2, further affirming a CRBN-dependent mechanism (Figure 6). Our investigation also unveiled that the proteasome inhibitor MG132 could rescue the degradation triggered by MPD2, underscoring the dependence of MPD2’s activity on proteasome-mediated degradation (Figure 7). Taken together, MPD2 has proven to be a potent and effective M^Pro^ degrader that relies on a multifaceted mechanism encompassing M^Pro^ binding, CRBN binding, and proteasome-mediated degradation. Importantly, MPD2 exhibited remarkable antiviral activity against SARS-CoV-2 with an EC_50_ value of 492 nM, and successfully degraded M^Pro^ in SARS-CoV-2-infected A549-ACE2 cells (Figure 8). Notably, our findings revealed an intriguing synergy between the M^Pro^ degrader MPD2 and nirmatrelvir, enhancing the overall antiviral efficacy against SARS-CoV-2. The presence of 1 μM MPD2 enhanced the dose-response curve of nirmaltrevir, causing a leftward shift in a dose-dependent fashion. (Figure 8c).

In summary, we have developed the first series of SARS-CoV-2 M^Pro^ degraders, with MPD2 emerging as a standout lead compound. MPD2 effectively reduced M^Pro^ protein levels in 293T cells, relying on a time-dependent, CRBN-mediated, and proteasome-driven mechanism. Furthermore, MPD2 exhibited remarkable efficacy in diminishing M^Pro^ protein levels in SARS-CoV-2-infected A549-ACE2 cells, concurrently demonstrating potent anti-SARS-CoV-2 activity, with a promising EC_50_ value of 492 nM. This proof-of-concept study highlights the potential of leveraging PROTAC-mediated targeted protein degradation as a novel approach to target M^Pro^ for COVID-19 drug discovery. While M^Pro^ is generally considered to be more conserved across CoVs compared to the Spike protein, the emergence of drug-resistant variants carrying mutations in M^Pro^ has become increasingly evident with the widespread use of nirmatrelvir/ritonavir. Therefore, the development of antiviral PROTAC degraders utilizing M^Pro^ ligands, which target more conserved and non-catalytic binding pockets, would offer a promising strategy to address the pressing issue of drug resistance in the treatment of COVID-19.

### EXPERIMENTAL SECTION

#### Materials

We purchased yeast extract from Thermo Fisher Scientific, tryptone from Gibco, Sub3 from Bachem, HEK 293T/17 cells from ATCC, DMEM with GlutaMax from Gibco, FBS from Gibco, polyethyleneimine from Polysciences, and the trypsin-EDTA solution from Gibco. Chemicals used in this work were acquired from Sigma Aldrich, Chem Impex, Ambeed, and A2B. Pooled human liver microsome (1910096) was obtained from Xenotech.

#### M^Pro^ Expression and Purification

The expression plasmid pET28a-His-SUMO-M^Pro^ was constructed in a previous study. We used this construct to transform *E. coli* BL21(DE3) cells. A single colony grown on an LB plate containing 50 μg/mL kanamycin was picked and grown in 5 mL LB media supplemented with 50 μg/mL kanamycin overnight. We inoculated this overnight culture to 6 L 2YT media with 50 μg/mL kanamycin. Cells were grown to OD_600_ as 0.8. At this point, we added 1 mM IPTG to induce the expression of His-SUMO-M^Pro^. Induced cells were let grown for 3 h and then harvested by centrifugation at 12,000 rpm, 4 °C for 30 min. We resuspended cell pellets in 150 mL lysis buffer (20 mM Tris-HCl, 100 mM NaCl, 10 mM imidazole, pH 8.0) and lysed the cells by sonication on ice. We clarified the lysate by centrifugation at 16,000 rpm, 4 °C for 30 min. We decanted the supernatant and mixed with Ni-NTA resins (GenScript). We loaded the resins to a column, washed the resins with 10 volumes of lysis buffer, and eluted the bound protein using elution buffer (20 mM Tris-HCl, 100 mM NaCl, 250 mM imidazole, pH 8.0). We exchanged buffer of the elute to another buffer (20 mM Tris-HCl, 100 mM NaCl, 10 mM imidazole, 1 mM DTT, pH 8.0) using a HiPrep 26/10 desalting column (Cytiva) and digested the elute using 10 units SUMO protease overnight at 4 °C. The digested elute was subjected to Ni-NTA resins in a column to remove His-tagged SUMO protease, His-tagged SUMO tag, and undigested His-SUMO-M^Pro^. We loaded the flow-through onto a Q-Sepharose column and purified M^Pro^ using FPLC by running a linear gradient from 0 to 500 mM NaCl in a buffer (20 mM Tris-HCl, 1 mM DTT, pH 8.0). Fractions eluted from the Q-Sepharose column was concentrated and loaded onto a HiPrep 16/60 Sephacryl S-100 HR column and purified using a buffer containing 20 mM Tris-HCl, 100 mM NaCl, 1 mM DTT, and 1 mM EDTA at pH 7.8. The final purified was concentrated and stored in a -80 °C freezer.

#### *In Vitro* M^Pro^ Inhibition Potency Characterizations

We conducted the assay using 20 nM M^Pro^ and 10 μM Sub3 (Figure S2). We dissolved all compounds in DMSO as 10 mM stock solutions. Sub3 was dissolved in DMSO as a 1 mM stock solution and diluted 100 times in the final assay buffer containing 10 mM Na_x_H_y_PO_4_, 10 mM NaCl, 0.5 mM EDTA, and 1.25% DMSO at pH 7.6. We incubated M^Pro^ and an inhibitor in the final assay buffer for 30 min before adding the substrate to initiate the reaction catalyzed by M^Pro^. The production format was monitored in a fluorescence plate reader with excitation at 336 nm and emission at 490 nm.

#### Establishment of 293T Cells Stably Expressing M^Pro^-eGFP

To establish a 293T cell line that stably expresses M^Pro^-eGFP, we packaged lentivirus particles using the pLVX-M^Pro^-eGFP plasmid, detailed preparation was shown in a previous publication53. Briefly, we transfected 293T cells at 90% confluency with three plasmids including pLVX-M^Pro^-eGFP, pMD2.G, and psPAX2 using 30 mg/mL polyethylenimine. We collected supernatants at 48 and 72 h after transfection separately. We concentrated and collected lentiviral particles from collected supernatant using ultracentrifugation. We then transduced fresh 293T cells using the collected lentivirus particles. After 48 h of transduction, we added puromycin to the culture media to a final concentration of 2 μg/mL. We gradually raised the puromycin concentration 10 μg/mL in 2 weeks. The final stable cells were maintained in media containing 10 μg/mL puromycin.

#### Western blot and DC_50_ (half-maximal degradation concentration) analysis

For protein extraction, 5×105 cells per well were plated onto twelve-well plates and treated with a PROTAC molecule at indicated doses and time duration. We did protein extraction and Western blot analysis. In a solution containing 50 mM Tris (pH 7.5), 150 mM NaCl, 5 g/ml aprotinin, 1 g/ml pepstatin, 1% Nonidet P-40, 1 mM EDTA, and 0.25% deoxycholate, cells were lysed. In a cold room, protein lysates were sonicated for two minutes before being centrifuged at maximum speed (14,000 rpm) for 15 minutes. Using the Bradford Reagent (cat. no. 97065-020, VWR), the protein content in the supernatants was measured. After normalizing the protein concentration, the samples were reduced in 4×Laemmli’s SDS-sample buffer and desaturated at 95 °C for 10 minutes. Using 10% SDS–PAGE, an equal quantity of protein samples (20 μg each lane) were separated. Subsequent protein signals were produced using Pierce ECL Western blotting substrate (Thermo Scientific; 32106) and visualized using the ChemiDoc MP Imaging System (Bio-Rad). Data were represented as relative band intensities adjusted to equal loading control. Antibody against SARS-CoV-2 M^Pro^ was acquired from Thermo Fisher Scientific (cat. no. PA5-116940). β-actin antibody was purchased from ABCAM (Cat. No. ab124964). Goat Anti-Rabbit IgG H&L (HRP) was acquired from ABCAM as the secondary antibody (Cat. No. ab6721).

Dose-response curves for the tested drugs in M^Pro^-eGFP 293T stable cells. Various dilutions of the drugs were applied to the 80% confluent cell monolayers and assayed after 48 h to determine the DC_50_ value. β-actin was used as a loading control in all immunoblot analyses. The normalized protein content in the immunoblots is presented as mean values ± s.e.m. (n = 3 biologically independent experiments). Nonlinear regression analysis of GraphPad Prism software (version 8.0) was used to calculate DC_50_ by plotting log compounds concentration versus normalized response (variable compounds).

#### CRBN knockout by CRISPR/Cas9 genomic editing

To deplete CRBN, CRISPR/Cas9 KO Plasmids, consisting of CRBN-specific 20 nt guide RNA sequences derived from the GeCKO (v2) library, were purchased from Santa Cruz (sc-425519). Briefly, In a 6-well tissue culture plate seed 1.5 x 105 - 2.5 x 105 cells in 3 ml of antibiotic-free standard growth medium per well, 24 hours prior to transfection. Grow cells to a 40–80% confluency. Initial cell seeding and cell confluency after 24 hours are determined based on the rate of cell growth of the cells used for transfection. Healthy and subconfluent cells are required for successful knockout.

#### Live virus antiviral testing

SARS-CoV-2 delta variant hCoV-19/USA/MD-HP05647/2021 (BEI Resources, NR-55672) was propagated in A549-hACE2 cells (BEI Resources NR-53522) for antiviral testing, at 37ºC in an air-jacketed incubator, with 5% CO_2_ and >90% relative humidity, under BSL-3 conditions at the Texas A&M Global Health Research Complex. A low-dose, multi-step growth protocol was used for live virus EC_50_ assays. Briefly, A549-hACE2 cells were cultured in DMEM supplemented with 10% fetal bovine serum overnight. Approximately 5×10^4^ A549-hACE2 cells were inoculated by adding 10^3^ infectious units of SARS-CoV-2, as determined by tissue culture infectious dose 50% (TCID50) assay, and incubated at 37ºC for 1 hour. Cells were then aspirated and rinsed three times with room temperature phosphate buffered saline, to remove residual inoculum, before replacing DMEM with 10% FBS. Serial three-fold dilutions of candidate antivirals were made in DMEM with 10% FBS and added to three replicate wells per treatment condition. Infected, treated cells were then cultured for 72h at 37ºC, 5% CO_2_. At 48h, and 72h after inoculation, 50 microliters of tissue culture medium were removed from each sample, for quantitative SARS-CoV-2 reverse transcriptase quantitative PCR (RT-qPCR). After 72h, cell culture supernatant was removed by aspiration, cells were fixed in 10% formalin, 1× phosphate buffered saline for at least 30 min, then stained with crystal violet in order to qualitatively assess cytopathic effect. EC_50_ values were calculated from the slope and intercept of log-transformed RT-qPCR results, at the point where the linear portion of the transformed dose-response curve showed 50% reduced growth compared to infected, untreated controls. Samples were processed and RT-qPCR was performed as per protocol established previously.^62^ Samples were diluted 1:1 in 2× Tris-borate-EDTA [TBE] containing 1% Tween-20 and heated at 95°C for 15 min to lyse and inactivate virions. RT-qPCR screening was performed using the CDC N1 oligonucleotide pair/FAM probe (CDC N1-F, 5′-GACCCCAAAATCAGCGAAAT-3′; CDC N1-R, 5′-TCTGGTTACTGCCAGTTGAATCTG-3′; and Probe CDC N1, 5′ FAM-ACCCCGCATTACGTTTGGTGGACC-BHQ1 3′) and the Luna Universal Probe one-step RT-qPCR kit (catalog no. E3006; New England Biolabs). A 20-μL RT-qPCR mixture contained 7 μL of sample, 0.8 μL each of forward and reverse oligonucleotides (10 μM), 0.4 μL of probe (10 μM), and 11 μL of NEB Luna one-step RT-qPCR 2× master mix. Samples were incubated at 55°C for 10 min for cDNA synthesis, followed by 95°C for 1 min (1 cycle), then 41 cycles of 95°C for 10 s and 60°C for 30 s. Genome copies were quantitated relative to quantitative PCR control RNA from heat-inactivated SARS-Related Coronavirus 2, Isolate USA-WA1/2020 (BEI Resources, NR-52347.

#### Cytotoxicity Assay of the PROTAC degraders

To assess the half-maximal cytotoxic concentration (CC_50_), stock solutions of the tested PROTAC compounds were dissolved in DMSO (10mM) and diluted further to the working solutions with DMEM. HEK293T cells were seeded in 96 well-plates and incubated at 37°C and 5% CO_2_ for 24h. The cells were then treated with different concentrations (200 mM, 100 mM, 50 mM, 25 mM, 12.5 mM, 6.25 mM, 3.125 mM, 1.5625 mM, 0.78125 mM, 0 mM) of the tested compounds in triplicates for 48h. Cell viability was assessed by MTT assay to determine CC_50_. 20 mL MTT (5mg/mL) was incubated per well for 4h and then after removing supernatant, 200 mL DMSO was added per well. The absorbance was recorded at 490 nm to determine the CC_50_. The CC_50_ values were obtained by plotting the normalization % cell viability versus log_10_ sample concentration.

#### *In Vitro* Metabolic Stability in Human Liver Microsomes

This metabolic stability measurements were based on previous publications and modified as described below.^63^ The metabolic stability profile of the inhibitor, including CL_int, pred_ and *in vitro* t_1/2_ was determined by the estimation of the remaining compound concentration after incubation with human liver microsome, NADPH (cofactor), EDTA, and MgCl_2_ in a 0.1 M phosphate buffer (pH 7.4). 5 μM of each inhibitor was preincubated with 40 μL of human liver microsome (0.5 mg/mL) in 0.1 M phosphate buffer (pH 7.4) at 37°C for 5 min to set optimal a condition for metabolic reactions. After pre incubation, NADPH (5 mM, 10 μL) or 0.1 M PB (10 μL) was added to initiate metabolic reaction at 37°C. The reactions were conducted in triplicate. At 0, 5, 15, 30, 45, 60 min, 200 μL acetonitrile (with internal standard Diclofenac, 10 μg/mL) was added in order to quench the reaction. The samples were subjected to centrifugation at 4°C for 20 min at 4000 rpm. Then 50 μL of clear supernatants were analyzed by HPLC-MS/MS. The percentage of test compound remaining was determined by following formula: % remaining = (Area at t_x_ /Average area at t_0_) × 100. The half-life (t_1/2_) was calculated using the slope (k) of the log-linear regression from the % remaining parent compound versus time (min) relationship: t_½_ (min) = –ln 2/k. CL_int, pred_ (mL/min/kg) was calculated through following formula CLi_nt, pred_ = (0.693/t_1/2_) × (1/ (microsomal protein concentration (0.5 mg/mL)) × Scaling Factor (1254.16 for human liver microsome).

#### The Synthesis of MPDs

All reagents and solvents for the synthesis were purchased from commercial sources and used without purification. All glassware was flame-dried prior to use. Thin-layer chromatography (TLC) was carried out on aluminum plates coated with 60 F254 silica gel. TLC plates were visualized under UV light (254 or 365 nm) or stained with 5% phosphomolybdic acid. Normal phase column chromatography was carried out using a Yamazen Small Flash AKROS system. NMR spectra were recorded on a Bruker AVANCE Neo 400 MHz spectrometer in specified deuterated solvents. Analytical liquid chromatography-mass spectrometry was performed on a PHENOMENEX C18 Column (150 x 2.00 mm 5u micron, gradient from 10% to 100% B [A = 10 mmol/L HCOONH4 /H2O, B = MeOH], flow rate: 0.3 mL/min) using a Thermo Scientific Ultimate 3000 with a UV-detector (detection at 215 nm), equipped with a Thermo Scientific Orbitrap Q Exactive Focus System.

## ACKNOWLEDGMENT

This work was supported by the National Institutes of Health (grants R21AI164088 to S.X., and R35GM145351 and R21EB032983 to W.R.L.), the Welch Foundation (grant A-1715), and the Texas A&M X Grants.

## Notes

### Competing Interest Statement

The authors have declared no competing interest.

