## Supplementary Material for "Discovery of First-in-Class PROTAC Degraders of SARS-CoV-2 Main Protease"

Supplementary Figures

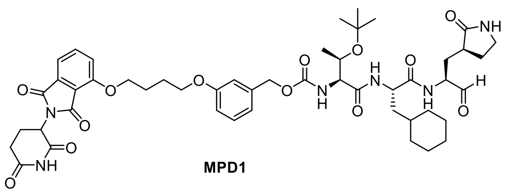

MPD1

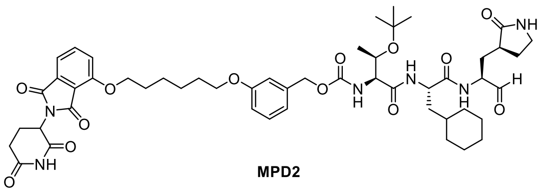

MPD2

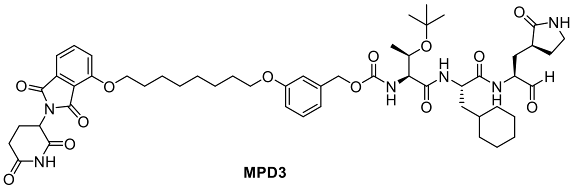

MPD3

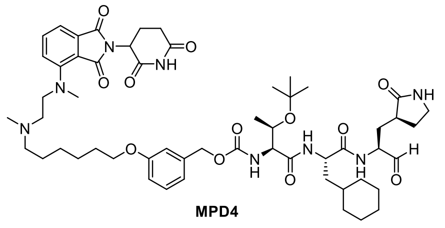

MPD4

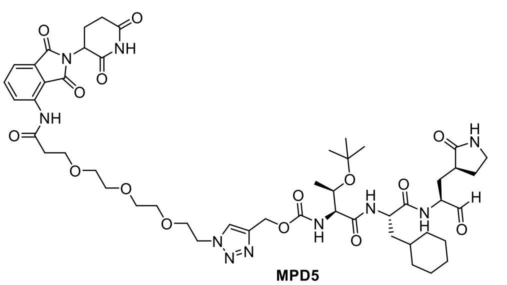

MPD5

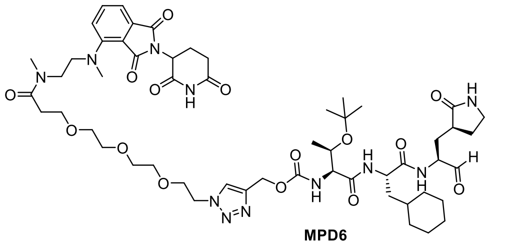

MPD6

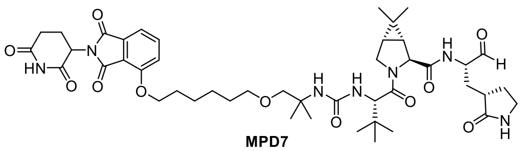

MPD7

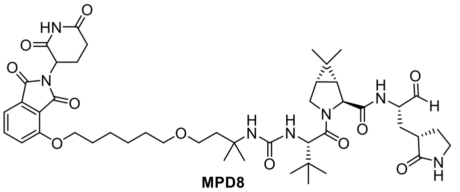

MPD8

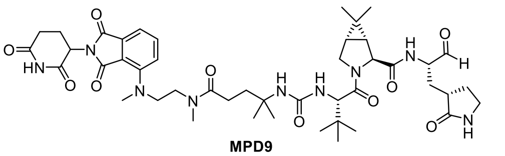

MPD9

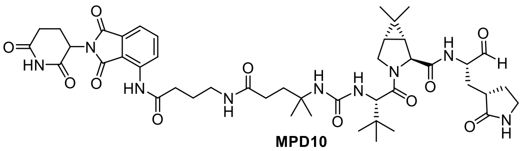

MPD10

**Figure S1**. Structures of MPD1-10.

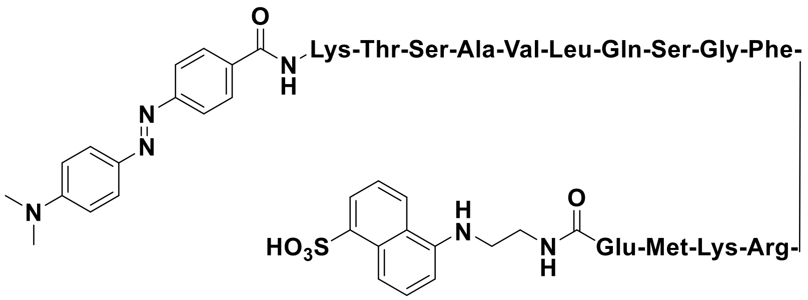


**Figure S2**. Chemical structure of Sub3.

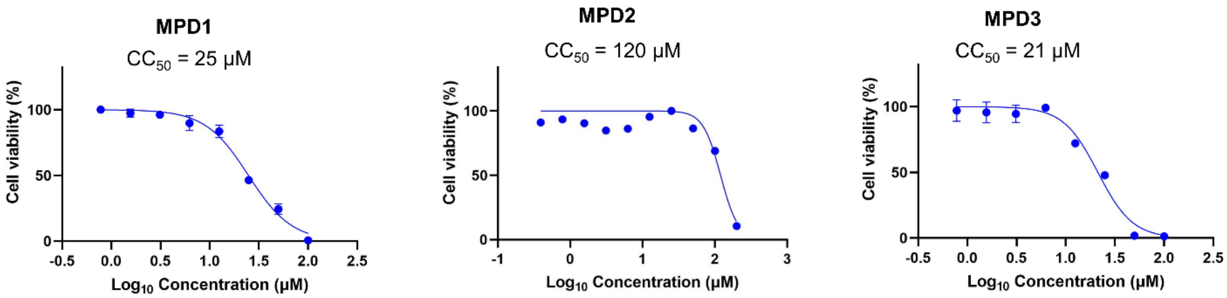


**Figure S3.** Cytotoxicity of the MPD1-MPD3 in 293T cells using the MTT assay. Dose-response curves for the tested drugs in 293T cells. Various dilutions of the drugs were applied to the 80% confluent cell monolayers and assayed after 48 h to determine the $CC_{50}$ (half-maximal cytotoxic concentrations). Nonlinear regression analysis of GraphPad Prism software (version 8.0) was used to calculate $CC_{50}$ by plotting log compounds versus normalized response (variable compounds).

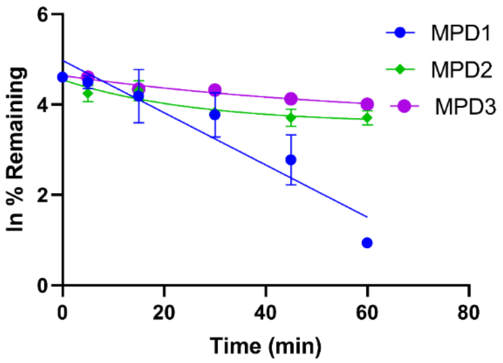


| Sample ID | Half life (min) | Intrinsic clearance (mL/min/kg) |
|---|---|---|
| MPD1 | 11.7 ± 1.04 | 149.4 ± 12.7 |
| MPD2 | 48.05 ± 3.22 | 36.28 ± 2.51 |
| MPD3 | 70.2 ± 1.5 | 24.7 ± 0.5 |

**Figure S4.** Stability of MPD1-MPD3 in human liver microsomes.

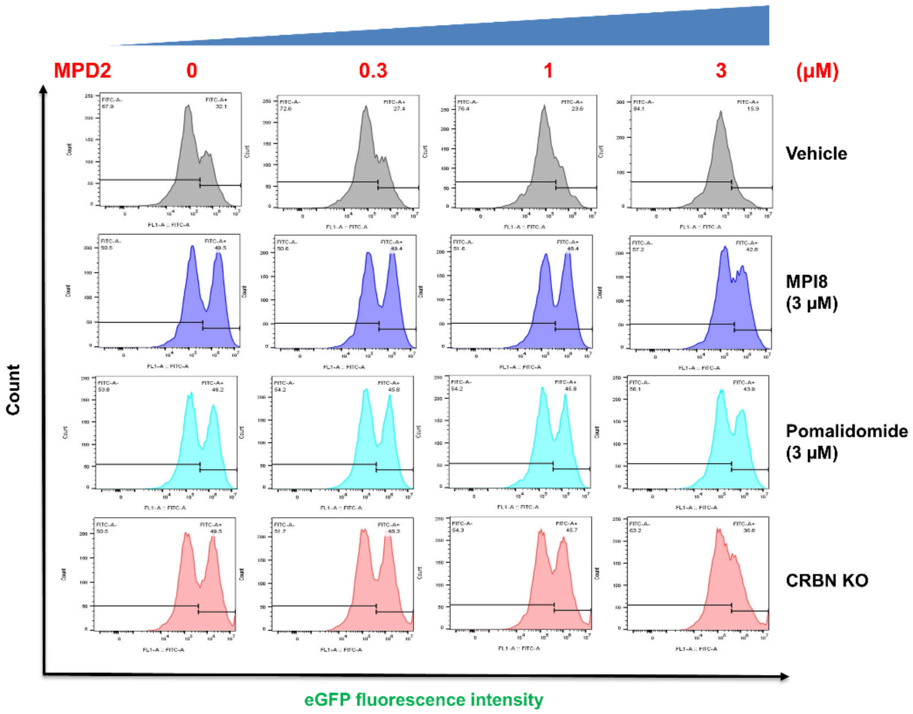


**Figure S5.** Flow Cytometry analysis of the potency of MPD2 in degrading $M^{Pro}$ in pretreatments of vehicle, MPI8, CRBN ligand (Pomalidomide) in $M^{Pro}$-eGFP 293T stable cell line or CRISPR knockout of CRBN in $M^{Pro}$-eGFP 293T stable cell line. The percentage of positively expressed $M^{Pro}$-eGFP fusion protein was displayed on the upright top panel in every histogram.

#### Synthesis of MPD1-MPD10

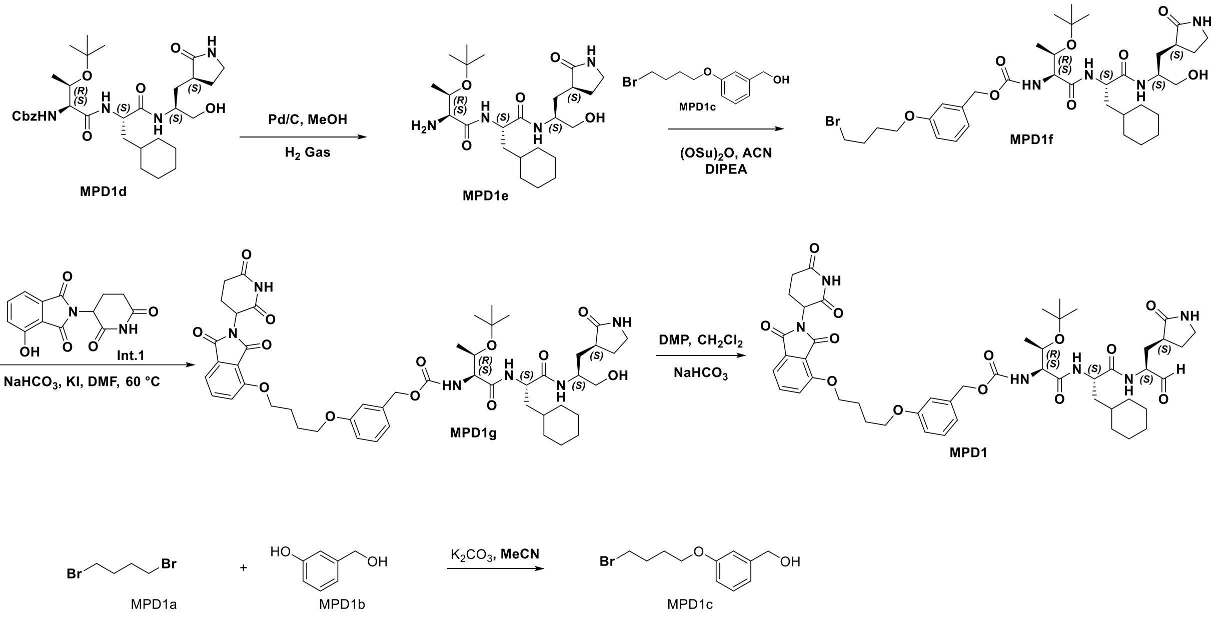


**Scheme S1:** The synthesis of compound **MPD1**

**Synthesis of (3-(4-bromobutoxy)phenyl)methanol (MPD1c):** To a solution of MPD1a (13.05 g, 60 mmol) and MPD1b (1.5 g, 12.09 mmol) in acetonitrile (80 Ml) was added $K_2CO_3$ (8.3 g, 60 mmol). The reaction mixture was then heated to reflux overnight. The reaction mixture was then evaporated *in vacuo*, the residue was then diluted with $H_2O$ and DCM. The organic layer was separated, washed with saturated brine solution, dried over anhydrous $Na_2SO_4$, and concentrated *in vacuo*. The residue was purified by flash chromatography (0~10% EtOAc in hexanes as eluent) to yield **MPD1c** as white solid (2.5 g, 66%). $^1$H NMR (400 MHz, $CDCl_3$) δ 7.23 – 7.14 (m, 1H), 6.89 – 6.82 (m, 2H), 6.78 – 6.70 (m, 1H), 4.59 (s, 2H), 3.93 (t, *J* = 6.0 Hz, 2H), 3.42 (t, *J* = 6.6 Hz, 2H), 2.06 – 1.93 (m, 2H), 1.92 – 1.81 (m, 2H).

**Synthesis of (2S,3R)-2-amino-3-(tert-butoxy)-N-((S)-3-cyclohexyl-1-(((S)-1-hydroxy-3-((S)-2-oxopyrrolidin-3-yl)propan-2-yl)amino)-1-oxopropan-2-yl)butanamide (MPD1e):** To a solution of **MPD1d** (602 mg, 1 mmol) in methanol (20 Ml) was added 10% Pd/C (100 mg). The reaction flask was sealed and the air inside was removed under Schlenk line. A hydrogen balloon was then attached to the reaction flask and the reaction mixture was stirred under room temperature for 3 h. After the completion of deprotection, the reaction mixture was filtered with a pad of Celite.

The filtrate was concentrated *in vacuo* and the residue was used for the next step without further purification.

**Synthesis of 3-(4-bromobutoxy)benzyl ((2S,3R)-3-(tert-butoxy)-1-(((S)-3-cyclohexyl-1-(((S)-1-hydroxy-3-((S)-2-oxopyrrolidin-3-yl)propan-2-yl)amino)-1-oxopropan-2-yl)amino)-1-oxobutan-2-yl)carbamate (MPD1f):** To a solution of **MPD1c** (200 mg, 0.775 mmol) in anhydrous CAN (5 Ml) was added NMM (236 mg, 2.32 mmol) at 0 °C. bis(2,5-dioxopyrrolidin-1-yl) carbonate (278 mg, 1.08 mmol) was slowly added to the solution at the same temperature. The reaction mixture was then allowed to warm up to room temperature and stirred overnight. Then Compound 5 (362 mg, 0.775) was added to reaction and stirred for 2 h. Solvent was removed vacuum and 20 ml Ethyl acetate added to residue. The reaction mixture was then washed by 1 M HCl (2×10 Ml). The organic layer was separated, dried over anhydrous $Na_2SO_4$, and concentrated *in vacuo*. The residue was purified by flash chromatography (50~100% EtOAc in hexanes as eluent) to yield **MPD1f** as white solid (250 mg, 49%). $^1$H NMR (400 MHz, $CDCl_3$) δ 7.49 (d, *J* = 7.9 Hz, 1H), 7.34 (d, *J* = 7.7 Hz, 1H), 7.21 – 7.14 (m, 1H), 6.88 – 6.73 (m, 3H), 6.19 (s, 1H), 6.06 (d, *J* = 5.3 Hz, 1H), 5.08 – 4.92 (m, 2H), 4.34 (td, *J* = 8.5, 5.6 Hz, 1H), 4.09 (dd, *J* = 6.1, 3.3 Hz, 2H), 4.00 – 3.84 (m, 3H), 3.60 – 3.49 (m, 2H), 3.34 (t, *J* = 6.8 Hz, 2H), 3.28 – 3.17 (m, 2H), 2.38 – 2.26 (m, 2H), 2.05 – 1.93 (m, 1H), 1.85 – 1.21 (m, 21H), 1.19 (s, 9H), 1.14 – 1.06 (m, 3H), 1.01 (d, *J* = 6.1 Hz, 4H), 0.96 – 0.75 (m, 2H).

**Synthesis of 3-(4-((2-(2,6-dioxopiperidin-3-yl)-1,3-dioxoisoindolin-4-yl)oxy)butoxy)benzyl ((2S,3R)-3-(tert-butoxy)-1-(((S)-3-cyclohexyl-1-(((S)-1-hydroxy-3-((S)-2-oxopyrrolidin-3-yl)propan-2-yl)amino)-1-oxopropan-2-yl)amino)-1-oxobutan-2-yl)carbamate (MPD1g):** To a solution of **MPD1f** (100 mg, 0.13 mmol) in DMF (2 Ml) was added **Int.1** (44 mg, 0.15 mmol), $NaHCO_3$ (23 mg, 0.26 mmol), KI (2 mg, 0.013 mmol). The reaction mixture was then stirred at 60 °C for 24 h. The reaction mixture was then diluted with EtOAc (20 Ml), washed with $H_2O$ (20 Ml) and saturated brine solution (20 Ml). The organic layer was separated, dried over anhydrous $Na_2SO_4$, and concentrated *in vacuo*. The residue was purified with flash chromatography (5~10% methanol in EtOAc as eluent) to yield **MPD1g** as white solid (85 mg, 55%). $^1$H NMR (400 MHz, DMSO) δ 11.10 (s, 1H), 7.82 (dt, *J* = 8.5, 7.0 Hz, 2H), 7.71 (d, *J* = 8.9 Hz, 1H), 7.55 – 7.48 (m, 2H), 7.44 (d, *J* = 7.2 Hz, 1H), 7.25 (t, *J* = 7.8 Hz, 1H), 6.95 – 6.81 (m, 4H), 5.12 – 4.95 (m, 3H), 4.66 (t, *J* = 5.6 Hz, 1H), 4.30 (q, *J* = 7.6 Hz, 1H), 4.21 (t, *J* = 6.4 Hz, 2H), 4.01 (dd, *J* = 9.1, 4.0

Hz, 1H), 3.97 (t, *J* = 2.1 Hz, 2H), 3.90 – 3.83 (m, 1H), 3.76 (s, 1H), 3.26 – 3.11 (m, 2H), 3.09 – 2.99 (m, 1H), 2.88 (ddd, *J* = 17.4, 14.1, 5.4 Hz, 1H), 2.58 (d, *J* = 13.8 Hz, 1H), 2.25 – 2.10 (m, 2H), 2.05 – 1.99 (m, 1H), 1.86 – 1.24 (m, 23H), 1.10 (s, 9H), 1.02 (d, *J* = 6.2 Hz, 3H), 0.88 (dt, *J* = 25.2, 13.9 Hz, 2H).

**Synthesis of 3-(4-((2-(2,6-dioxopiperidin-3-yl)-1,3-dioxoisoindolin-4-yl)oxy)butoxy)benzyl ((2S,3R)-3-(tert-butoxy)-1-(((S)-3-cyclohexyl-1-oxo-1-(((S)-1-oxo-3-((S)-2-oxopyrrolidin-3-yl)propan-2-yl)amino)propan-2-yl)amino)-1-oxobutan-2-yl)carbamate (MPD1):** To a solution of **MPD1g** (80 mg, 0.084 mmol) in anhydrous DCM (5 Ml) was added Dess-Martin periodinane (44 mg, 0.101 mmol). The reaction mixture was stirred at room temperature for 5 h. Then the reaction was quenched by the addition of 10% $Na_2S_2O_3$ in saturated $NaHCO_3$ solution (10 Ml). The mixture was further stirred until the organic layer became clear. The organic layer was then separated, dried with anhydrous $Na_2SO_4$, and concentrated *in vacuo*. The residue was then purified with flash chromatography (0~10 methanol in DCM as eluent) to yield **MPD1** as white solid (60 mg, 73%). $^1$H NMR (400 MHz, DMSO) δ 11.10 (s, 1H), 9.40 (s, 1H), 8.49 (d, *J* = 7.5 Hz, 1H), 8.08 – 7.89 (m, 1H), 7.81 (dd, *J* = 8.5, 7.3 Hz, 1H), 7.63 (s, 1H), 7.57 – 7.48 (m, 1H), 7.44 (d, *J* = 7.2 Hz, 1H), 7.25 (t, *J* = 7.8 Hz, 1H), 6.95 – 6.83 (m, 5H), 5.80 – 5.60 (m, 1H), 5.16 – 4.94 (m, 3H), 4.39 (d, *J* = 7.7 Hz, 1H), 4.20 (q, *J* = 7.0 Hz, 2H), 4.17 – 3.98 (m, 2H), 3.95 (t, *J* = 6.5 Hz, 2H), 3.87 (d, *J* = 5.9 Hz, 1H), 3.25 – 3.05 (m, 2H), 2.88 (ddd, *J* = 17.7, 13.9, 5.3 Hz, 1H), 2.58 (d, *J* = 13.8 Hz, 1H), 2.25 (d, *J* = 12.1 Hz, 1H), 2.10 (d, *J* = 20.2 Hz, 1H), 2.02 (dd, *J* = 11.7, 6.1 Hz, 1H), 1.96 – 1.28 (m, 20H), 1.10 (s, 9H), 1.06 – 0.99 (m, 3H), 0.86 (d, *J* = 13.5 Hz, 3H). $^{13}$C NMR (126 MHz, DMSO) δ 200.96, 178.60, 173.21, 172.94, 172.57, 170.38, 169.94, 167.32, 165.77, 159.20, 156.52, 156.38, 138.99, 137.48, 133.74, 129.87, 120.29, 119.97, 116.73, 115.60, 114.18, 114.05, 74.10, 69.30, 68.03, 67.85, 65.88, 60.16, 56.79, 50.83, 49.23, 37.69, 34.67, 33.87, 33.49, 32.70, 31.43, 29.67, 29.18, 29.14, 29.10, 28.88, 28.54, 27.70, 26.49, 26.22, 26.01, 25.95, 25.91, 25.71, 25.26, 22.52, 22.48, 19.92. ESI-HRMS m/z calculated for $C_{49}H_{65}N_6O_{13}$ $(M+H^+)$: 944.4531; found: 945.4581.

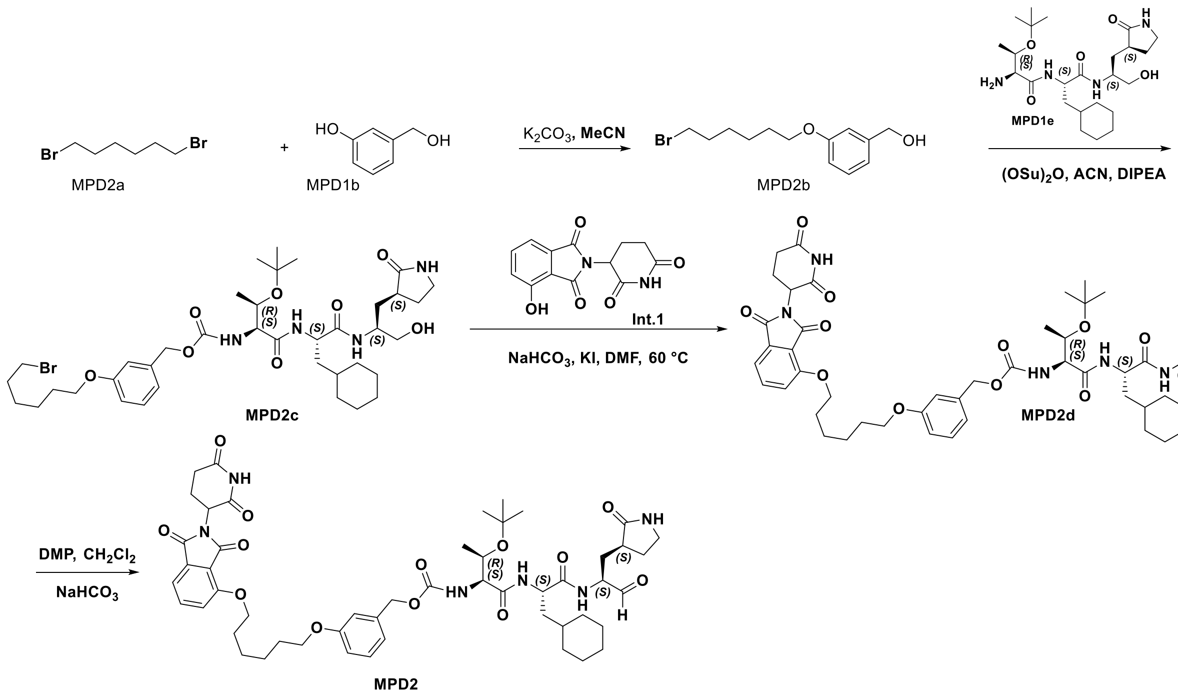


**Scheme S2:** The synthesis of compound **MPD2**

**Synthesis of (3-((6-bromohexyl)oxy)phenyl)methanol (MPD2b): MPD2b** was prepared as a white solid following a similar procedure to **MPD1c** (yield 77%). $^1$H NMR (400 MHz, Chloroform-*d*) δ 7.36 – 7.24 (m, 1H), 7.00 – 6.91 (m, 2H), 6.85 (dd, *J* = 8.4, 2.5 Hz, 1H), 4.69 (s, 2H), 4.00 (t, *J* = 6.4 Hz, 2H), 3.45 (t, *J* = 6.8 Hz, 2H), 1.92 (p, *J* = 6.9 Hz, 2H), 1.83 (p, *J* = 6.8 Hz, 2H), 1.59 – 1.47 (m, 4H).

**Synthesis of 3-((6-bromohexyl)oxy)benzyl ((2S,3R)-3-(tert-butoxy)-1-(((S)-3-cyclohexyl-1-(((S)-1-hydroxy-3-((S)-2-oxopyrrolidin-3-yl)propan-2-yl)amino)-1-oxopropan-2-yl)amino)-1-oxobutan-2-yl)carbamate (MPD2c): MPD2c** was prepared as a white solid following a similar procedure to **MPD1f** (yield 50%).$^1$H NMR (400 MHz, DMSO-$d_6$) δ 7.84 (d, *J* = 8.0 Hz, 1H), 7.71 (d, *J* = 8.9 Hz, 1H), 7.53 (s, 1H), 7.26 (t, *J* = 7.8 Hz, 1H), 6.99 – 6.90 (m, 2H), 6.90 – 6.78 (m, 2H), 5.12 – 4.90 (m, 2H), 4.65 (t, *J* = 5.6 Hz, 1H), 4.31 (q, *J* = 7.7 Hz, 1H), 4.01 (dd, *J* = 9.0, 4.1 Hz, 1H), 3.96 (t, *J* = 6.4 Hz, 2H), 3.87 (dd, *J* = 6.4, 4.4 Hz, 1H), 3.75 (d, *J* = 7.2 Hz, 1H), 3.55 (t, *J* = 6.7 Hz, 2H), 3.21 (dt, *J* = 10.4, 6.4 Hz, 1H), 3.14 (t, *J* = 9.1 Hz, 1H), 3.10 – 2.97 (m, 1H), 2.25

– 2.08 (m, 2H), 1.88 – 1.78 (m, 2H), 1.77 – 1.51 (m, 9H), 1.51 – 1.35 (m, 7H), 1.33 – 1.22 (m, 1H), 1.19 – 1.06 (m, 12H), 1.03 (d, *J* = 6.2 Hz, 3H), 0.96 – 0.77 (m, 3H).

**Synthesis of 3-((6-((2-(2,6-dioxopiperidin-3-yl)-1,3-dioxoisoindolin-4-yl)oxy)hexyl)oxy)benzyl ((2S,3R)-3-(tert-butoxy)-1-(((S)-3-cyclohexyl-1-(((S)-1-hydroxy-3-((S)-2-oxopyrrolidin-3-yl)propan-2-yl)amino)-1-oxopropan-2-yl)amino)-1-oxobutan-2-yl)carbamate (MPD2d): MPD2d** was prepared as a white solid following a similar procedure to **MPD1g** (yield 64%). $^{1}$H NMR (400 MHz, DMSO-$d_6$) δ 11.09 (s, 1H), 7.90 – 7.75 (m, 2H), 7.70 (d, *J* = 8.9 Hz, 1H), 7.55 – 7.47 (m, 2H), 7.44 (d, *J* = 7.2 Hz, 1H), 7.24 (t, *J* = 7.8 Hz, 1H), 6.96 – 6.79 (m, 4H), 5.11 – 4.94 (m, 3H), 4.74 – 4.63 (m, 1H), 4.30 (q, *J* = 7.7 Hz, 1H), 4.22 (t, *J* = 6.2 Hz, 2H), 4.03 – 3.93 (m, 3H), 3.91 – 3.82 (m, 1H), 3.80 – 3.69 (m, 1H), 3.25 – 3.17 (m, 2H), 3.17 – 3.08 (m, 1H), 3.08 – 2.97 (m, 1H), 2.87 (s, 2H), 2.13 (d, *J* = 15.5 Hz, 2H), 2.08 – 2.01 (m, 1H), 1.86 – 1.64 (m, 7H), 1.65 – 1.33 (m, 11H), 1.14 – 1.04 (m, 12H), 1.01 (d, *J* = 6.2 Hz, 3H), 0.93 – 0.75 (m, 4H).

**Synthesis of 3-((6-((2-(2,6-dioxopiperidin-3-yl)-1,3-dioxoisoindolin-4-yl)oxy)hexyl)oxy)benzyl ((2S,3R)-3-(tert-butoxy)-1-(((S)-3-cyclohexyl-1-oxo-1-(((S)-1-oxo-3-((S)-2-oxopyrrolidin-3-yl)propan-2-yl)amino)propan-2-yl)amino)-1-oxobutan-2-yl)carbamate (MPD2): MPD2** was prepared as a white solid following a similar procedure to **MPD1** (yield 60%).$^{1}$H NMR (400 MHz, DMSO-$d_6$) δ 11.11 (s, 1H), 9.38 (s, 1H), 8.51 (d, *J* = 8.0 Hz, 1H), 7.94 (d, *J* = 7.8 Hz, 1H), 7.80 (t, *J* = 8.0 Hz, 1H), 7.65 (s, 1H), 7.51 (d, *J* = 8.8 Hz, 1H), 7.44 (d, *J* = 7.3 Hz, 1H), 7.24 (t, *J* = 7.9 Hz, 1H), 6.88 (dd, *J* = 22.4, 9.4 Hz, 4H), 5.15 – 4.92 (m, 3H), 4.44 – 4.26 (m, 1H), 4.26 – 4.17 (m, 2H), 4.17 – 4.09 (m, 1H), 4.07 – 3.99 (m, 1H), 3.95 (t, *J* = 6.4 Hz, 2H), 3.91 – 3.80 (m, 1H), 3.24 – 2.98 (m, 2H), 2.94 – 2.80 (m, 1H), 2.63 – 2.58 (m, 1H), 2.58 – 2.53 (m, 1H), 2.31 – 2.18 (m, 1H), 2.18 – 2.07 (m, 1H), 2.06 – 1.96 (m, 1H), 1.82 – 1.44 (m, 17H), 1.14 – 1.00 (m, 16H), 0.96 – 0.81 (m, 4H). $^{13}$C NMR (126 MHz, DMSO) δ 200.97, 178.61, 173.21, 172.94, 170.38, 169.94, 167.32, 165.78, 162.59, 159.20, 156.50, 156.39, 138.98, 137.49, 133.74, 129.88, 120.30, 119.99, 116.75, 115.62, 114.16, 74.10, 69.24, 68.03, 67.78, 65.89, 60.16, 56.79, 50.83, 49.23, 37.69, 33.87, 33.49, 32.70, 31.44, 29.67, 29.11, 28.84, 28.54, 27.70, 26.49, 26.22, 26.01, 25.65, 25.52, 22.48, 21.18, 19.91. ESI-HRMS m/z calculated for $C_{51}H_{69}N_6O_{13}$ ($M+H^+$): 973.4917; found: 973.4903.

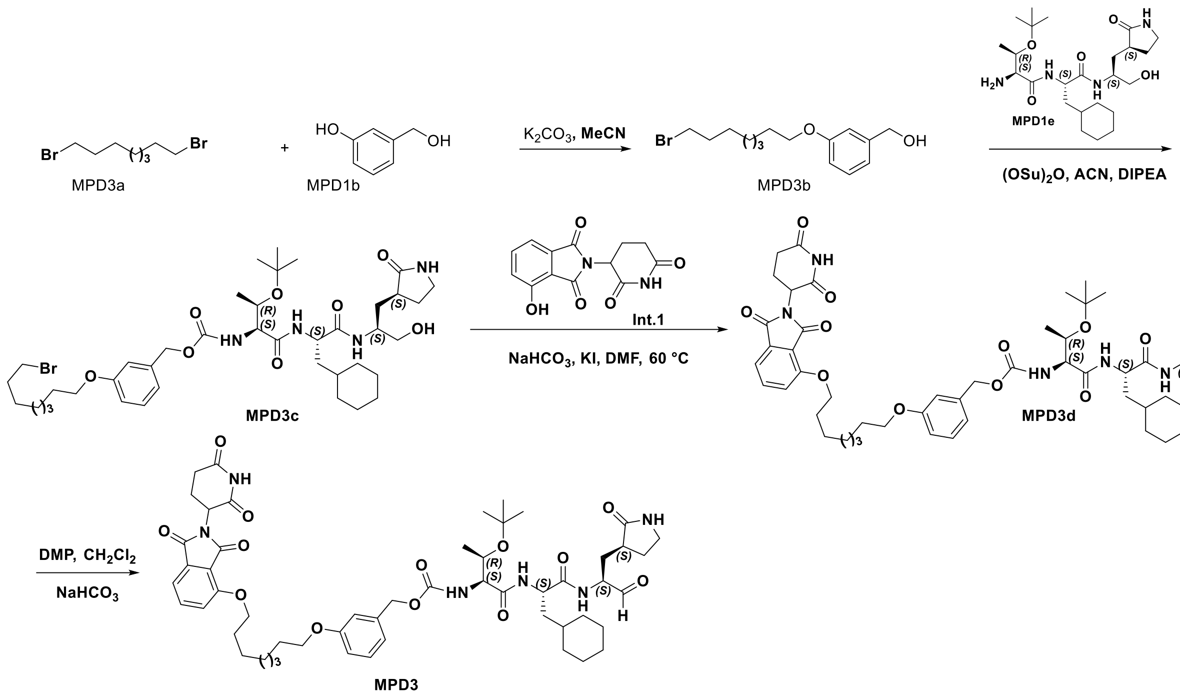


**Scheme S3:** The synthesis of compound **MPD3**

**Synthesis of (3-((8-bromooctyl)oxy)phenyl)methanol (MPD3b): MPD3b** was prepared as a white solid following a similar procedure to **MPD1c** (yield 66%). $^1$H NMR (400 MHz, $CDCl_3$) δ 7.23 – 7.14 (m, 1H), 6.88 – 6.82 (m, 2H), 6.75 (ddd, *J* = 8.2, 2.6, 1.0 Hz, 1H), 4.59 (s, 2H), 3.89 (t, *J* = 6.5 Hz, 2H), 3.34 (t, *J* = 6.8 Hz, 2H), 1.79 (p, *J* = 6.9 Hz, 2H), 1.71 (dq, *J* = 8.1, 6.5 Hz, 2H), 1.46 – 1.22 (m, 8H).

**Synthesis of 3-((8-bromooctyl)oxy)benzyl ((2S,3R)-3-(tert-butoxy)-1-(((S)-3-cyclohexyl-1-(((S)-1-hydroxy-3-((S)-2-oxopyrrolidin-3-yl)propan-2-yl)amino)-1-oxopropan-2-yl)amino)-1-oxobutan-2-yl)carbamate (MPD3c): MPD3c** was prepared as a white solid following a similar procedure to **MPD1f** (yield 49%).. $^{1}$H NMR (400 MHz, $CDCl_3$) δ 7.51 (d, *J* = 7.8 Hz, 1H), 7.35 (dd, *J* = 7.8, 2.4 Hz, 1H), 7.23 – 7.14 (m, 1H), 6.90 – 6.73 (m, 3H), 6.24 – 6.03 (m, 2H), 5.07 – 4.92 (m, 2H), 4.34 (td, *J* = 8.5, 5.7 Hz, 1H), 4.07 (dd, *J* = 19.3, 6.3 Hz, 2H), 3.93 (t, *J* = 6.0 Hz, 3H), 3.54 (t, *J* = 3.4 Hz, 1H), 3.42 (t, *J* = 6.6 Hz, 2H), 3.22 (d, *J* = 8.3 Hz, 2H), 2.38 – 2.27 (m, 2H), 1.99 (ddt, *J* = 10.2, 7.2, 5.0 Hz, 3H), 1.87 (dq, *J* = 9.7, 6.1 Hz, 2H), 1.78 – 1.43 (m, 9H), 1.19 (s, 9H), 1.15 – 1.05 (m, 3H), 1.01 (d, *J* = 6.0 Hz, 3H), 0.96 – 0.76 (m, 2H).

**Synthesis of 3-((8-((2-(2,6-dioxopiperidin-3-yl)-1,3-dioxoisoindolin-4-yl)oxy)octyl)oxy)benzyl ((2S,3R)-3-(tert-butoxy)-1-(((S)-3-cyclohexyl-1-(((S)-1-hydroxy-3-((S)-2-oxopyrrolidin-3-yl)propan-2-yl)amino)-1-oxopropan-2-yl)amino)-1-oxobutan-2-yl)carbamate (MPD3d): MPD3d** was prepared as a white solid following a similar procedure to **MPD1g** (yield 45%). $^{1}$H NMR (400 MHz, DMSO) δ 11.03 (s, 1H), 7.75 (dd, *J* = 8.6, 7.3 Hz, 2H), 7.63 (d, *J* = 9.0 Hz, 1H), 7.49 – 7.42 (m, 2H), 7.38 (d, *J* = 7.2 Hz, 1H), 7.18 (t, *J* = 7.9 Hz, 1H), 6.90 – 6.73 (m, 4H), 5.01 (dd, *J* = 12.8, 5.4 Hz, 1H), 4.98 – 4.88 (m, 2H), 4.58 (t, *J* = 5.6 Hz, 1H), 4.23 (d, *J* = 7.1 Hz, 3H), 4.02 – 3.90 (m, 4H), 3.79 (t, *J* = 5.4 Hz, 1H), 3.68 (s, 1H), 3.18 – 3.11 (m, 1H), 3.06 (t, *J* = 9.0 Hz, 1H), 3.01 – 2.94 (m, 1H), 2.81 (ddd, *J* = 17.7, 13.8, 5.3 Hz, 1H), 2.56 – 2.45 (m, 1H), 2.08 (s, 2H), 1.99 – 1.92 (m, 1H), 1.86 (s, 3H), 1.57 (ddd, *J* = 62.7, 22.3, 10.9 Hz, 7H), 1.39 – 1.26 (m, 3H), 1.14 (d, *J* = 30.4 Hz, 2H), 1.03 (s, 9H), 1.00 – 0.92 (m, 6H), 0.77 (t, *J* = 12.3 Hz, 2H).

**Synthesis of 3-((8-((2-(2,6-dioxopiperidin-3-yl)-1,3-dioxoisoindolin-4-yl)oxy)octyl)oxy)benzyl ((2S,3R)-3-(tert-butoxy)-1-(((S)-3-cyclohexyl-1-oxo-1-(((S)-1-oxo-3-((S)-2-oxopyrrolidin-3-yl)propan-2-yl)amino)propan-2-yl)amino)-1-oxobutan-2-yl)carbamate (MPD3): MPD3** was prepared as a white solid following a similar procedure to **MPD1** (yield 73%). $^{1}$H NMR (400 MHz, DMSO) δ 11.03 (s, 1H), 9.33 (s, 1H), 8.42 (d, *J* = 7.6 Hz, 1H), 7.84 (d, *J* = 7.8 Hz, 1H), 7.75 (dd, *J* = 8.5, 7.3 Hz, 1H), 7.56 (s, 1H), 7.42 (dd, *J* = 31.9, 7.9 Hz, 2H), 7.18 (t, *J* = 7.9 Hz, 1H), 6.83 (ddd, *J* = 17.9, 13.3, 8.4 Hz, 4H), 5.72 – 5.50 (m, 1H), 5.06 – 4.87 (m, 3H), 4.31 (q, *J* = 7.6 Hz, 1H), 4.22 (s, 2H), 4.07 (ddd, *J* = 11.6, 7.6, 4.0 Hz, 1H), 4.02 – 3.90 (m, 3H), 3.84 – 3.77 (m, 1H), 3.06 (dt, *J* = 25.5, 8.5 Hz, 1H), 2.82 (ddd, *J* = 16.8, 13.8,

5.3 Hz, 1H), 2.44 (d, *J* = 1.9 Hz, 2H), 2.16 (dd, *J* = 14.4, 6.7 Hz, 1H), 2.06 (s, 1H), 1.99 – 1.92 (m, 1H), 1.88 – 1.76 (m, 5H), 1.73 – 1.14 (m, 13H), 1.15 – 0.90 (m, 16H), 0.86 – 0.67 (m, 3H). $^{13}$C NMR (126 MHz, DMSO) δ 200.97, 179.46, 178.61, 173.21, 172.94, 172.61, 172.26, 170.38, 169.94, 169.68, 167.32, 165.81, 159.12, 156.41, 156.38, 138.94, 137.51, 133.73, 129.89, 120.28, 120.05, 116.77, 115.68, 114.20, 74.11, 74.05, 69.04, 68.03, 67.55, 65.91, 60.15, 56.79, 52.73, 51.07, 50.83, 49.24, 37.69, 33.87, 33.76, 33.49, 32.70, 31.44, 29.66, 28.57, 28.54, 28.14, 27.70, 26.49, 26.23, 26.01, 25.81, 25.67, 22.49, 19.91, 19.84. ESI-HRMS calculated for $C_{53}H_{72}N_6O_{13}$ (M-H$^+$): 999.5157; found: 999.5088. ESI-HRMS m/z calculated for $C_{53}H_{72}N_6O_{13}^+$ (M + H)$^+$ 1001.5191, found 1001.522.

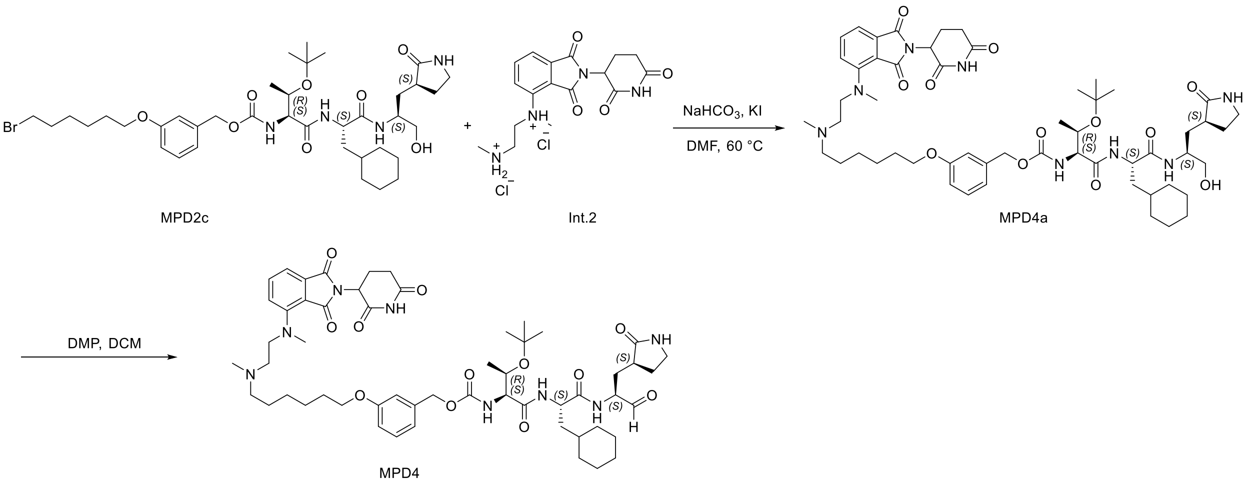


**Scheme 4:** The synthesis of compound **MPD4**

**Synthesis of 3-((6-((2-((2-(2,6-dioxopiperidin-3-yl)-1,3-dioxoisoindolin-4-yl)(methyl)amino)ethyl)(methyl)amino)hexyl)oxy)benzyl ((2S,3R)-3-(tert-butoxy)-1-(((S)-3-cyclohexyl-1-(((S)-1-hydroxy-3-((S)-2-oxopyrrolidin-3-yl)propan-2-yl)amino)-1-oxopropan-2-yl)amino)-1-oxobutan-2-yl)carbamate (MPD4a): MPD4a** was prepared as a white solid following a similar procedure to **MPD1g** (yield 35%).

**Synthesis of 3-((6-((2-((2-(2,6-dioxopiperidin-3-yl)-1,3-dioxoisoindolin-4-yl)(methyl)amino)ethyl)(methyl)amino)hexyl)oxy)benzyl ((2S,3R)-3-(tert-butoxy)-1-(((S)-3-cyclohexyl-1-oxo-1-(((S)-1-oxo-3-((S)-2-oxopyrrolidin-3-yl)propan-2-yl)amino)propan-2-**

**yl)amino)-1-oxobutan-2-yl)carbamate (MPD4): MPD4**was prepared as a white solid following a similar procedure to **MPD1** (yield 56%). $^{1}$H NMR (400 MHz, Chloroform-*d*) δ 9.62 – 9.37 (m, 1H), 8.15 – 7.96 (m, 1H), 7.37 (t, *J* = 8.0 Hz, 2H), 7.23 (d, *J* = 7.8 Hz, 1H), 7.14 (d, *J* = 7.5 Hz, 1H), 6.97 (d, *J* = 8.2 Hz, 1H), 6.93 – 6.86 (m, 2H), 6.83 (d, *J* = 8.3 Hz, 1H), 6.71 (s, 1H), 5.95 (d, *J* = 25.4 Hz, 2H), 5.17 – 4.99 (m, 2H), 4.78 (t, *J* = 8.5 Hz, 1H), 4.57 – 4.31 (m, 2H), 4.18 (s, 2H), 3.93 (t, *J* = 6.5 Hz, 2H), 3.76 (s, 3H), 3.59 – 3.45 (m, 2H), 3.38 – 3.26 (m, 3H), 3.18 (s, 4H), 3.05 – 2.97 (m, 1H), 2.82 (s, 3H), 2.57 – 2.16 (m, 6H), 2.02 – 1.86 (m, 4H), 1.84 – 1.58 (m, 11H), 1.54 – 1.31 (m, 7H), 1.31 – 1.05 (m, 19H), 1.04 – 0.64 (m, 4H). ESI-HRMS calculated for $C_{55}H_{78}N_8O_{12}Na$ (M+Na$^+$): 1065.5631; found: 1065.5625.

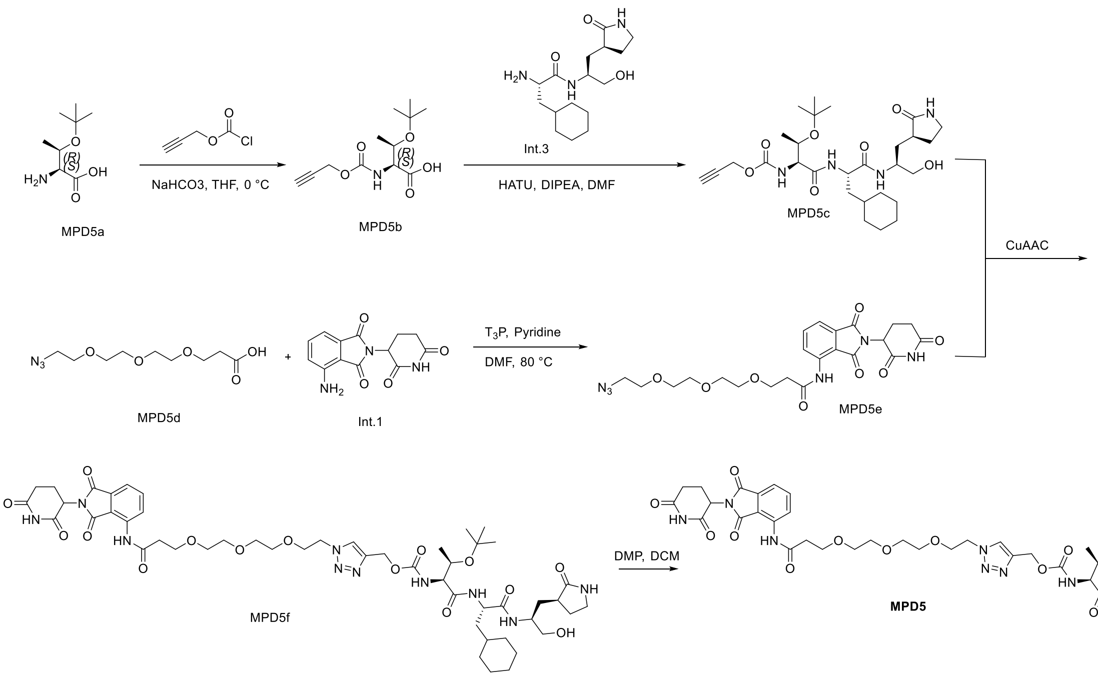


**Scheme 5:** The synthesis of compound **MPD5**

**Synthesis of O-(tert-butyl)-N-((prop-2-yn-1-yloxy)carbonyl)-L-threonine (MPD5b):** Dissolve O-(tert-butyl)-L-threonine (MPD5a) (2.0 g, 11.41 mmol) in 40mL saturated sodium bicarbonate solution, The temperature was lowered to 0° C., a THF solution of propargyl chloroformate (13.7 mmol) was added dropwise, and the mixture was stirred vigorously. After dropping, the reaction solution was heated to 25°C, and the reaction was continued for 2 hours. After the reaction is complete, adjust the pH of the reaction solution to 11 with saturated sodium hydroxide solution. Wash the water phase with 40mL ethyl acetate three times, after washing, adjust the pH of the reaction solution to 2 with concentrated hydrochloric acid. The aqueous phase was extracted three times with 50 mL of ethyl acetate, the organic phases were combined and dried with anhydrous sodium sulfate, The organic solvent was evaporated under reduced pressure, 2.3 g of white solid was obtained with a yield of 78%.$^{1}$H NMR (400 MHz, $CDCl_3$) δ 5.76 (d, *J* = 6.8 Hz, 1H), 4.72 (t, *J* = 2.9 Hz, 2H), 4.32 (ddt, *J* = 15.9, 6.5, 3.4 Hz, 2H), 2.51 (q, *J* = 3.1 Hz, 1H), 1.27 (s, 9H), 1.19 (d, *J* = 6.3 Hz, 3H). $^{13}$C NMR (101 MHz, $CDCl_3$) δ 172.80, 155.41, 77.80, 76.33, 74.99, 66.68, 60.45, 58.71, 28.10, 18.42.

**Synthesis of Prop-2-yn-1-yl ((2S,3S)-3-(tert-butoxy)-1-(((S)-3-cyclohexyl-1-(((S)-1-hydroxy-3-((S)-2-oxopyrrolidin-3-yl)propan-2-yl)amino)-1-oxopropan-2-yl)amino)-1-oxobutan-2-yl)carbamate (MPD5c):** The acid compound **MPD5b** (0.181 g, 0.70 mmol) and the **amine Int.3** (0.2 g, 0.64 mmol) were dissolved in dry DMF (10 mL) and the reaction was cooled to 0 °C. HATU (317 mg, 0.83 mmol) and DIPEA (0.46 mL, 2.57 mmol) were added, and the reaction mixture was allowed warm up to room temperature and stirred for 12 h. The mixture was then poured into water (50 mL) and extracted with ethyl acetate (4×20 mL). The organic layer was washed with aqueous hydrochloric acid 10% v/v (2×20 mL), saturated aqueous $NaHCO_3$ (2×20 mL), brine (2×20 mL) and dried over $Na_2SO_4$. The organic phase was evaporated to dryness and the crude material purified by silica gel column chromatography (0-10% MeOH in $CH_2Cl_2$ as the eluent) to afford **MPD5c** white gummy solid (180 mg, 56%). $^{1}$H NMR (400 MHz, $CDCl_3$) δ 7.57 (d, *J* = 7.9 Hz, 1H), 7.37 (d, *J* = 7.6 Hz, 1H), 6.26 (d, *J* = 5.2 Hz, 1H), 6.10 (s, 1H), 4.62 (qd, *J* = 15.6, 2.5 Hz, 2H), 4.34 (q, *J* = 7.7 Hz, 1H), 4.08 (d, *J* = 5.5 Hz, 2H), 3.95 (s, 1H), 3.64 – 3.51 (m, 2H), 3.31 – 3.20 (m, 2H), 2.44 (t, *J* = 2.4 Hz, 1H), 2.36 (d, *J* = 8.4 Hz, 2H), 2.03 (t, *J* = 12.6 Hz, 1H), 1.84 – 1.72 (m, 1H), 1.69 – 1.31 (m, 9H), 1.21 – 1.10 (m, 12H), 1.01 (d, *J* = 5.9 Hz, 3H), 0.85 (dq, *J* = 22.5, 11.2 Hz, 2H). $^{13}$C NMR (101 MHz, $CDCl_3$) δ 181.05, 172.74, 169.58, 155.51, 78.08, 75.49, 74.95, 66.69, 65.84, 59.15, 55.45, 52.78, 51.84, 50.30, 40.57, 39.68, 38.25, 34.22, 33.59, 32.64, 28.52, 28.23, 26.36, 26.22, 26.03, 18.64.

**Synthesis of 3-(2-(2-(2-azidoethoxy)ethoxy)ethoxy)-N-(2-(2,6-dioxopiperidin-3-yl)-1,3-dioxoisoindolin-4-yl)propenamide (MPD5e):** In a 50 mL round-bottom flask, 4-amino-2-(2,6-dioxo-3-piperidyl)-isoindoline-1,3-dione (Int.1) (500 mg, 1 eq) was dissolved in DMF (5 mL) and cooled to 0 °C in an ice bath. 3-{2-[2-(2-Azidoethoxy)-ethoxy]ethoxy}propanoic acid **(MPD5d)** (905 mg, 2 eq) was added dropwise. Then, N-propylphosphonic acid anhydride, cyclic trimer (7.0 g, 6.5 mL, 6 eq) and pyridine (1.5 mL, 10 eq) were added. The mixture was stirred at 80 °C for 3 h. The reaction mixture concentrated and purified by by silica gel column chromatography (0-10% MeOH in $CH_2Cl_2$ as the eluent) to afford **MPD5e** white gummy solid (600 mg, 65%). $^1$H NMR (400 MHz, DMSO) δ 11.21 (s, 1H), 9.94 (s, 1H), 8.61 (d, *J* = 8.4 Hz, 1H), 7.90 (t, *J* = 7.9 Hz, 1H), 7.68 (d, *J* = 7.3 Hz, 1H), 5.21 (dd, *J* = 12.7, 5.4 Hz, 1H), 3.81 (t, *J* = 6.0 Hz, 2H), 3.73 – 3.51 (m, 11H), 3.43 (dt, *J* = 14.5, 4.9 Hz, 2H), 2.96 (ddd, *J* = 16.7, 13.7, 5.4 Hz, 1H), 2.77 (t, *J* = 6.0 Hz, 2H), 2.71 – 2.57 (m, 2H), 2.18 – 2.09 (m, 1H).

**Synthesis of (1-(2-(2-(2-(3-((2-(2,6-dioxopiperidin-3-yl)-1,3-dioxoisoindolin-4-yl)amino)-3-oxopropoxy)ethoxy)ethoxy)ethyl)-1H-1,2,3-triazol-4-yl)methyl ((2S,3R)-3-(tert-butoxy)-1-(((S)-3-cyclohexyl-1-(((S)-1-hydroxy-3-((S)-2-oxopyrrolidin-3-yl)propan-2-yl)amino)-1-oxopropan-2-yl)amino)-1-oxobutan-2-yl)carbamate (MPD5f):** The alkyne compound **(MPD5c)** (80 mg, 0.145 mmol) and the azide **(MPD5e)** (73 mg, 0.145 mmol) were dissolved in tert-butanol (5 mL). Ascorbic acid (28 mg, 0.145 mmol) and copper sulfate (3.5 mg, 0.015 mmol) was dissolved in water (3 mL) and the two solutions were added together to stir overnight with monitoring by TLC. The reaction mixture was diluted with water and cooled in ice for 10 minutes, and the brown/black precipitate was collected by filtration. The precipitate was extracted with chloroform, washed with cold water (2x5 mL), dried with $MgSO_4$ and concentrated via rotavapor. The residue was purified by flash column chromatography (0 to 10 % MeOH in $CH_2Cl_2$) to afford **MPD5f** as gummy solid 50 mg with a 33% yield. $^1$H NMR (400 MHz, $CDCl_3$) δ 10.49 (d, *J* = 55.1 Hz, 1H), 9.93 (d, *J* = 10.4 Hz, 1H), 8.83 (dd, *J* = 8.5, 4.0 Hz, 1H), 7.73 – 7.64 (m, 1H), 7.61 – 7.33 (m, 3H), 5.02 (s, 1H), 4.53 – 4.48 (m, 2H), 4.36 – 4.31 (m, 1H), 4.17 (d, *J* = 6.7 Hz, 1H), 4.11 – 3.96 (m, 2H), 3.89 – 3.79 (m, 4H), 3.70 – 3.56 (m, 8H), 3.30 – 3.25 (m, 2H), 2.88 – 2.83 (m, 2H), 2.73 (q, *J* = 6.1 Hz, 2H), 2.37 – 2.33 (m, 2H), 2.15 – 2.10 (m, 2H), 1.67 (t, *J* = 27.1 Hz, 11H), 1.32 (s, 1H), 1.26 – 1.12 (m, 1H), 1.19 – 1.07 (m, 14H), 0.91 (d, *J* = 15.1 Hz, 2H).

**Synthesis of (1-(2-(2-(2-(3-((2-(2,6-dioxopiperidin-3-yl)-1,3-dioxoisoindolin-4-yl)amino)-3-oxopropoxy)ethoxy)ethoxy)ethyl)-1H-1,2,3-triazol-4-yl)methyl ((2S,3R)-3-(tert-butoxy)-1-(((S)-3-cyclohexyl-1-oxo-1-(((S)-1-oxo-3-((S)-2-oxopyrrolidin-3-yl)propan-2-**

yl)amino)propan-2-yl)amino)-1-oxobutan-2-yl)carbamate(MPD5): MPD5** was prepared as a white solid following a similar procedure to **MPD1** (yield 60%).$^1$H NMR (400 MHz, $CDCl_3$) δ 10.14 (d, *J* = 21.4 Hz, 1H), 9.97 (s, 1H), 9.52 (s, 1H), 8.87 (dd, *J* = 8.5, 2.8 Hz, 1H), 8.04 (d, *J* = 7.8 Hz, 1H), 7.80 (d, *J* = 6.0 Hz, 1H), 7.73 (d, *J* = 8.5 Hz, 1H), 7.60 – 7.53 (m, 2H), 6.61 (d, *J* = 14.9 Hz, 1H), 5.94 (dd, *J* = 15.5, 6.9 Hz, 1H), 5.27 (d, *J* = 12.7 Hz, 1H), 5.17 (dd, *J* = 12.8, 6.5 Hz, 1H), 5.06 – 5.02 (m, 1H), 4.54 – 4.50 (m, 2H), 4.49 – 4.44 (m, 1H), 4.38 – 4.34 (m, 1H), 4.24 – 4.19 (m, 1H), 4.16 – 4.11 (m, 4H), 3.92 – 3.85 (m, 5H), 3.74 – 3.69 (m, 2H), 3.69 – 3.57 (m, 6H), 3.33 (d, *J* = 9.2 Hz, 2H), 2.91 – 2.83 (m, 2H), 2.81 – 2.74 (m, 2H), 2.43 – 2.39 (m, 2H), 2.19 – 2.14 (m, 1H), 1.96 – 1.92 (m, 2H), 1.87 – 1.65 (m, 11H), 1.22 – 1.11 (m, 14H), 0.99 – 0.90 (m, 2H). ESI-HRMS m/z calculated for $C_{50}H_{70}N_{10}O_{15}^+$ $(M + H)^+$ 1051.5056, found 1051.5066.

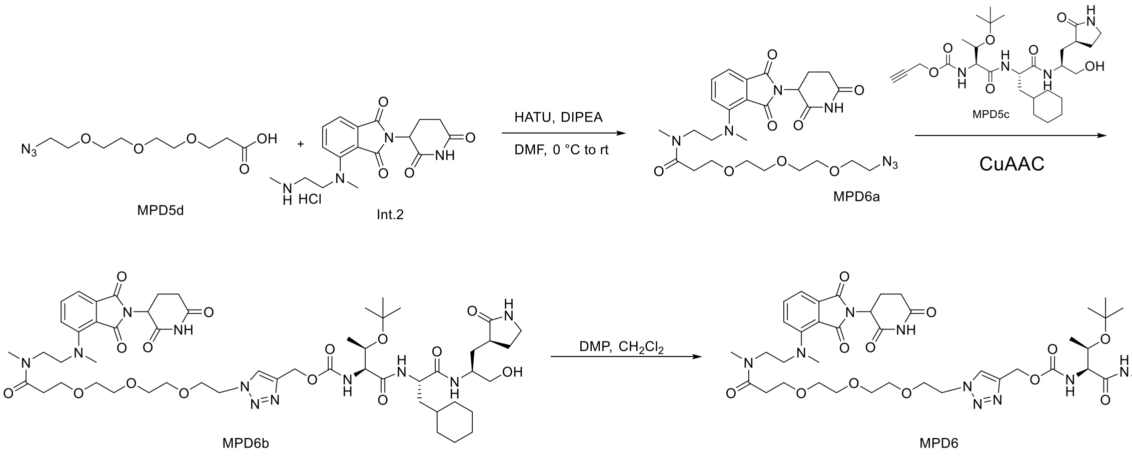


**Scheme 6:** The synthesis of compound **MPD6**

**Synthesis of 3-(2-(2-(2-azidoethoxy)ethoxy)ethoxy)-N-(2-((2-(2,6-dioxopiperidin-3-yl)-1,3-dioxoisoindolin-4-yl)(methyl)amino)ethyl)-N-methylpropanamide (MPD6a): ): MPD6a** was prepared as a white solid following a similar procedure to **MPD5c** (yield 66%).$^1$H NMR (400 MHz, $CDCl_3$) δ 8.21 (s, 1H), 8.04 (s, 1H), 7.64 – 7.51 (m, 1H), 7.45 – 7.32 (m, 1H), 7.20 (dd, *J* = 8.6, 3.8 Hz, 1H), 5.02 – 4.91 (m, 1H), 3.80 – 3.52 (m, 14H), 3.40 (q, *J* = 4.7 Hz, 2H), 3.14 (s, 2H), 2.97 (d, *J* = 2.0 Hz, 3H), 2.90 (d, *J* = 0.9 Hz, 5H), 2.85 – 2.67 (m, 2H), 2.38 (qt, *J* = 15.8, 6.9 Hz, 2H), 2.22 – 2.11 (m, 1H).

**Synthesis of (1-(2-(2-(2,6-dioxopiperidin-3-yl)-1,3-dioxoisoindolin-4-yl)-5-methyl-6-oxo-9,12,15-trioxa-2,5-diazaheptadecan-17-yl)-1H-1,2,3-triazol-4-yl)methyl ((2S,3R)-3-(tert-butoxy)-1-(((S)-3-cyclohexyl-1-(((S)-1-hydroxy-3-((S)-2-oxopyrrolidin-3-yl)propan-2-yl)amino)-1-oxopropan-2-yl)amino)-1-oxobutan-2-yl)carbamate (MPD6b): MPD6b** was prepared as a white solid following a similar procedure to **MPD5f** (yield 44%). $^{1}$H NMR (400 MHz, $CDCl_3$) δ 10.03 – 9.40 (m, 1H), 7.66 – 7.26 (m, 3H), 7.08 (d, *J* = 8.8 Hz, 1H), 5.99 (s, 1H), 5.15 (s, 1H), 4.92 (s, 1H), 4.51 – 4.47 (m, 2H), 4.35 – 4.30 (m, 1H), 4.17 – 4.11 (m, 1H), 4.08 – 3.91 (m, 2H), 3.82 – 3.69 (m, 4H), 3.66 – 3.43 (m, 14H), 3.26 – 3.20 (m, 2H), 3.05 – 2.93 (m, 3H), 2.87 – 2.69 (m, 5H), 2.61 – 2.52 (m, 1H), 2.38 – 2.20 (m, 4H), 2.08 – 2.01 (m, 1H), 1.75 – 1.43 (m, 10H), 1.31 – 1.23 (m, 1H), 1.19 – 1.03 (m, 17H), 0.91 – 0.80 (m, 2H).

**Synthesis of (1-(2-(2-(2,6-dioxopiperidin-3-yl)-1,3-dioxoisoindolin-4-yl)-5-methyl-6-oxo-9,12,15-trioxa-2,5-diazaheptadecan-17-yl)-1H-1,2,3-triazol-4-yl)methyl ((2S,3R)-3-(tert-butoxy)-1-(((S)-3-cyclohexyl-1-oxo-1-(((S)-1-oxo-3-((S)-2-oxopyrrolidin-3-yl)propan-2-yl)amino)propan-2-yl)amino)-1-oxobutan-2-yl)carbamate (MPD6): MPD6** was prepared as a white solid following a similar procedure to **MPD1** (yield 56%). $^{1}$H NMR (400 MHz, $CDCl_3$) δ 9.43 (s, 1H), 9.19 (d, *J* = 44.4 Hz, 1H), 8.02 (s, 1H), 7.74 (d, *J* = 4.5 Hz, 1H), 7.48 (dt, *J* = 20.9, 7.8 Hz, 1H), 7.41 – 7.28 (m, 1H), 7.08 (d, *J* = 8.8 Hz, 1H), 6.40 (d, *J* = 14.9 Hz, 1H), 5.90 – 5.82 (m, 1H), 5.26 – 5.13 (m, 1H), 5.12 – 5.03 (m, 1H), 4.96 – 4.85 (m, 1H), 4.58 – 4.37 (m, 3H), 4.30 – 4.26 (m, 1H), 4.14 (s, 1H), 4.09 – 4.05 (m, 1H), 3.84 – 3.70 (m, 3H), 3.62 – 3.45 (m, 11H), 3.25 (d, *J* = 8.7 Hz, 2H), 3.06 – 2.94 (m, 3H), 2.90 – 2.66 (m, 5H), 2.62 – 2.52 (m, 1H), 2.35 – 2.22 (m, 3H), 2.05 (d, *J* = 9.5 Hz, 1H), 1.95 – 1.49 (m, 12H), 1.32 – 1.26 (m, 1H), 1.18 – 1.00 (m, 18H), 0.93 – 0.80 (m, 2H). ESI-HRMS m/z calculated for $C_{54}H_{79}N_{11}O_{15}^{+}$ $(M + H)^{+}$ 1122.5791, found 1122.5812.

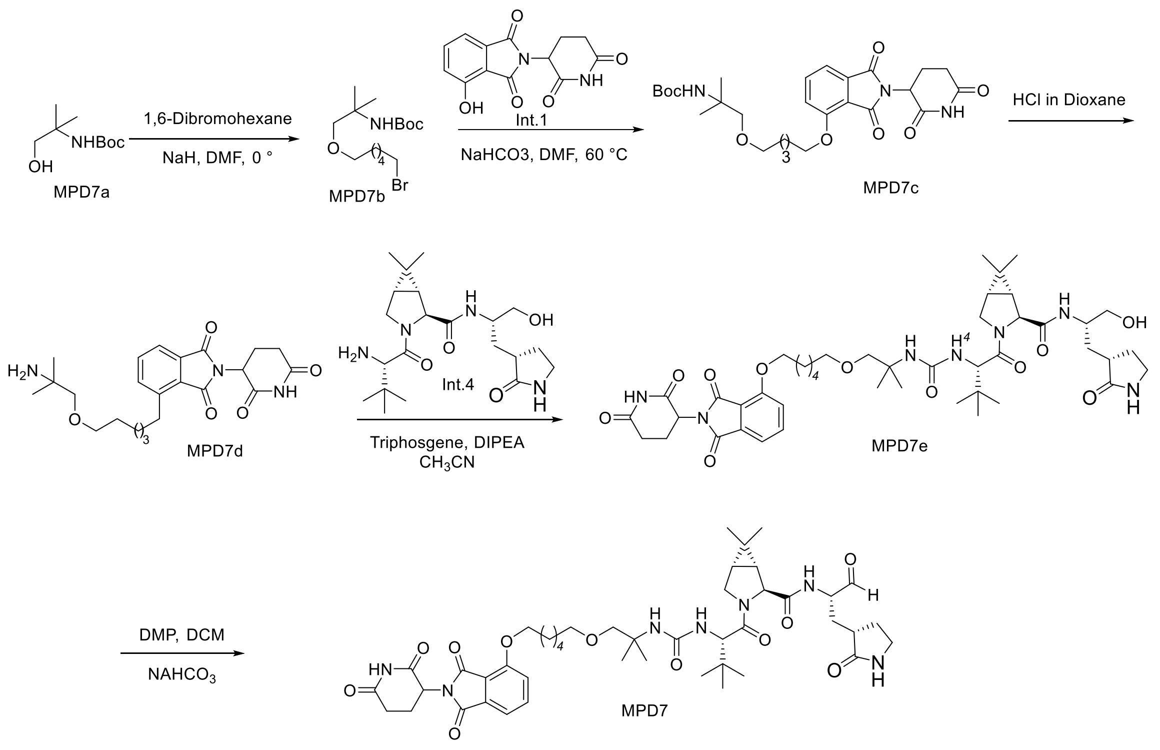


**Scheme 7:** The synthesis of compound **MPD7**

**Synthesis of tert-butyl (1-((6-bromohexyl)oxy)-2-methylpropan-2-yl)carbamate (MPD7b):** To a solution of **NaH** (100%, 116 mg, 2.91 mmol) in DMF (20 mL) was added compound **MPD7a** (500 mg, 7.93 mmol) at 0 °C. Stir it 20 min at same temperature. Then 1,6-Dibromo hexane (1.21 mL, 31.74 mmol) was added to reaction as drop wise at 0 °C. Raise the temperature slowly to room temperature and stir for 2h. The mixture was then poured into water (100 mL) and extracted with ethyl acetate (2×50 mL). The organic layer was washed with aqueous hydrochloric acid 10% v/v (2×50 mL), saturated aqueous $NaHCO_3$ (2×50 mL), brine (2×50 mL) and dried over $Na_2SO_4$. The organic phase was evaporated to dryness and the crude material purified by silica gel column chromatography (0-35% EtOAc in Hexanes as the eluent) to afford **MPD7b** white gummy solid (900 mg, 95%). $^1$H NMR (400 MHz, $CDCl_3$) δ 4.67 (s, 1H), 3.36 (dt, *J* = 12.0, 6.6 Hz, 4H), 3.25 (s, 2H), 1.80 (p, *J* = 6.9 Hz, 2H), 1.57 – 1.46 (m, 2H), 1.46 – 1.27 (m, 13H), 1.21 (s, 6H). $^{13}$C NMR (101 MHz, $CDCl_3$) δ 157.80, 74.80, 58.41, 40.28, 33.81, 32.64, 29.61, 27.81, 26.16, 25.30.

**Synthesis of tert-butyl (1-((6-((2-(2,6-dioxopiperidin-3-yl)-1,3-dioxoisoindolin-4-yl)oxy)hexyl)oxy)-2-methylpropan-2-yl)carbamate (MPD7c): MPD7c** was prepared as a white solid following a similar procedure to **MPD1g** (yield 43%).$^{1}$H NMR (400 MHz, $CDCl_3$) δ 8.30 (s, 1H), 7.68 (dd, *J* = 8.5, 7.3 Hz, 1H), 7.46 (d, *J* = 7.3 Hz, 1H), 7.23 (d, *J* = 8.5 Hz, 1H), 4.97 (dd, *J* = 12.1, 5.4 Hz, 1H), 4.80 (s, 1H), 4.19 (t, *J* = 6.5 Hz, 2H), 3.47 (t, *J* = 6.5 Hz, 2H), 3.34 (s, 2H), 2.95 – 2.68 (m, 3H), 2.20 – 2.09 (m, 1H), 1.91 (p, *J* = 6.8 Hz, 2H), 1.69 – 1.52 (m, 4H), 1.46 – 1.42 (m, 11H), 1.29 (s, 6H). $^{13}$C NMR (101 MHz, $CDCl_3$) δ 171.01, 168.14, 167.09, 165.68, 156.71, 154.93, 136.48, 133.83, 118.94, 117.14, 115.73, 71.35, 69.37, 60.40, 52.73, 49.10, 31.41, 29.40, 28.88, 28.47, 25.82, 25.65, 24.22, 22.64, 21.06.

**Synthesis of 4-((6-(2-amino-2-methylpropoxy)hexyl)oxy)-2-(2,6-dioxopiperidin-3-yl)isoindoline-1,3-dione hydrogen chloride (MPD7d): ):** To a solution of **MPD7c** (600 mg, 1.10 mmol) in dioxane (3 mL) was added **HCl in dioxane** (2.75 mL, 4M) at room temperature, The reaction mixture was then stirred at RT for 2h. After completion of reaction by TLC. Remove the solvent by using rotavapor. The solid crude compound directly used for next step. $^{1}$H NMR (400 MHz, $CDCl_3$) δ 12.29 (s, 1H), 9.28 (s, 3H), 9.00 (dd, *J* = 8.5, 7.2 Hz, 1H), 8.71 (d, *J* = 8.5 Hz, 1H), 8.63 (d, *J* = 7.3 Hz, 1H), 6.27 (dd, *J* = 12.7, 5.4 Hz, 1H), 5.39 (t, *J* = 6.3 Hz, 2H), 4.64 (t, *J* = 6.4 Hz, 2H), 4.53 (s, 2H), 4.08 (ddd, *J* = 16.6, 13.7, 5.4 Hz, 1H), 3.83 – 3.62 (m, 3H), 3.22 (tdd, *J* = 7.9, 6.8, 3.5 Hz, 1H), 2.96 (p, *J* = 6.5 Hz, 2H), 2.81 – 2.58 (m, 6H), 2.40 (s, 6H).

**Synthesis of (1R,2S,5S)-3-((2S)-2-(3-(1-((6-((2-(2,6-dioxopiperidin-3-yl)-1,3-dioxoisoindolin-4-yl)oxy)hexyl)oxy)-2-methylpropan-2-yl)ureido)-3,3-dimethylbutanoyl)-N-((S)-1-hydroxy-3-((S)-2-oxopyrrolidin-3-yl)propan-2-yl)-6,6-dimethyl-3-azabicyclo[3.1.0]hexane-2-carboxamide (MPD7e):** To a solution of this deprotected product **MPD7d** (150 mg, 0.31 mmol) in anhydrous ACN (5 mL) was added DIPEA (0.167 mL, 0.93 mmol). Then a solution of triphosgene (46 mg, 0.156 mmol) in anhydrous ACN (2 mL) was added to the solution at 0 °C. The resulting reaction mixture was stirred at room temperature for 6 h before the addition of a solution of **Int.4** (139 mg, 0.1 mmol) in anhydrous ACN (3 mL). The reaction mixture was then stirred at room temperature overnight. Remove the solvent by rotavapor. Then it was diluted by EtOAc (20 mL), washed with 1 M HCl (20 mL), $H_2O$ (20 mL) and saturated brine solution (20 mL). The organic layer was separated, dried over anhydrous $Na_2SO_4$, and concentrated *in vacuo*. The residue was purified with flash chromatography (0~10 % MeOH in DCM as eluent) to yield

**MPD7e** as white solid (160 mg, 58%). $^{1}$H NMR (400 MHz, $CDCl_3$) δ 10.76 (s, 0.5H), 10.58 (s, 0.5H), 7.64 (ddd, *J* = 8.4, 7.3, 5.3 Hz, 1H), 7.42 (dd, *J* = 9.8, 7.2 Hz, 1H), 7.29 – 7.09 (m, 2H), 6.23 (s, 0.5H), 6.08 (s, 0.5H), 5.61 (dd, *J* = 13.1, 9.8 Hz, 1H), 5.17 – 5.01 (m, 1H), 4.96 (s, 0.5H), 4.84 (s, 0.5H), 4.35 – 4.24 (m, 1.5H), 4.23 – 4.11 (m, 2.5H), 4.10 – 3.97 (m, 2H), 3.93 – 3.84 (m, 1H), 3.68 – 3.57 (m, 2H), 3.54 – 3.33 (m, 3H), 3.33 – 3.08 (m, 3H), 3.03 – 2.73 (m, 3H), 2.59 – 2.46 (m, 1H), 2.41 – 2.28 (m, 1H), 2.16 – 1.95 (m, 2H), 1.91 – 1.65 (m, 2H), 1.64 – 1.44 (m, 8H), 1.26 – 1.09 (m, 5H), 1.06 – 0.81 (m, 17H).

**Synthesis of (1R,2S,5S)-3-((2S)-2-(3-(1-((6-((2-(2,6-dioxopiperidin-3-yl)-1,3-dioxoisoindolin-4-yl)oxy)hexyl)oxy)-2-methylpropan-2-yl)ureido)-3,3-dimethylbutanoyl)-6,6-dimethyl-N-((S)-1-oxo-3-((S)-2-oxopyrrolidin-3-yl)propan-2-yl)-3-azabicyclo[3.1.0]hexane-2-carboxamide (MPD7): MPD7**was prepared as a white solid following a similar procedure to **MPD1** (yield 67%).$^{1}$H NMR (400 MHz, $CDCl_3$) δ 10.75 (d, *J* = 36.4 Hz, 0.5H), 10.48 (s, 0.5H), 9.54 (s, 0.5H), 9.52 (s, 0.5H), 8.23 – 8.13 (m, 0.5H), 7.96 – 7.80 (m, 0.5H), 7.70 – 7.50 (m, 2H), 7.35 (ddd, *J* = 12.3, 7.2, 1.6 Hz, 1H), 7.13 (d, *J* = 8.5 Hz, 1H), 6.54 – 6.25 (m, 1H), 5.61 – 5.51 (m, 1H), 5.09 – 4.78 (m, 2H), 4.50 – 4.41 (m, 1H), 4.32 – 4.19 (m, 2H), 4.17 – 3.91 (m, 4H), 3.89 – 3.79 (m, 1H), 3.49 – 3.32 (m, 3H), 3.27 – 3.06 (m, 3H), 2.93 – 2.67 (m, 3H), 2.50 (pd, *J* = 9.2, 4.1 Hz, 1H), 2.32 – 2.16 (m, 1H), 2.03 – 1.92 (m, 2H), 1.90 – 1.62 (m, 4H), 1.61 – 1.27 (m, 9H), 1.20 – 1.05 (m, 6H), 1.03 – 0.90 (m, 6H), 0.86 – 0.78 (m, 9H). ESI-HRMS m/z calculated for $C_{45}H_{63}N_7O_{11}^+$ $(M + H)^+$ 878.4619, found 878.464.

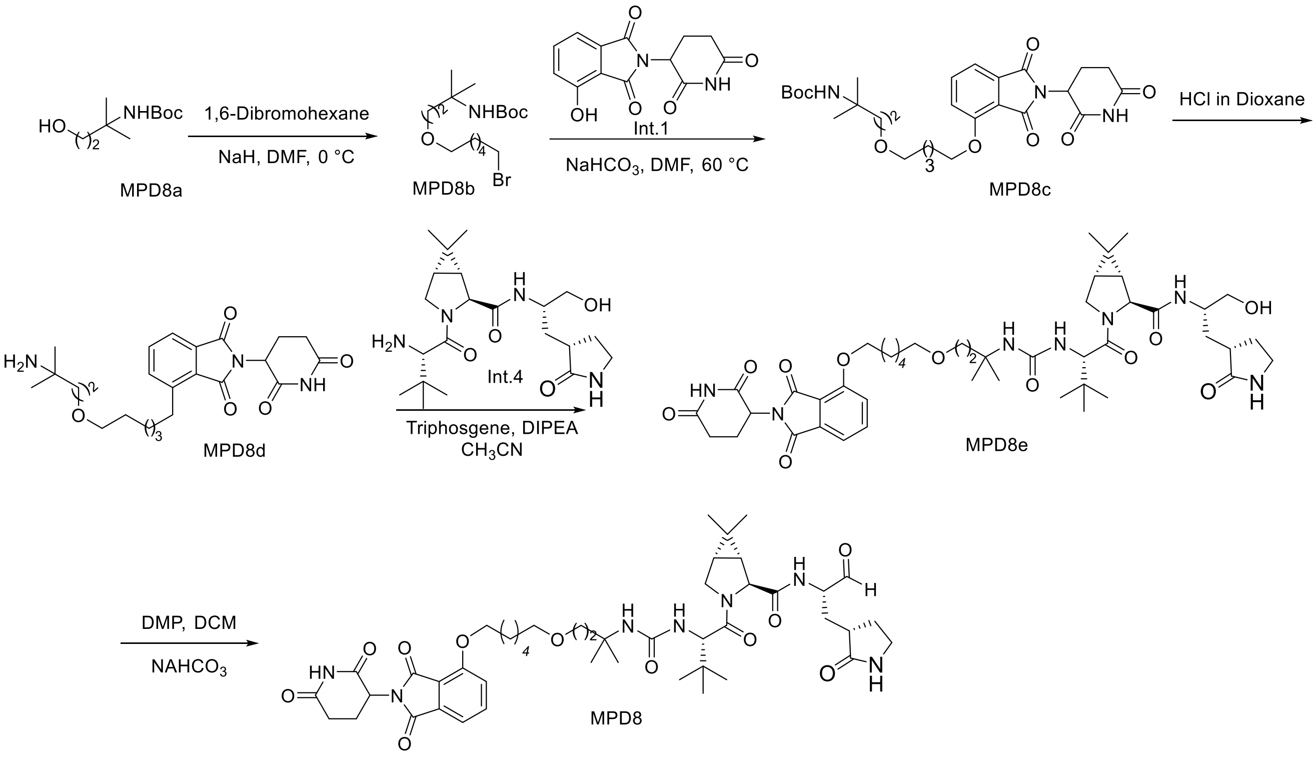


**Scheme 8:** The synthesis of compound **MPD8**

**Synthesis of tert-butyl (4-((6-bromohexyl)oxy)-2-methylbutan-2-yl)carbamate (MPD8b): MPD8b** was prepared as a white solid following a similar procedure to **MPD7b** (yield 77%). $^{1}$H NMR (400 MHz, Chloroform-*d*) δ 5.29 (s, 1H), 3.52 (t, *J* = 6.0 Hz, 2H), 3.40 (td, *J* = 6.6, 2.8 Hz, 4H), 1.86 (p, *J* = 6.8 Hz, 2H), 1.81 (t, *J* = 6.0 Hz, 2H), 1.58 (p, *J* = 6.6 Hz, 2H), 1.48 – 1.36 (m, 13H), 1.32 (s, 6H).

**Synthesis of tert-butyl (4-((6-((2-(2,6-dioxopiperidin-3-yl)-1,3-dioxoisoindolin-4-yl)oxy)hexyl)oxy)-2-methylbutan-2-yl)carbamate (MPD8c): MPD8c** was prepared as a white solid following a similar procedure to **MPD1g** (yield 85%).$^{1}$H NMR (400 MHz, Chloroform-*d*) δ 8.04 (s, 1H), 7.67 (dd, *J* = 8.5, 7.3 Hz, 1H), 7.45 (d, *J* = 7.2 Hz, 1H), 7.21 (d, *J* = 8.4 Hz, 1H), 5.28 (s, 1H), 5.00 – 4.89 (m, 1H), 4.18 (t, *J* = 6.5 Hz, 2H), 3.52 (t, *J* = 6.1 Hz, 2H), 3.41 (t, *J* = 6.4 Hz, 2H), 2.93 – 2.66 (m, 3H), 2.18 – 2.09 (m, 1H), 1.89 (p, *J* = 6.7 Hz, 2H), 1.81 (t, *J* = 6.1 Hz, 2H), 1.65 – 1.46 (m, 6H), 1.42 (s, 9H), 1.32 (s, 6H).

**Synthesis of (1R,2S,5S)-3-((2S)-2-(3-(4-((6-((2-(2,6-dioxopiperidin-3-yl)-1,3-dioxoisoindolin-4-yl)oxy)hexyl)oxy)-2-methylbutan-2-yl)ureido)-3,3-dimethylbutanoyl)-N-((S)-1-hydroxy-3-((S)-2-oxopyrrolidin-3-yl)propan-2-yl)-6,6-dimethyl-3-azabicyclo[3.1.0]hexane-2-carboxamide (MPD8e): MPD8e** was prepared as a white solid following a similar procedure to **MPD7d** and **MPD7e** (yield 74%).$^{1}$H NMR (400 MHz, Chloroform-*d*) δ 10.75 (d, *J* = 13.9 Hz, 1H), 7.66 (ddd, *J* = 8.6, 7.3, 1.5 Hz, 1H), 7.45 (d, *J* = 7.2 Hz, 1H), 7.21 (dd, *J* = 8.5, 3.0 Hz, 1H), 7.13 (dd, *J* = 22.8, 8.4 Hz, 1H), 5.71 (d, *J* = 10.2 Hz, 1H), 5.47 (s, 1H), 5.12 (td, *J* = 12.3, 4.8 Hz, 1H), 4.81 (s, 1H), 4.30 (s, 1H), 4.25 – 4.15 (m, 2H), 4.11 – 3.96 (m, 2H), 3.89 (ddd, *J* = 10.5, 5.3, 3.7 Hz, 1H), 3.63 (dddd, *J* = 30.5, 19.0, 11.2, 4.9 Hz, 2H), 3.44 – 3.16 (m, 6H), 3.05 – 2.77 (m, 3H), 2.61 – 2.48 (m, 1H), 2.42 – 2.30 (m, 1H), 2.16 – 1.98 (m, 3H), 1.93 – 1.68 (m, 6H), 1.64 – 1.41 (m, 9H), 1.24 (d, *J* = 6.2 Hz, 3H), 1.15 (d, *J* = 4.9 Hz, 3H), 1.08 (s, 1H), 1.02 (d, *J* = 7.5 Hz, 3H), 0.97 – 0.87 (m, 11H).

**Synthesis of (1R,2S,5S)-3-((2S)-2-(3-(4-((6-((2-(2,6-dioxopiperidin-3-yl)-1,3-dioxoisoindolin-4-yl)oxy)hexyl)oxy)-2-methylbutan-2-yl)ureido)-3,3-dimethylbutanoyl)-6,6-dimethyl-N-((S)-1-oxo-3-((S)-2-oxopyrrolidin-3-yl)propan-2-yl)-3-azabicyclo[3.1.0]hexane-2-carboxamide (MPD8): MPD8** was prepared as a white solid following a similar procedure to **MPD1** (yield 80%).$^{1}$H NMR (400 MHz, Chloroform-*d*) δ 10.74 (s, 0.41H, diastereomer 1), 10.66 (s, 0.48H, diastereomer 2), 9.63 – 9.56 (m, 0.42H, diastereomer 1), 9.56 – 9.50 (m, 0.51H, diastereomer 2), 7.69 – 7.62 (m, 1H), 7.62 – 7.49 (m, 1H), 7.46 – 7.38 (m, 1H), 7.23 – 7.16 (m, 1H), 6.29 – 6.02 (m, 1H), 5.46 (dd, *J* = 10.1, 6.1 Hz, 1H), 5.13 – 5.01 (m, 1H), 4.92 – 4.82 (m, 1H), 4.56 – 4.44 (m, 1H), 4.36 – 4.26 (m, 2H), 4.22 – 4.12 (m, 2H), 4.01 (t, *J* = 10.4 Hz, 1H), 3.89 (dd, *J* = 10.4, 5.4 Hz, 1H), 3.42 – 3.32 (m, 4H), 3.32 – 3.14 (m, 2H), 3.05 – 2.72 (m, 3H), 2.60 – 2.49 (m, 1H), 2.37 – 2.26 (m, 1H), 2.13 – 1.97 (m, 3H), 1.93 – 1.70 (m, 6H), 1.53 – 1.40 (m, 4H), 1.28 – 1.10 (m, 8H), 1.07 – 1.00 (m, 3H), 0.98 – 0.81 (m, 13H). ESI-HRMS m/z calculated for $C_{46}H_{66}N_{7}O_{11}$ (M+H$^{+}$): 892.4815; found: 892.4798.

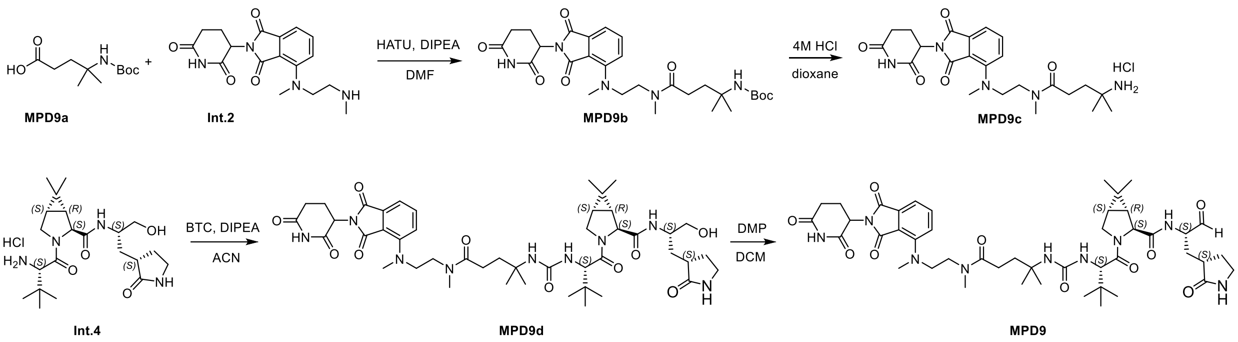


**Scheme 9:** The synthesis of compound **MPD9**

**Synthesis of Tert-butyl (5-((2-((2-(2,6-dioxopiperidin-3-yl)-1,3-dioxoisoindolin-4-yl)(methyl)amino)ethyl)(methyl)amino)-2-methyl-5-oxopentan-2-yl)carbamate (MPD9b).** **MPD9b** was prepared as a yellow oil following a similar procedure to **MPD5c**. (yield 61%). $^{1}$H NMR (400 MHz, Chloroform-*d*) δ 7.5 – 7.4 (m, 1H), 7.2 (d, *J* = 6.0 Hz, 1H), 7.1 (d, *J* = 8.6 Hz, 1H), 4.9 (dd, *J* = 12.1, 5.3 Hz, 1H), 3.6 (dddt, *J* = 82.1, 55.6, 13.8, 7.3 Hz, 2H), 3.0 (s, 1H), 2.8 (d, *J* = 20.5 Hz, 10H), 2.3 – 2.2 (m, 1H), 2.1 – 2.0 (m, 1H), 1.9 (ddd, *J* = 19.4, 13.0, 9.9 Hz, 2H), 1.7 (t, *J* = 7.8 Hz, 1H), 1.4 (s, 9H), 1.2 (s, 3H), 1.2 (s, 3H). $^{13}$C NMR (100 MHz, Chloroform-*d*) δ 177.8, 175.3, 171.7, 168.9, 167.5, 167.1, 149.7, 135.2, 134.2, 123.6, 114.2, 113.6, 77.4, 52.8, 52.0, 49.3, 45.5, 40.1, 38.7, 35.7, 31.5, 29.4, 28.5, 22.7, 20.9.

**Synthesis of 4-amino-N-(2-((2-(2,6-dioxopiperidin-3-yl)-1,3-dioxoisoindolin-4-yl)(methyl)amino)ethyl)-N,4-dimethylpentanamide hydrochloride (MPD9c).** To a solution of MPD9b in dioxane was added 4.0 M HCl in dioxane. The resulting solution was stirred at RT overnight. After the reaction was completed, remove the solvent in vacuo. The residue was used in the next without further purification.

**Synthesis of (1R,2S,5S)-3-((13S)-13-(tert-butyl)-2-(2-(2,6-dioxopiperidin-3-yl)-1,3-dioxoisoindolin-4-yl)-5,9,9-trimethyl-6,11-dioxo-2,5,10,12-tetraazatetradecan-14-oyl)-N-((S)-1-hydroxy-3-((S)-2-oxopyrrolidin-3-yl)propan-2-yl)-6,6-dimethyl-3-azabicyclo[3.1.0]hexane-2-carboxamide (MPD9d).** **MPD9d** was prepared as a white solid following a similar procedure to **MPD7e** (yield 48%). $^{1}$H NMR (400 MHz, Methanol-$d_4$) δ 7.7 – 7.6 (m, 1H), 7.4 – 7.2 (m, 2H), 5.2 – 5.1 (m, 1H), 4.3 – 4.2 (m, 4H), 4.1 – 4.0 (m, 5H), 3.9 (td, *J* = 6.2, 2.4 Hz, 1H), 3.7 (d, *J* = 9.3 Hz, 2H), 3.6 (d, *J* = 5.7 Hz, 4H), 3.3 (dt, *J* = 3.2, 1.6 Hz, 3H), 3.1

(s, 3H), 2.9 (d, *J* = 1.4 Hz, 3H), 2.8 – 2.7 (m, 2H), 2.7 – 2.6 (m, 2H), 2.4 – 2.3 (m, 2H), 2.3 (td, *J* = 6.6, 3.0 Hz, 1H), 2.2 (dtd, *J* = 12.9, 5.0, 2.1 Hz, 1H), 2.0 (tdd, *J* = 13.7, 6.5, 3.6 Hz, 4H), 1.8 – 1.7 (m, 3H), 1.6 (ddd, *J* = 6.8, 4.1, 2.4 Hz, 2H), 1.3 – 1.1 (m, 13H). $^{13}$C NMR (100 MHz, Methanol-$d_4$) δ 181.4, 174.5, 173.4, 172.7, 171.8, 170.3, 167.6, 166.9, 158.1, 149.5, 134.9, 134.2, 123.3, 115.7, 113.7, 64.3, 60.8, 57.5, 54.5, 53.5, 51.4, 50.7, 49.1, 49.0, 42.5, 40.1, 39.2, 37.9, 35.3, 34.5, 32.4, 31.0, 30.9, 27.9, 27.7, 26.5, 25.7, 25.2, 22.4, 18.9, 17.4, 16.0, 11.8.

**Synthesis of (1R,2S,5S)-3-((13S)-13-(tert-butyl)-2-(2-(2,6-dioxopiperidin-3-yl)-1,3-dioxoisoindolin-4-yl)-5,9,9-trimethyl-6,11-dioxo-2,5,10,12-tetraazatetradecan-14-oyl)-6,6-dimethyl-N-((S)-1-oxo-3-((S)-2-oxopyrrolidin-3-yl)propan-2-yl)-3-azabicyclo[3.1.0]hexane-2-carboxamide (MPD9).** MPD9 was prepared as a yellow solid following a similar procedure to **MPD1**. (yield 53%). $^{1}$H NMR (400 MHz, $CDCl_3$) δ 10.88 (s, 0.5H), 10.56 (s, 0.5H), 9.57 – 9.44 (m, 1H), 7.65 (d, *J* = 7.3 Hz, 1H), 7.44 (td, *J* = 8.7, 7.1 Hz, 1H), 7.13 – 7.00 (m, 1H), 6.30 (d, *J* = 29.2 Hz, 0.5H), 5.99 (s, 0.5H), 5.61 (d, *J* = 9.4 Hz, 0.5H), 5.52 (d, *J* = 9.8 Hz, 0.5H), 4.96 (ddd, *J* = 23.6, 12.3, 5.5 Hz, 1H), 4.54 – 4.38 (m, 2H), 4.33 – 4.18 (m, 2H), 4.07 – 3.80 (m, 2H), 3.71 – 3.60 (m, 1H), 3.28 – 3.13 (m, 2H), 3.09 – 3.01 (m, 3H), 2.90 – 2.67 (m, 5H), 2.50 (dq, *J* = 13.7, 4.7 Hz, 1H), 2.34 – 2.24 (m, 2H), 2.00 (td, *J* = 11.6, 6.7 Hz, 3H), 1.90 – 1.64 (m, 2H), 1.58 – 1.33 (m, 3H), 1.30 – 0.96 (m, 12H), 0.94 – 0.75 (m, 14H). ESI-HRMS m/z calculated for $C_{45}H_{63}N_9O_{10}^+$ $(M + H)^+$ 890.4731, found 890.4763.

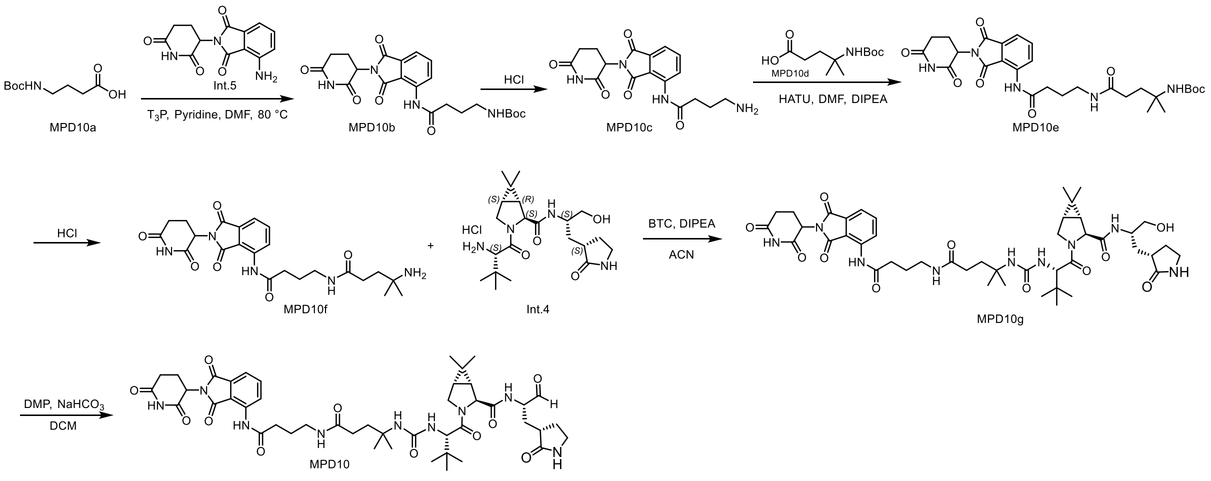


**Scheme 10:** The synthesis of compound **MPD10**

**Synthesis of tert-butyl (4-((2-(2,6-dioxopiperidin-3-yl)-1,3-dioxoisoindolin-4-yl)amino)-4-oxobutyl)carbamate (MPD10b). MPD10b** was prepared as a white solid following a similar procedure to **MPD5e** (yield 62%)$^{1}$H NMR (400 MHz, DMSO-$d_6$) δ 11.15 (s, 1H), 9.70 (s, 1H), 8.52 – 8.42 (m, 1H), 7.90 – 7.80 (m, 1H), 7.67 – 7.56 (m, 1H), 6.87 (t, *J* = 5.8 Hz, 1H), 5.23 – 5.03 (m, 1H), 3.67 – 3.59 (m, 1H), 3.00 (q, *J* = 6.5 Hz, 2H), 2.69 – 2.54 (m, 2H), 2.40 (t, *J* = 8.1 Hz, 1H), 2.14 – 2.01 (m, 1H), 1.95 – 1.84 (m, 2H), 1.82 – 1.64 (m, 2H), 1.45 (s, 6H), 1.38 (s, 8H).

**Synthesis of 4-amino-N-(2-(2,6-dioxopiperidin-3-yl)-1,3-dioxoisoindolin-4-yl)butanamide (MPD10c). MPD10c** was prepared as a white solid following a similar procedure to **MPD9c** (yield 77%)$^{1}$H NMR (400 MHz, DMSO-$d_6$) δ 11.18 (s, 1H), 9.86 (s, 1H), 8.39 (d, *J* = 8.3 Hz, 1H), 8.17 (d, *J* = 7.0 Hz, 2H), 7.88 (t, *J* = 7.9 Hz, 1H), 7.66 (d, *J* = 7.3 Hz, 1H), 3.78 – 3.61 (m, 1H), 3.32 (t, *J* = 7.0 Hz, 1H), 2.71 – 2.46 (m, 4H), 2.28 – 1.92 (m, 5H). $^{13}$C NMR (100 MHz, DMSO-$d_6$) δ 179.31, 173.57, 173.50, 171.68, 170.62, 170.31, 168.99, 167.78, 167.14, 163.32, 146.90, 136.60, 136.39, 136.06, 131.76, 127.32, 119.19, 118.09, 66.71, 60.51, 49.29, 42.51, 36.56, 33.47, 31.43, 31.27, 30.33, 22.89, 22.36, 20.56.

**Synthesis of tert-butyl (5-((4-((2-(2,6-dioxopiperidin-3-yl)-1,3-dioxoisoindolin-4-yl)amino)-4-oxobutyl)amino)-2-methyl-5-oxopentan-2-yl)carbamate (MPD10e). MPD10e** was prepared as a yellow oil following a similar procedure to **MPD5c**. (yield 45%). $^{1}$H NMR (400 MHz, DMSO-$d_6$) δ 11.15 (s, 1H), 9.70 (s, 1H), 8.85 – 8.70 (m, 1H), 8.59 – 8.43 (m, 1H), 7.89 – 7.73 (m, 2H), 7.67 – 7.57 (m, 1H), 7.57 – 7.45 (m, 1H), 7.09 – 6.93 (m, 1H), 5.24 – 4.90 (m, 1H), 3.10 (q, *J* = 6.6 Hz, 2H), 2.67 – 2.52 (m, 2H), 2.30 – 1.98 (m, 4H), 1.90 – 1.64 (m, 5H), 1.37 (d, *J* = 1.7 Hz, 11H), 1.30 – 1.22 (m, 2H), 1.16 (d, *J* = 1.4 Hz, 2H), 1.15 (d, *J* = 0.9 Hz, 6H).

**Synthesis of 4-amino-N-(2-(2,6-dioxopiperidin-3-yl)-1,3-dioxoisoindolin-4-yl)butanamide 4-amino-N-(4-((2-(2,6-dioxopiperidin-3-yl)-1,3-dioxoisoindolin-4-yl)amino)-4-oxobutyl)-4-methylpentanamide (MPD10f)**. **MPD10f** was prepared as a yellow oil following a similar procedure to **MPD9c**. (yield 61%). $^{1}$H NMR (400 MHz, DMSO-*d6*) δ 11.15 (s, 1H), 9.74 (s, 1H), 8.45 (d, *J* = 8.3 Hz, 1H), 8.24 – 8.02 (m, 4H), 7.84 (dd, *J* = 8.4, 7.3 Hz, 1H), 7.62 (dd, *J* = 7.3, 0.8 Hz, 1H), 7.57 – 7.38 (m, 1H), 7.15 – 6.86 (m, 1H), 5.22 – 5.06 (m, 1H), 3.73 – 3.62 (m, 1H), 3.54 – 3.45 (m, 1H), 3.12 (q, *J* = 6.6 Hz, 1H), 2.93 – 2.85 (m, 1H), 2.63 – 2.34 (m, 6H), 2.24 – 1.98 (m, 3H), 1.84 – 1.71 (m, 3H), 1.25 – 1.21 (m, 7H).

**Synthesis of (1R,2S,5S)-3-((2S)-2-(3-(5-((4-((2-(2,6-dioxopiperidin-3-yl)-1,3-dioxoisoindolin-4-yl)amino)-4-oxobutyl)amino)-2-methyl-5-oxopentan-2-yl)ureido)-3,3-dimethylbutanoyl)-N-((S)-1-hydroxy-3-((S)-2-oxopyrrolidin-3-yl)propan-2-yl)-6,6-dimethyl-3-azabicyclo[3.1.0]hexane-2-carboxamide (MPD10g). MPD10g** was prepared as a white solid following a similar procedure to **MPD7e** (yield 56%). $^{1}$H NMR (400 MHz, Methanol-*d*4) δ 8.55 (d, *J* = 8.4 Hz, 1H), 7.68 (dd, *J* = 8.5, 7.3 Hz, 1H), 7.49 (d, *J* = 7.3 Hz, 1H), 5.89 (d, *J* = 2.3 Hz, 1H), 5.76 (d, *J* = 9.5 Hz, 1H), 5.04 (dd, *J* = 12.5, 5.4 Hz, 1H), 4.48 (s, 1H), 4.19 – 4.09 (m, 2H), 3.96 – 3.82 (m, 2H), 3.69 – 3.56 (m, 2H), 3.46 – 3.37 (m, 2H), 3.20 – 3.08 (m, 5H), 2.83 – 2.72 (m, 1H), 2.72 – 2.60 (m, 1H), 2.60 – 2.48 (m, 1H), 2.45 (t, *J* = 7.4 Hz, 1H), 2.31 – 2.21 (m, 1H), 2.11 – 2.01 (m, 2H), 1.86 – 1.79 (m, 3H), 1.68 – 1.60 (m, 1H), 1.45 (dd, *J* = 7.7, 5.1 Hz, 1H), 1.37 (t, *J* = 2.7 Hz, 1H), 1.28 (s, 5H), 1.27 (d, *J* = 2.4 Hz, 6H), 1.16 – 1.11 (m, 3H), 1.10 (s, 2H), 0.92 (d, *J* = 1.6 Hz, 2H), 0.89 (s, 7H), 0.82 (s, 2H).

**Synthesis of (1R,2S,5S)-3-((2S)-2-(3-(5-((4-((2-(2,6-dioxopiperidin-3-yl)-1,3-dioxoisoindolin-4-yl)amino)-4-oxobutyl)amino)-2-methyl-5-oxopentan-2-yl)ureido)-3,3-dimethylbutanoyl)-6,6-dimethyl-N-((S)-1-oxo-3-((S)-2-oxopyrrolidin-3-yl)propan-2-yl)-3-azabicyclo[3.1.0]hexane-2-carboxamide (MPD10): MPD10** was prepared as a white solid following a similar procedure to **MPD1** (yield 48%). $^{1}$H NMR (400 MHz, Chloroform-*d*) δ 10.74 (d, *J* = 58.9 Hz, 1H), 9.56 – 9.33 (m, 1H), 8.76 – 8.60 (m, 1H), 7.80 (dd, *J* = 7.9, 1.2 Hz, 1H), 7.72 – 7.55 (m, 1H), 7.55 – 7.44 (m, 1H), 7.26 (dd, *J* = 7.5, 1.2 Hz, 1H), 6.99 – 6.90 (m, 1H), 6.33 – 6.06 (m, 1H), 5.14 – 4.96 (m, 1H), 4.27 – 4.14 (m, 1H), 4.10 – 3.80 (m, 2H), 3.69 (h, *J* = 6.7 Hz, 1H), 3.64 – 3.48 (m, 1H), 3.44 – 3.30 (m, 1H), 3.30 – 3.11 (m, 2H), 2.87 – 2.70 (m, 2H), 2.64 – 2.38 (m, 2H), 2.38 – 2.20 (m, 1H), 2.20 – 2.02 (m, 2H), 1.99 – 1.83 (m, 2H), 1.86 – 1.58 (m, 3H), 1.42 (dd, *J* = 20.5, 7.1 Hz, 6H), 1.25 – 1.15 (m, 7H), 1.12 – 1.02 (m, 2H), 1.00 – 0.71 (m, 11H). ESI-HRMS m/z calculated for $C_{45}H_{61}N_9O_{11}^+$ $(M + H)^+$ 904.4524, found 904.454.

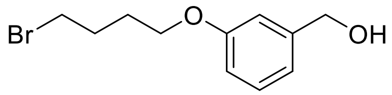
Br
O
OH

MPD1c

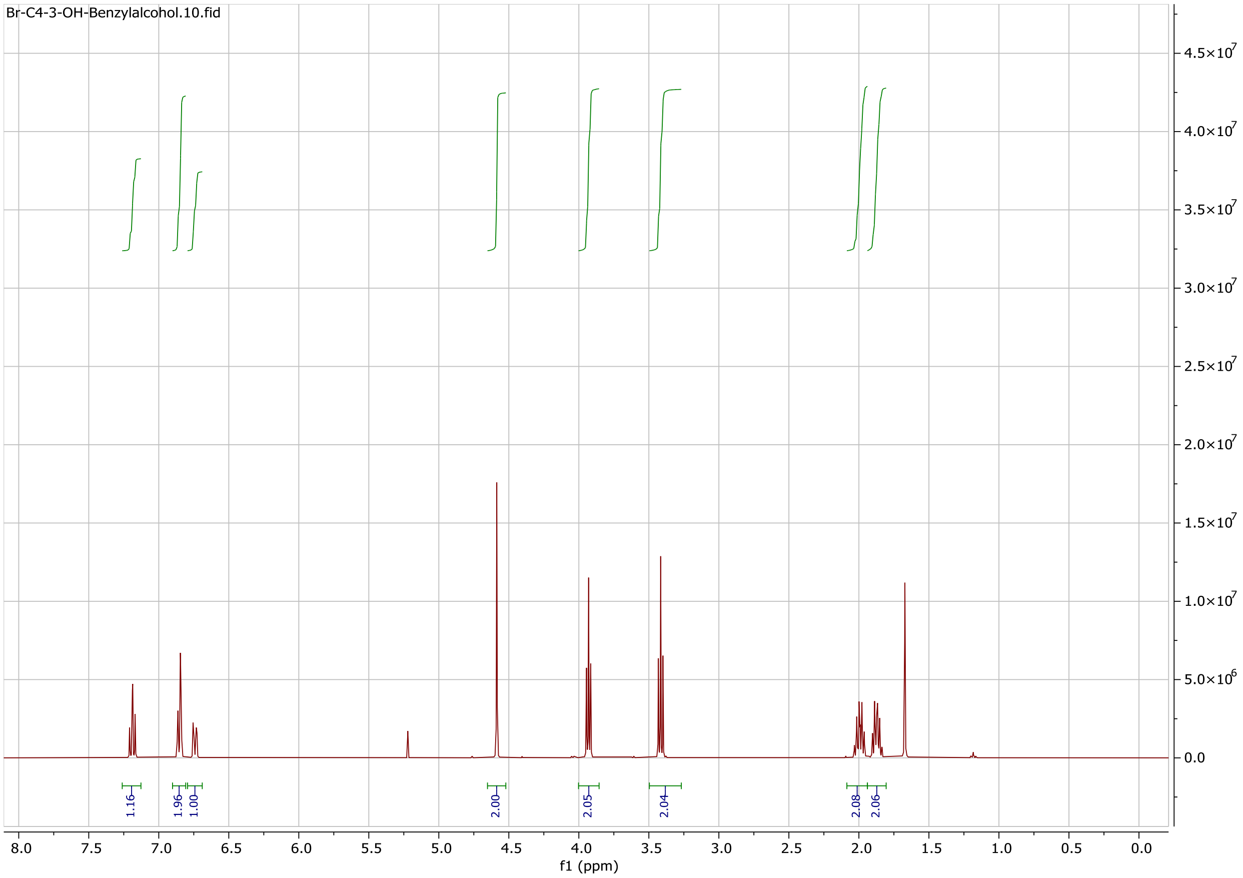
Br-C4-3-OH-Benzylalcohol.10.fid
4.5×10^7
4.0×10^7
3.5×10^7
3.0×10^7
2.5×10^7
2.0×10^7
1.5×10^7
1.0×10^7
5.0×10^6
0.0
1.16
1.96
1.00
2.00
2.05
2.04
2.08
2.06
8.0
7.5
7.0
6.5
6.0
5.5
5.0
4.5
4.0
3.5
3.0
2.5
2.0
1.5
1.0
0.5
0.0
f1 (ppm)

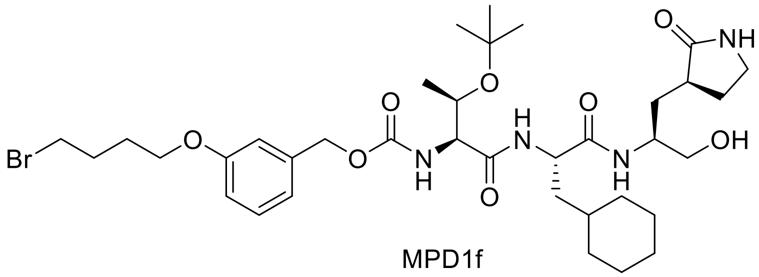

MPD1f

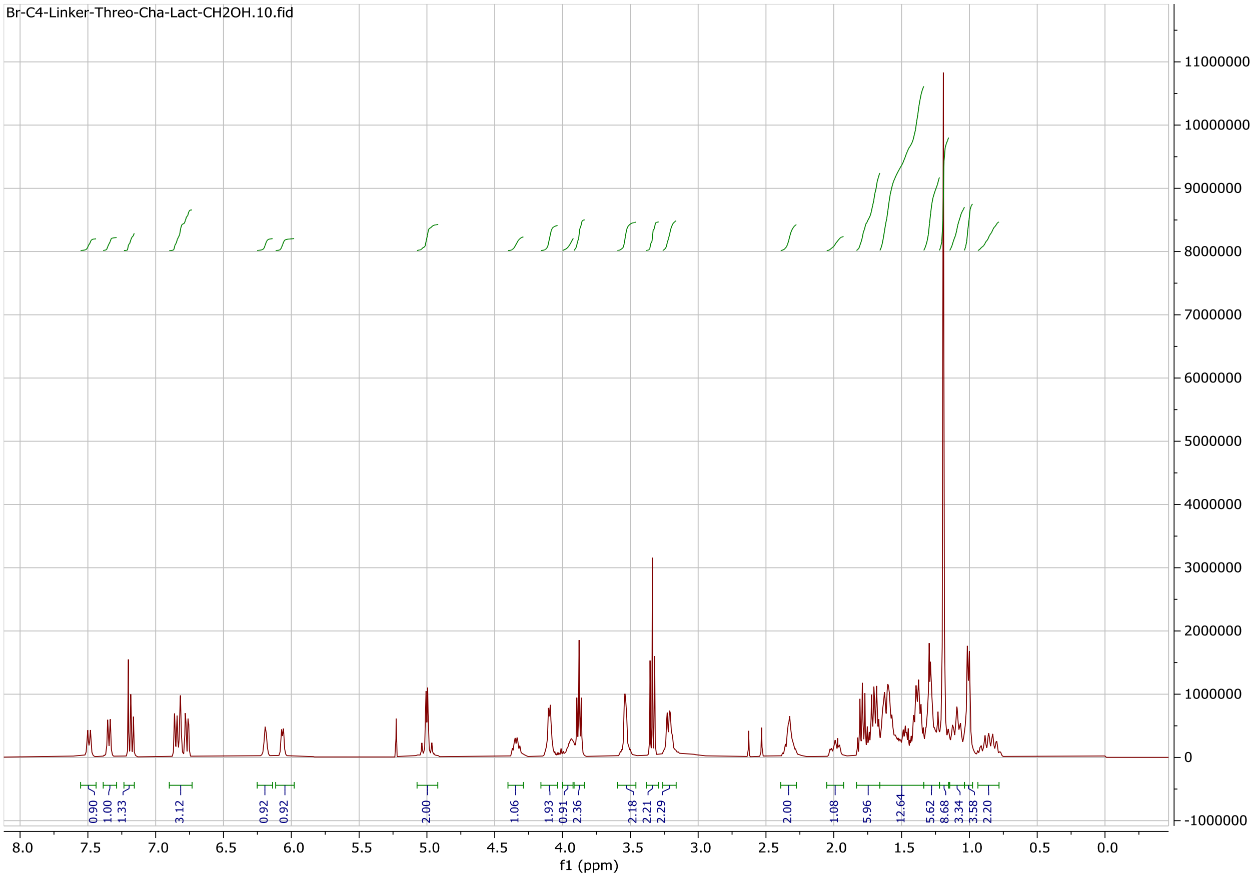

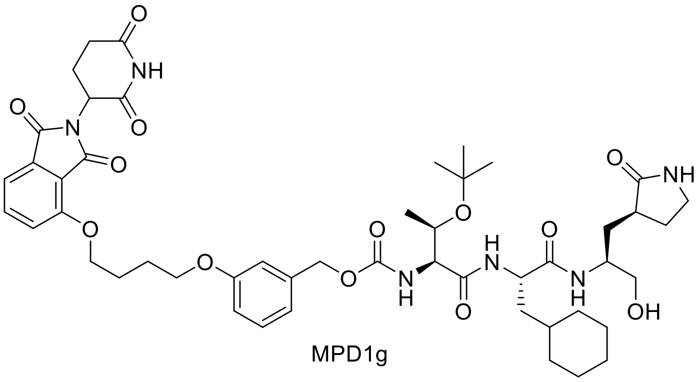

MPD1g

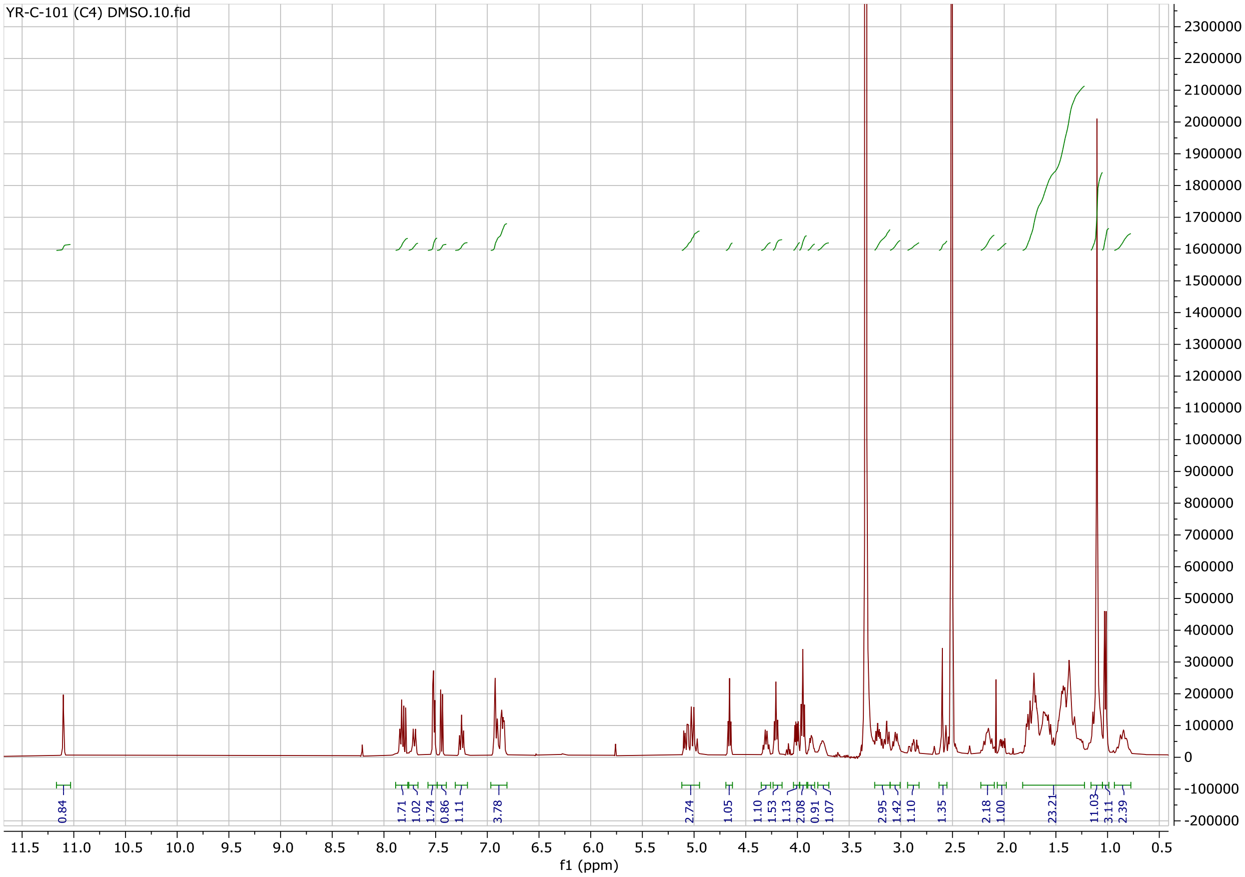

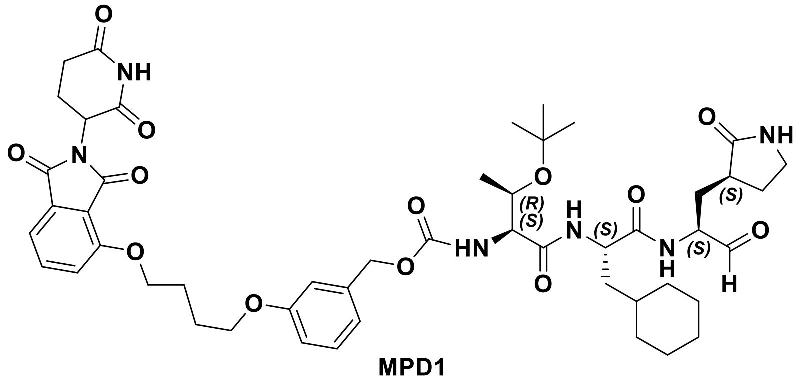

**MPD1**

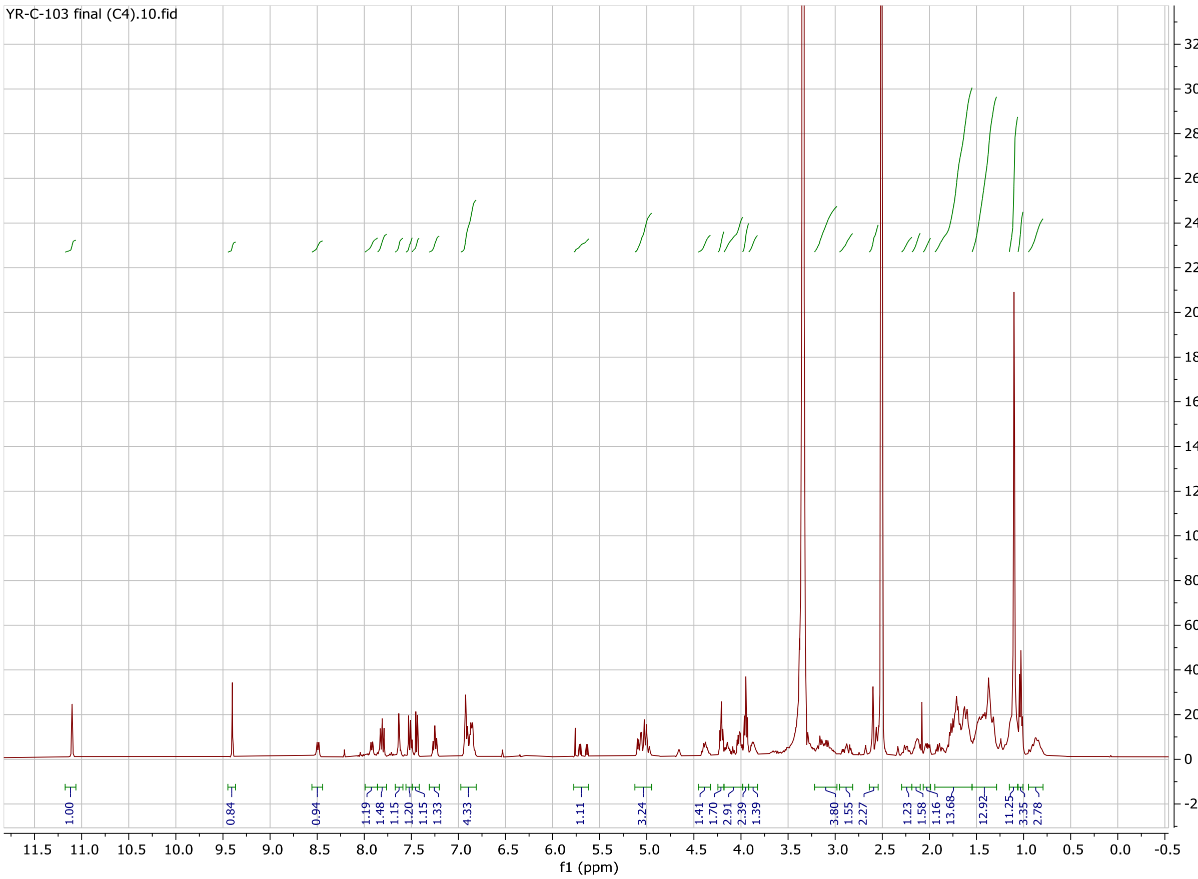

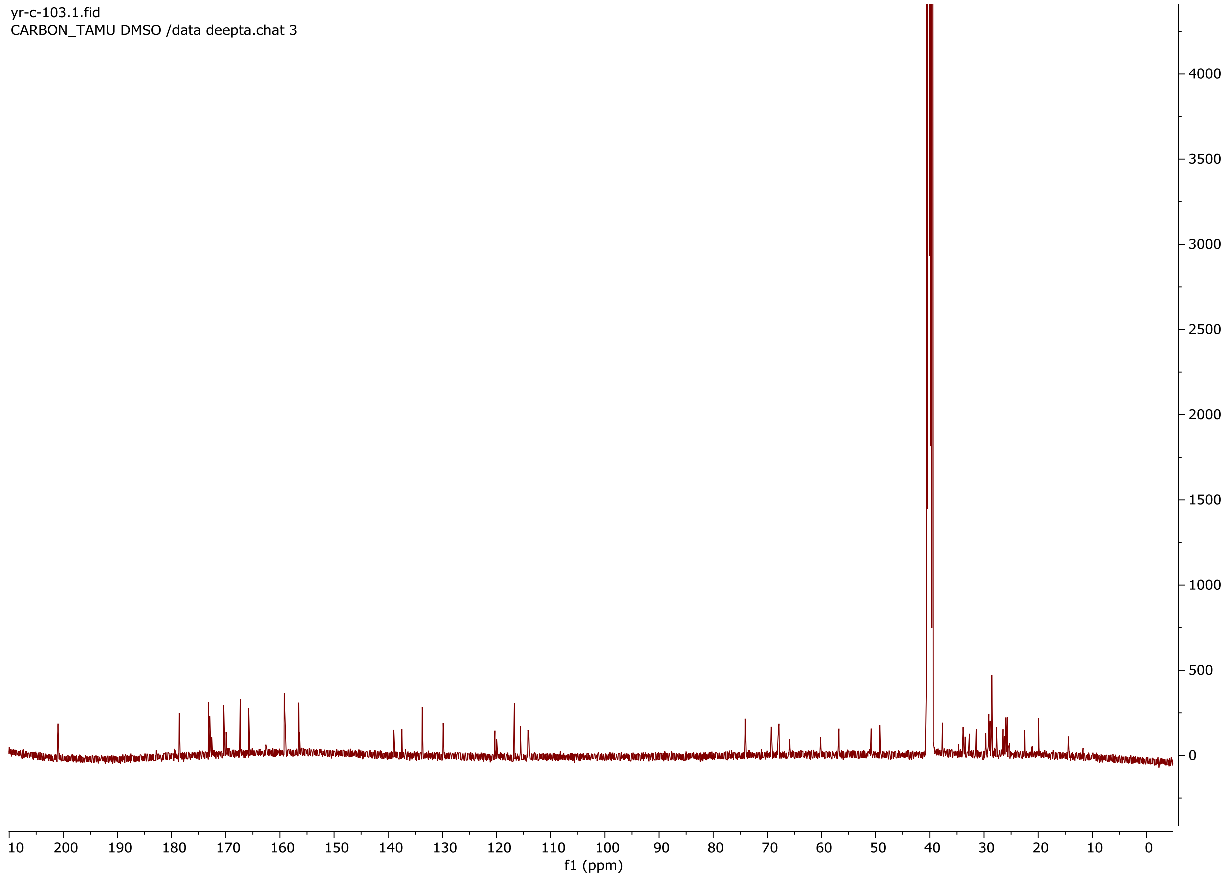
yr-c-103.1.fid
CARBON_TAMU DMSO /data deepta.chat 3
f1 (ppm)

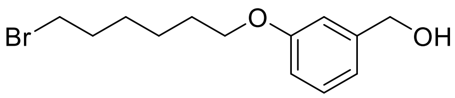


MPD2b

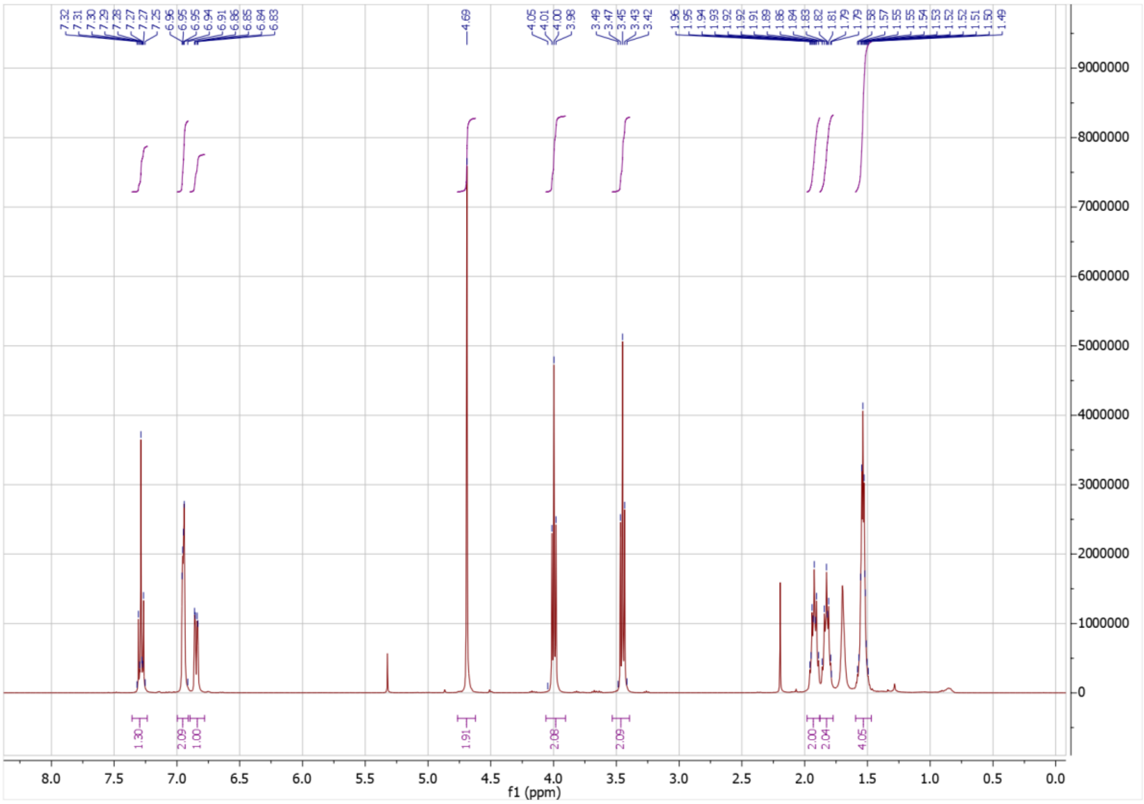

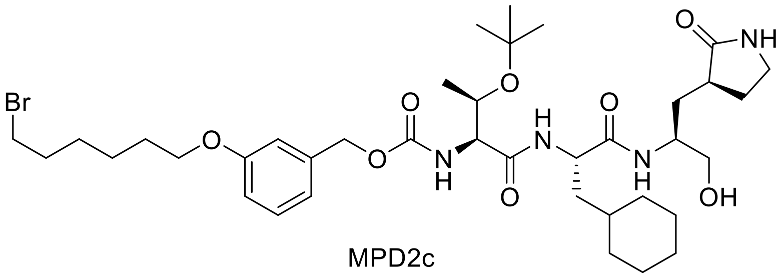

MPD2c

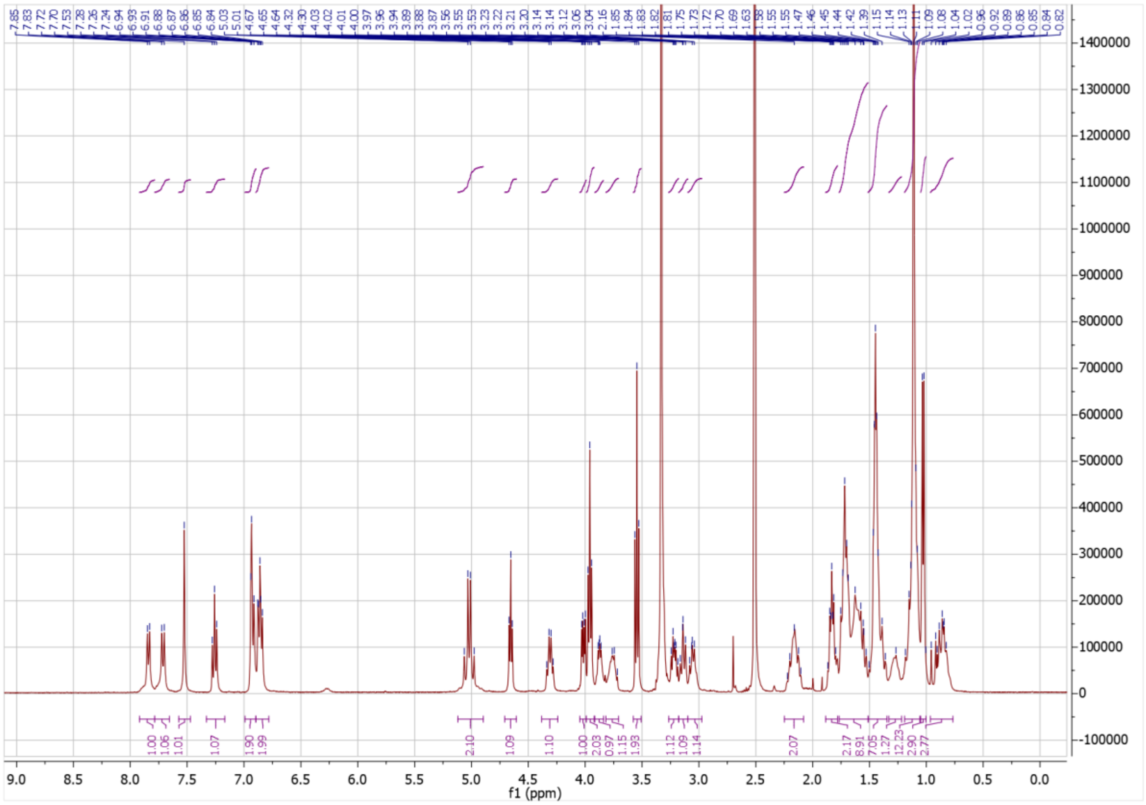

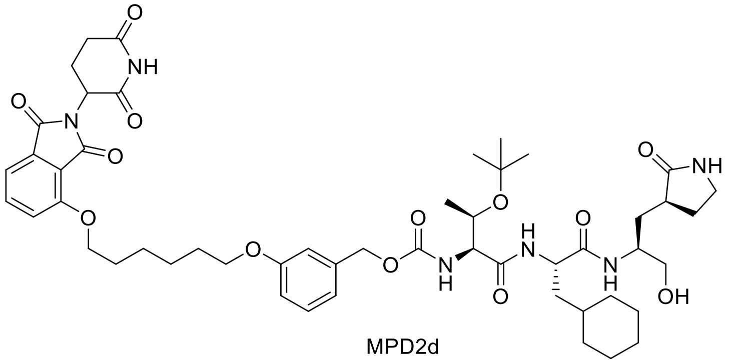

O
NH
O
N
O
O
O
O
O
O
N
H
O
H
N
O
O
NH
N
H
OH


MPD2d

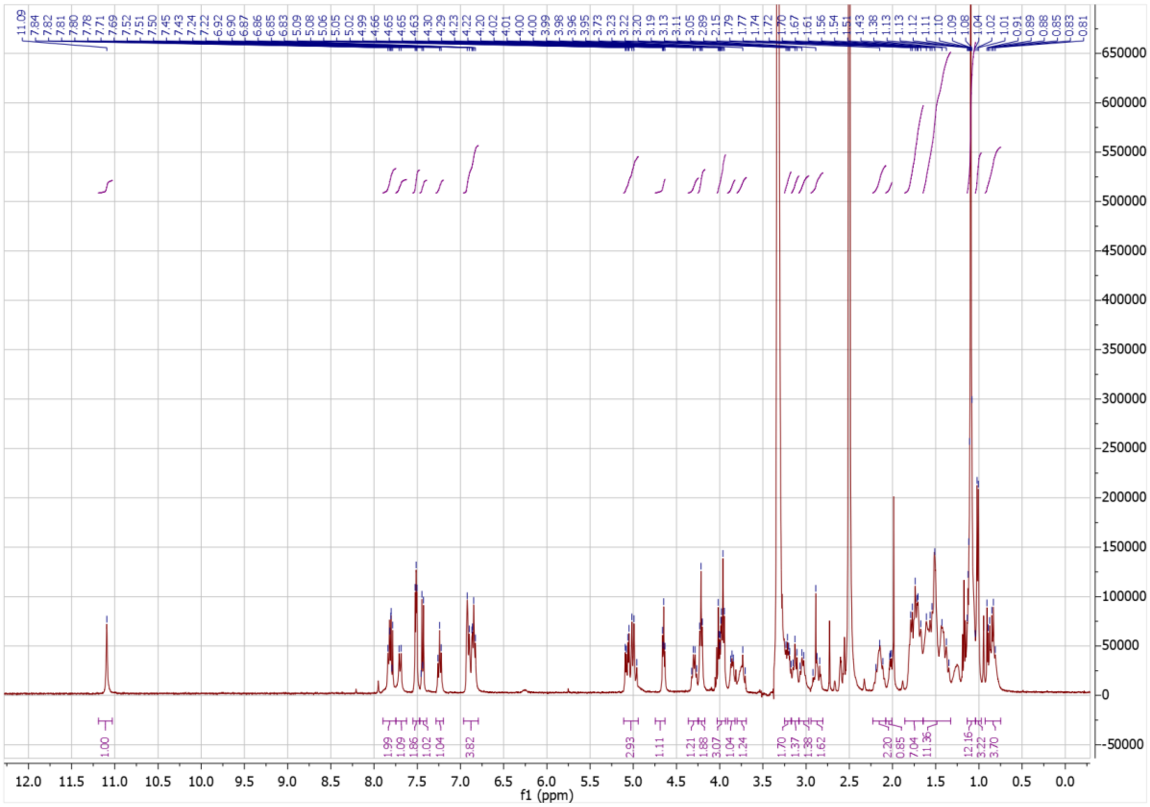

12.0 11.5 11.0 10.5 10.0 9.5 9.0 8.5 8.0 7.5 7.0 6.5 6.0 5.5 5.0 4.5 4.0 3.5 3.0 2.5 2.0 1.5 1.0 0.5 0.0
f1 (ppm)
650000 600000 550000 500000 450000 400000 350000 300000 250000 200000 150000 100000 50000 0 -50000

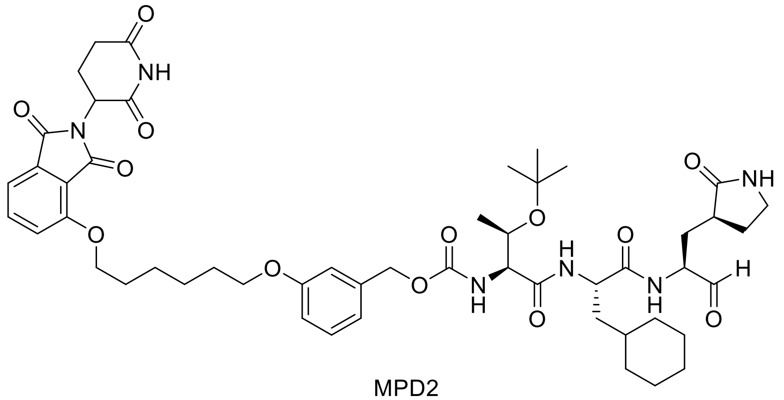

MPD2

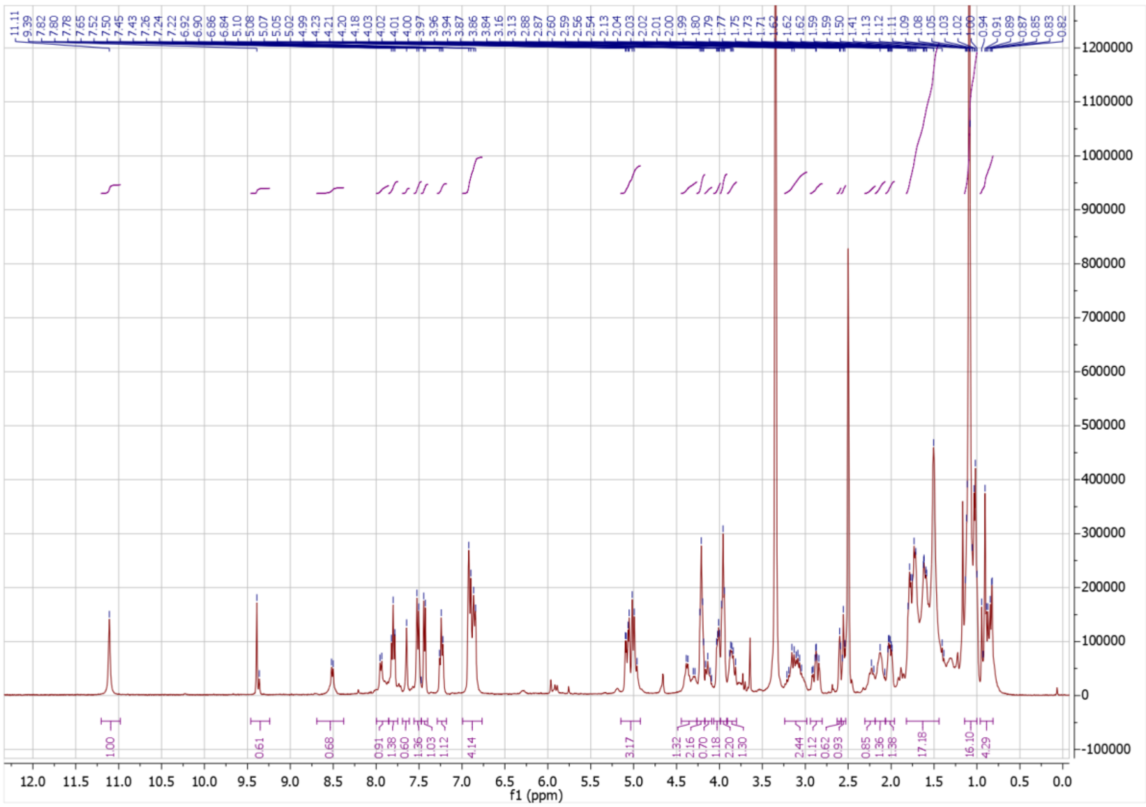

12.0 11.5 11.0 10.5 10.0 9.5 9.0 8.5 8.0 7.5 7.0 6.5 6.0 5.5 5.0 4.5 4.0 3.5 3.0 2.5 2.0 1.5 1.0 0.5 0.0
f1 (ppm)
1200000
1100000
1000000
900000
800000
700000
600000
500000
400000
300000
200000
100000
0
-100000

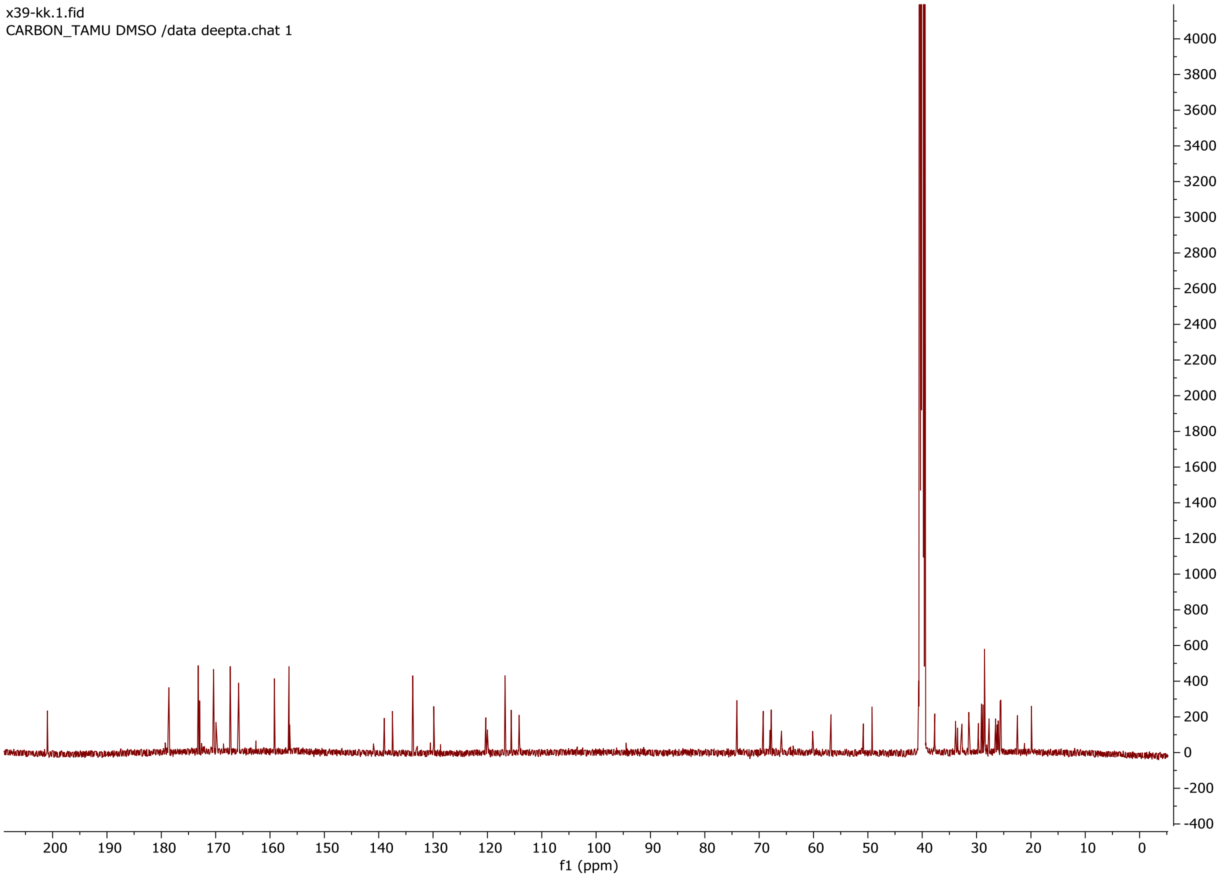
x39-kk.1.fid
CARBON_TAMU DMSO /data deepta.chat 1
f1 (ppm)

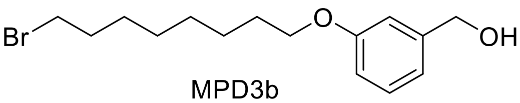

Br
O
OH


MPD3b

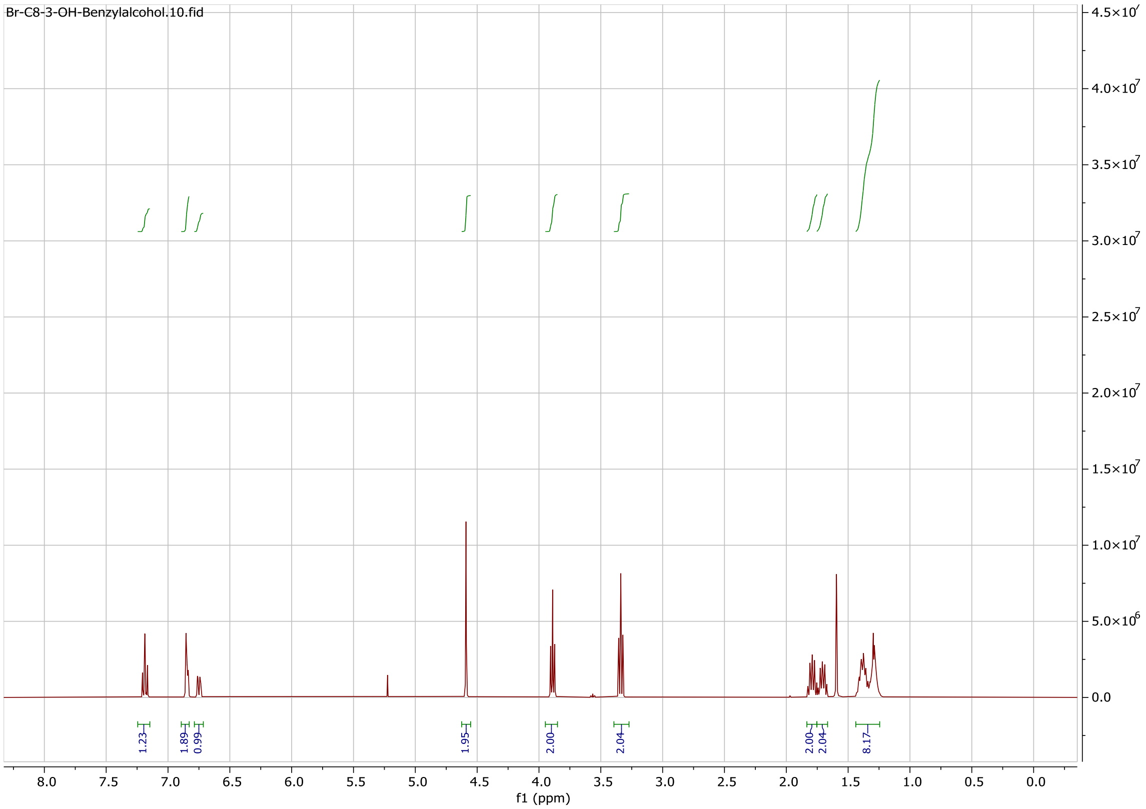

Br-C8-3-OH-Benzylalcohol.10.fid
f1 (ppm)
8.0 7.5 7.0 6.5 6.0 5.5 5.0 4.5 4.0 3.5 3.0 2.5 2.0 1.5 1.0 0.5 0.0
1.23 1.89 0.99 1.95 2.00 2.04 2.00 2.04 8.17
4.5×10^7 4.0×10^7 3.5×10^7 3.0×10^7 2.5×10^7 2.0×10^7 1.5×10^7 1.0×10^7 5.0×10^6 0.0

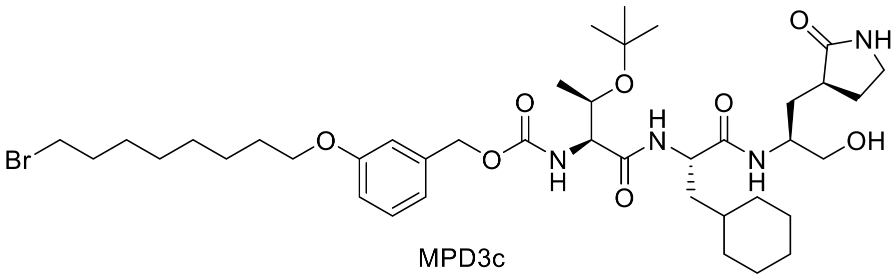

MPD3c

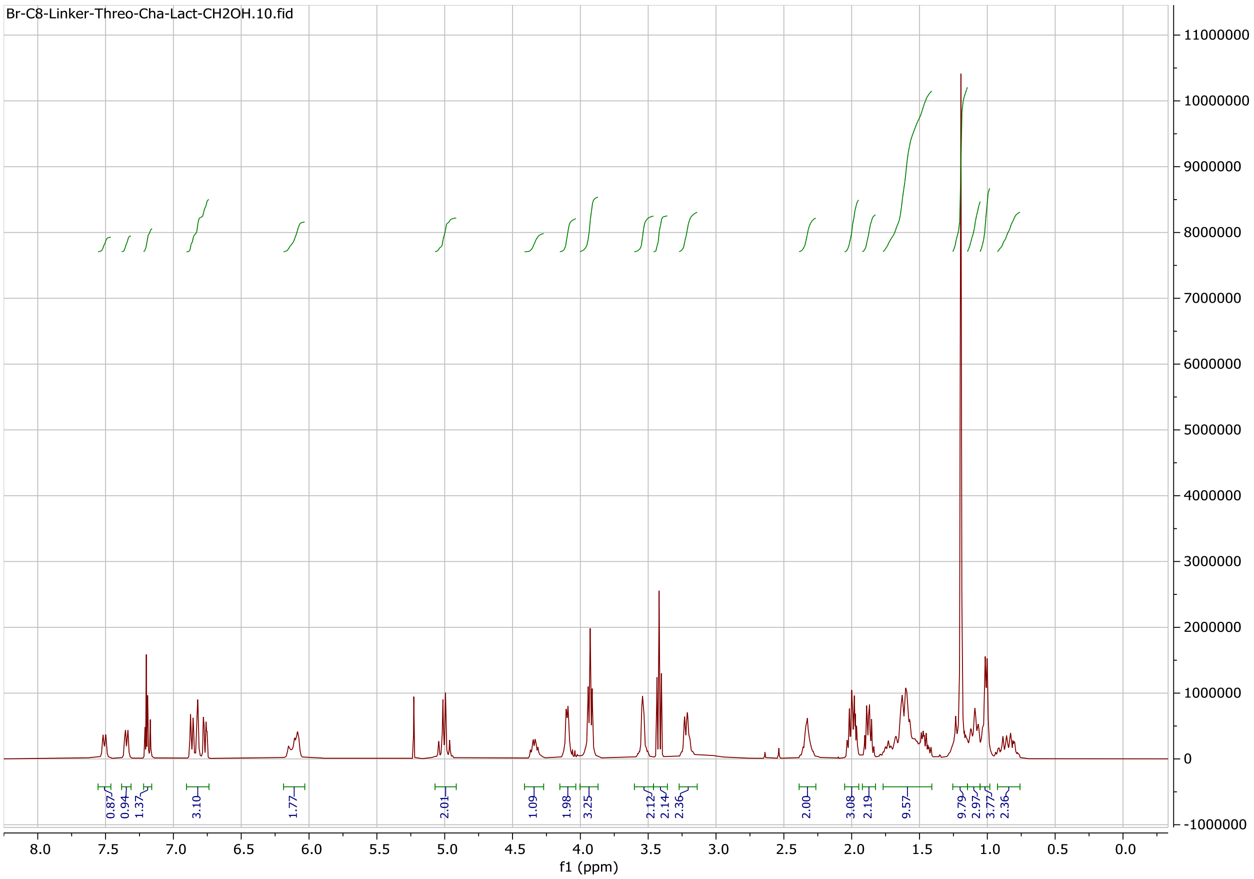

Br-C8-Linker-Threo-Cha-Lact-CH2OH.10.fid
11000000
10000000
9000000
8000000
7000000
6000000
5000000
4000000
3000000
2000000
1000000
0
-1000000
0.87
0.94
1.37
3.10
1.77
2.01
1.09
1.98
3.25
2.12
2.14
2.36
2.00
3.08
2.19
9.57
9.79
2.97
3.77
2.36
8.0 7.5 7.0 6.5 6.0 5.5 5.0 4.5 4.0 3.5 3.0 2.5 2.0 1.5 1.0 0.5 0.0
f1 (ppm)

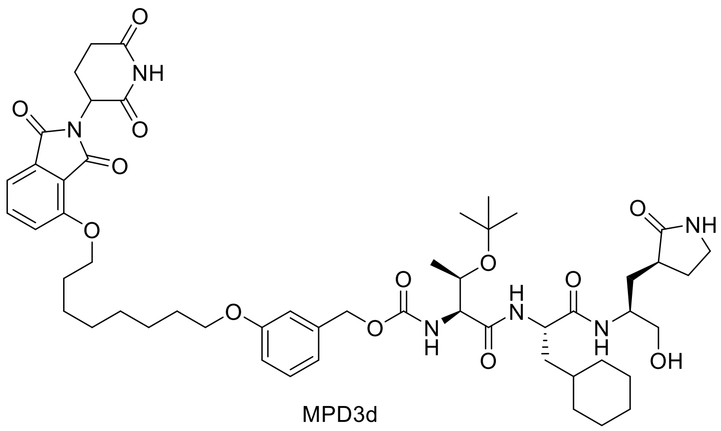

MPD3d

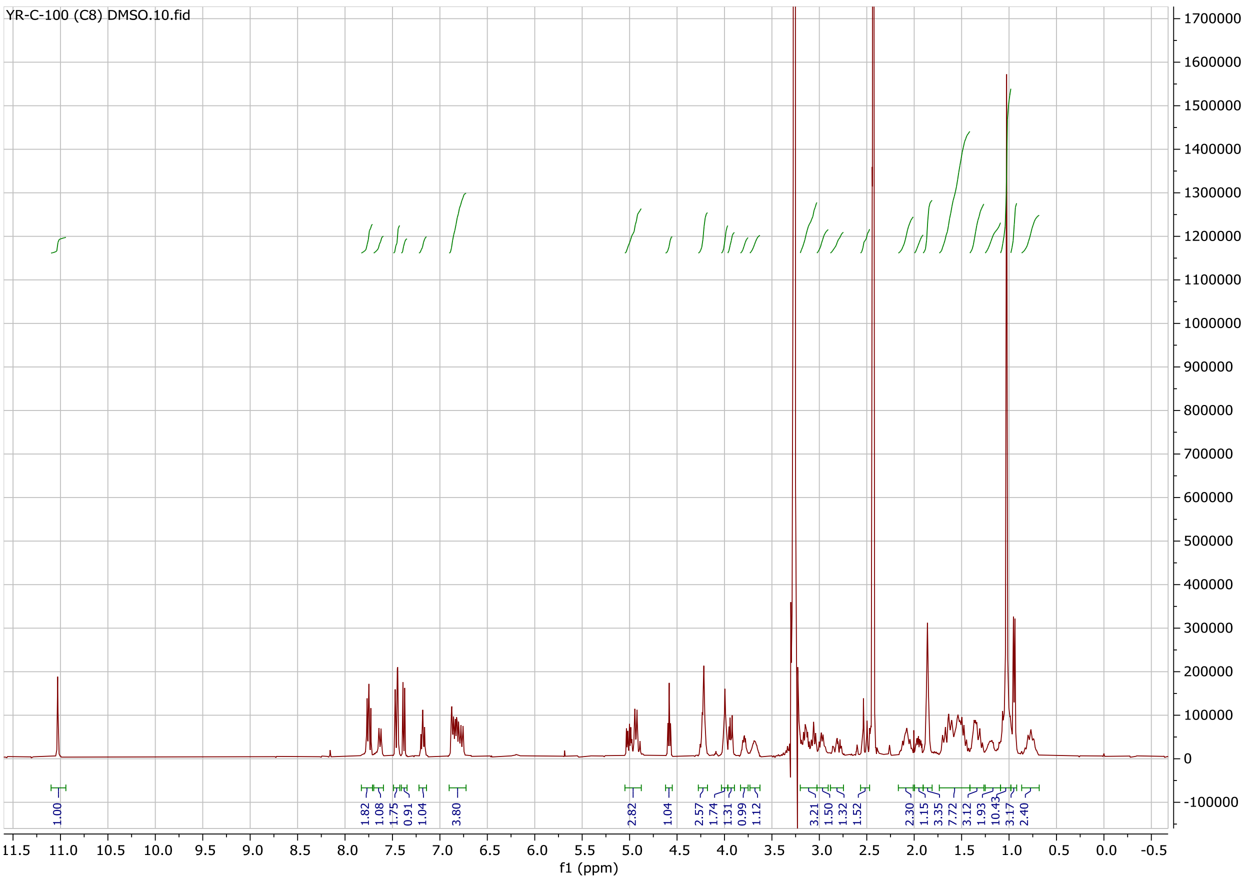

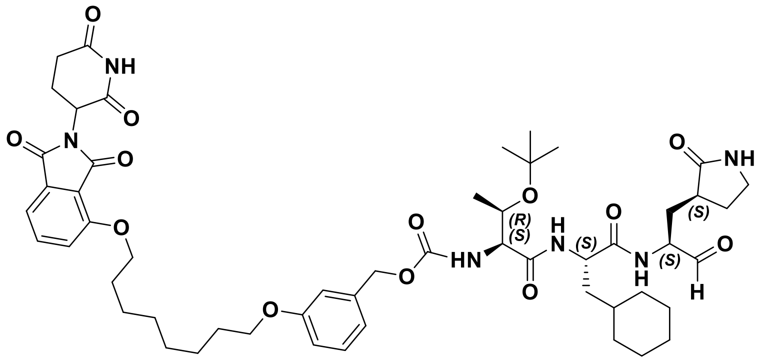
O
NH
O
O
N
O
O
O
O
HN
(R)
(S)
H
N
(S)
O
O
N
H
(S)
O
NH
(S)
O
H

**MPD3**

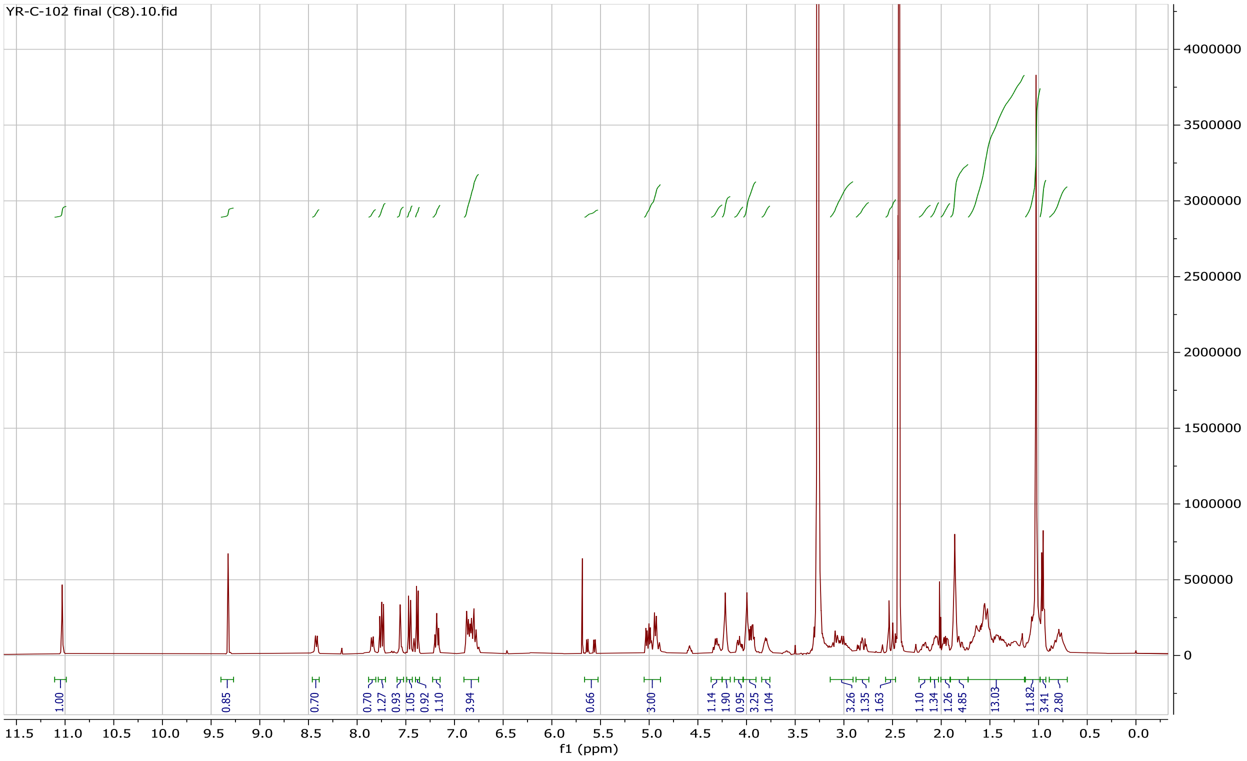
YR-C-102 final (C8).10.fid
1.00
0.85
0.70
0.70
1.27
0.93
1.05
0.92
1.10
3.94
0.66
3.00
1.14
1.90
0.95
3.25
1.04
3.26
1.35
1.63
1.10
1.34
1.26
4.85
13.03
11.82
3.41
2.80
f1 (ppm)

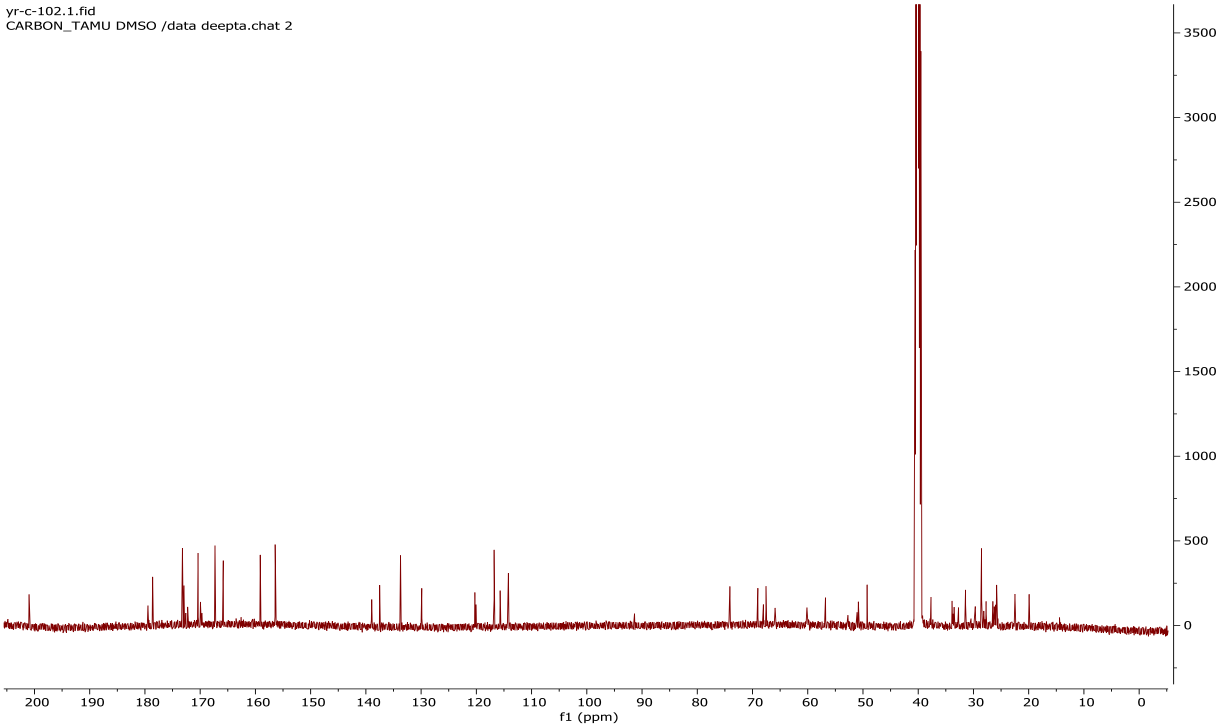
yr-c-102.1.fid
CARBON_TAMU DMSO /data deepta.chat 2
3500
3000
2500
2000
1500
1000
500
0
200 190 180 170 160 150 140 130 120 110 100 90 80 70 60 50 40 30 20 10 0
f1 (ppm)

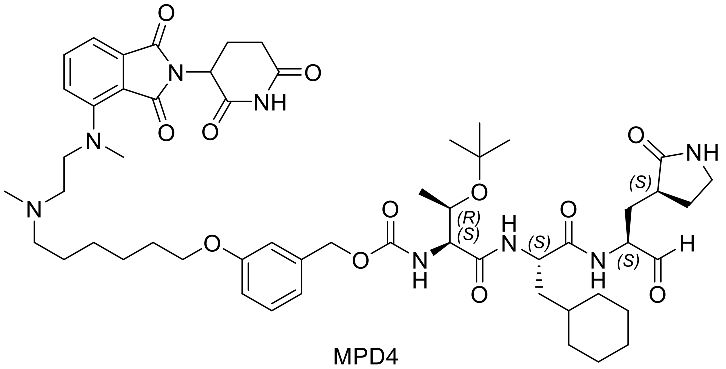

MPD4

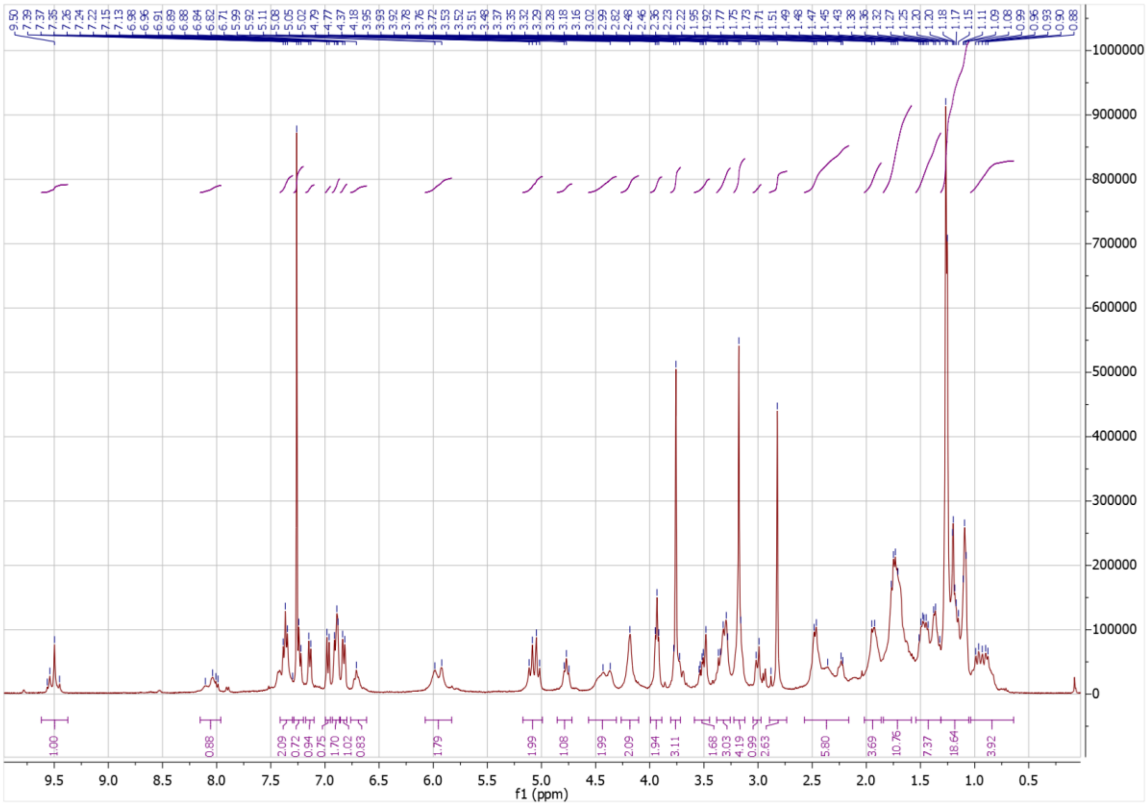
1000000
900000
800000
700000
600000
500000
400000
300000
200000
100000
0
9.5 9.0 8.5 8.0 7.5 7.0 6.5 6.0 5.5 5.0 4.5 4.0 3.5 3.0 2.5 2.0 1.5 1.0 0.5
f1 (ppm)

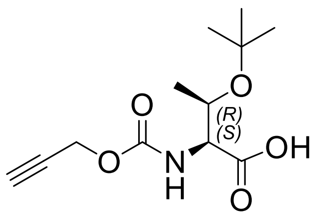

MPD5b

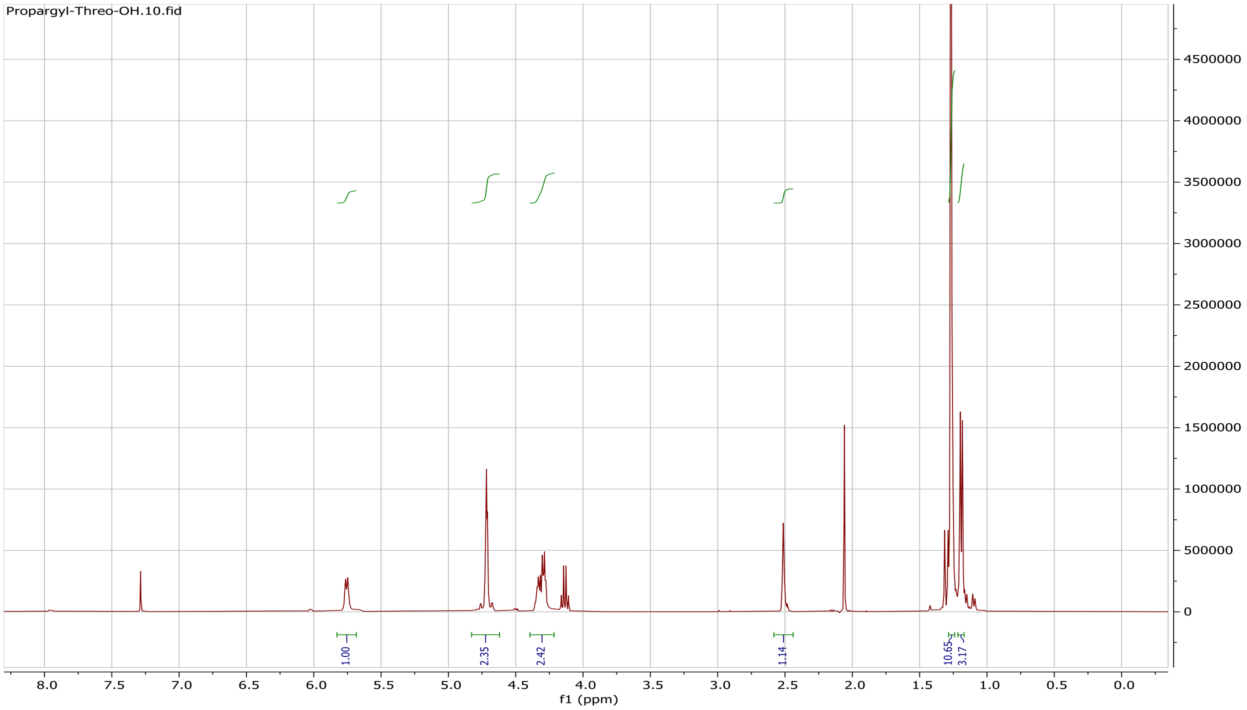

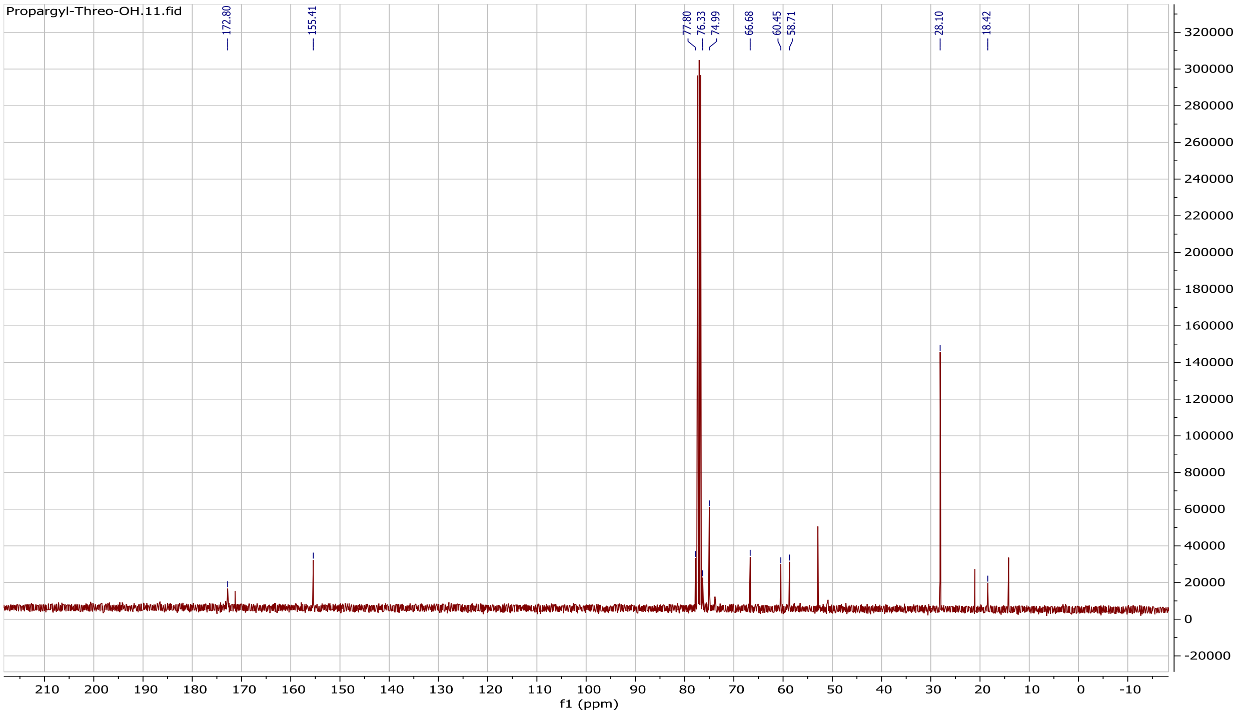

Propargyl-Threo-OH.11.fid
172.80
155.41
77.80
76.33
74.99
66.68
60.45
58.71
28.10
18.42
f1 (ppm)

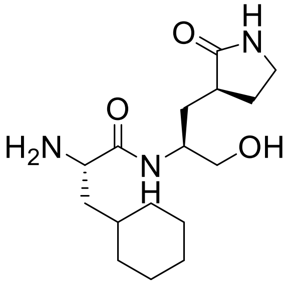

H
O
N
O
H2N
N
H
OH


Int.3

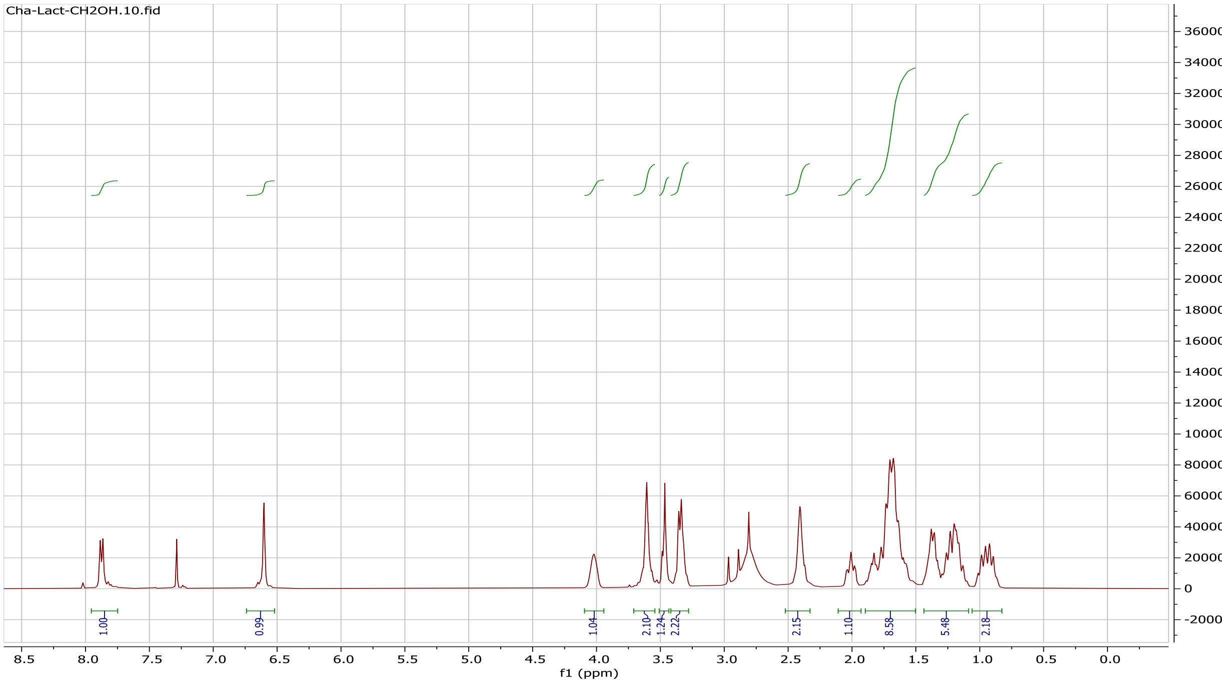

Cha-Lact-CH2OH.10.fid
1.00
0.99
1.04
2.10
1.24
2.22
2.15
1.10
8.58
5.48
2.18
f1 (ppm)

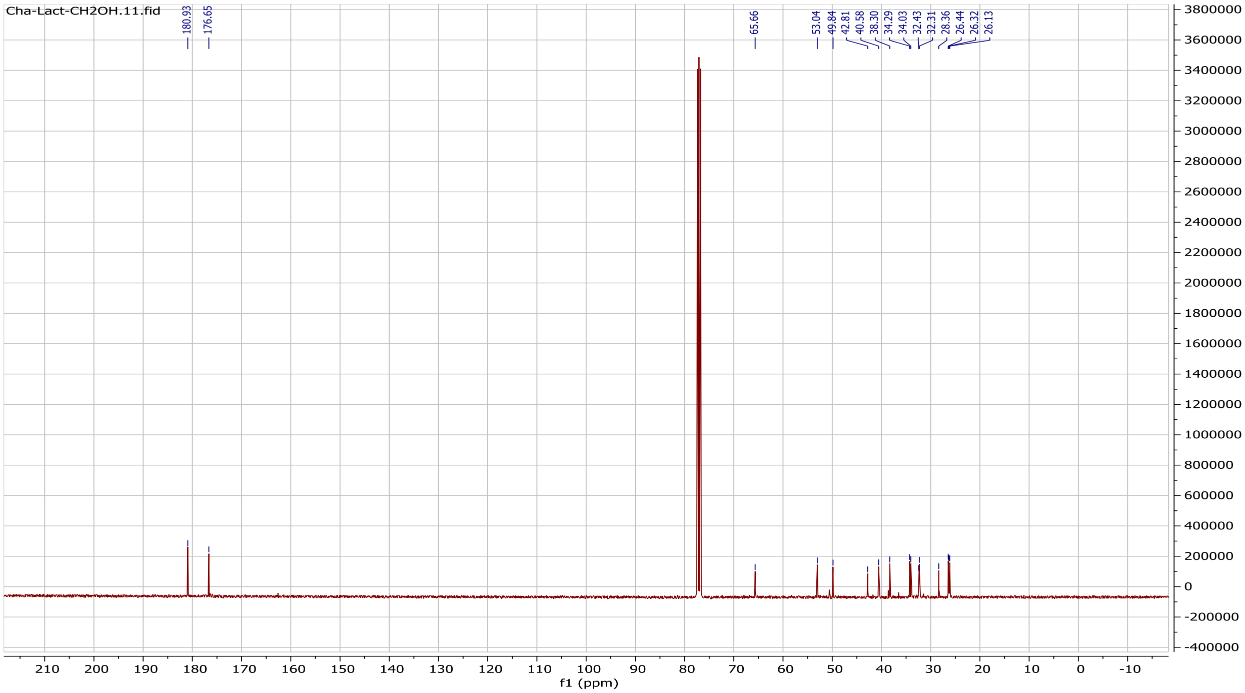
Cha-Lact-CH2OH.11.fid
180.93
176.65
65.66
53.04
49.84
42.81
40.58
38.30
34.29
34.03
32.43
32.31
28.36
26.44
26.32
26.13
f1 (ppm)

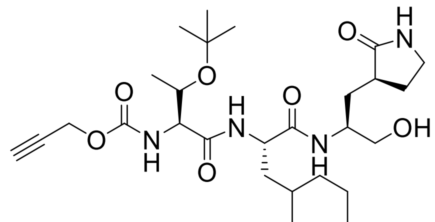
O
H
N
O
O
N
H
O
H
N
O
N
H
OH
O

MPD5c

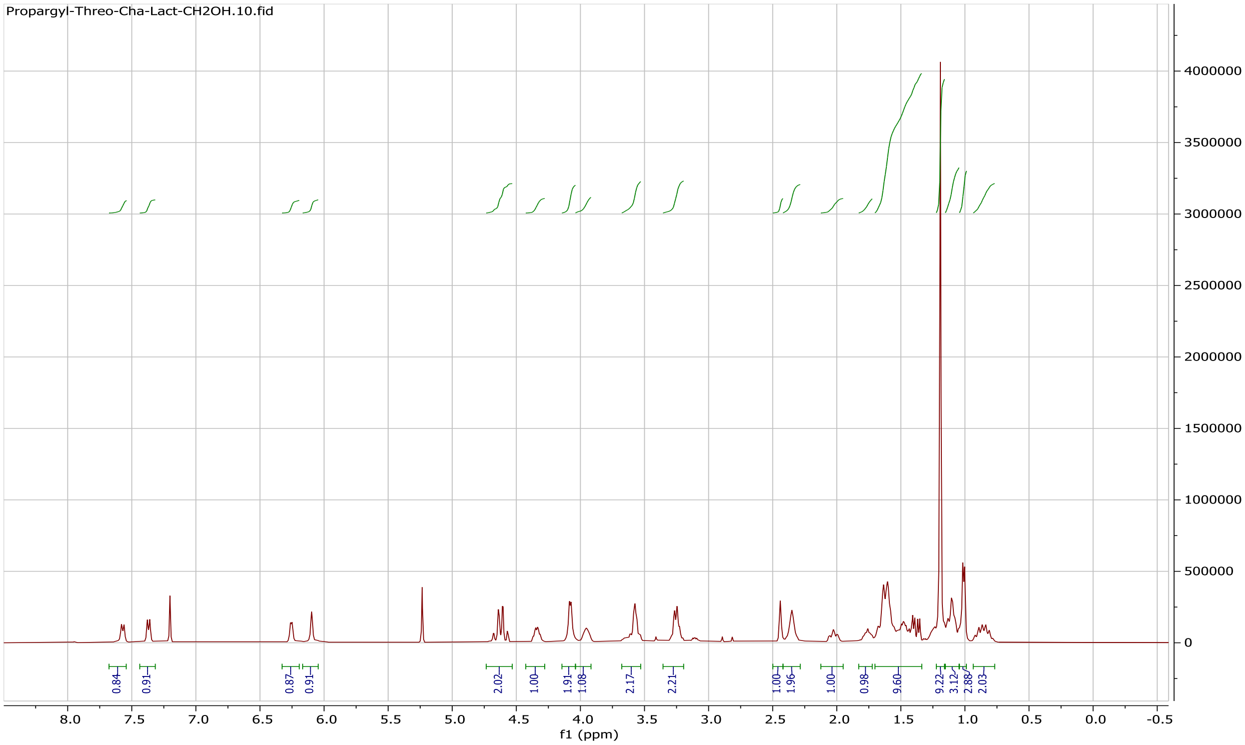
Propargyl-Threo-Cha-Lact-CH2OH.10.fid
f1 (ppm)

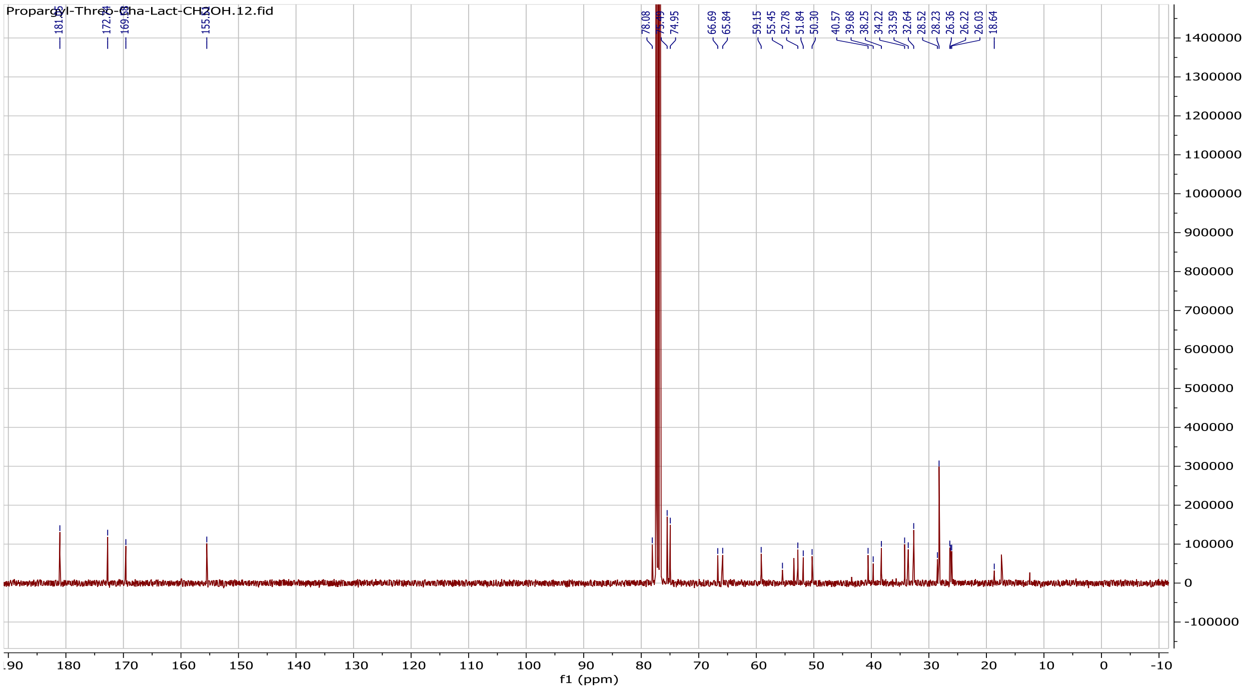
f1 (ppm)
78.08
74.95
66.69
65.84
59.15
55.45
52.78
51.84
50.30
40.57
39.68
38.25
34.22
33.59
32.64
28.52
28.23
26.36
26.22
26.03
18.64

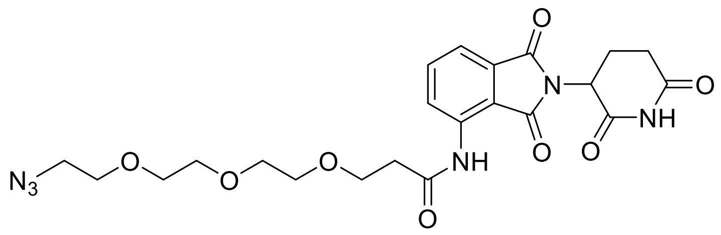

MPD5e

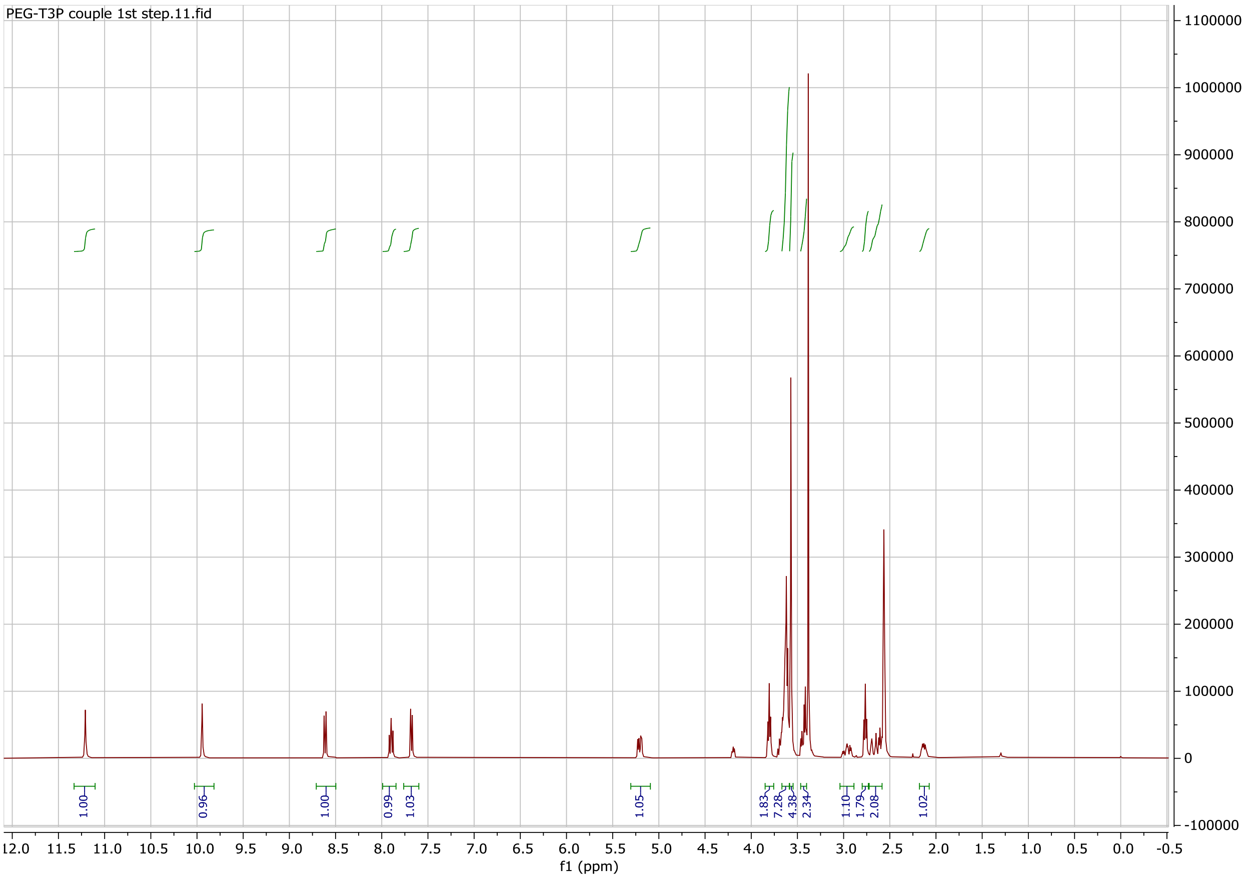

PEG-T3P couple 1st step.11.fid
1.00
0.96
1.00
0.99
1.03
1.05
1.83
7.28
4.38
2.34
1.10
1.79
2.08
1.02
f1 (ppm)

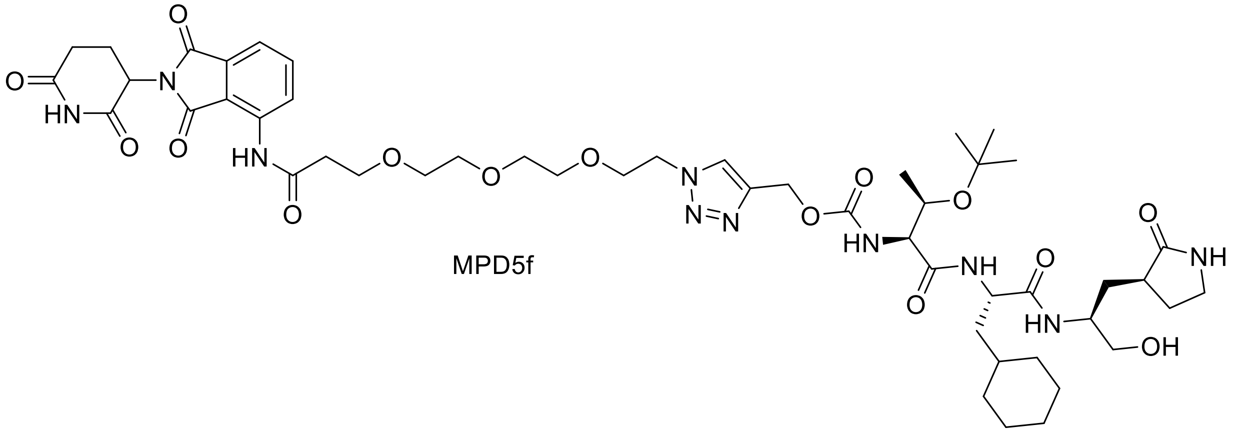

MPD5f

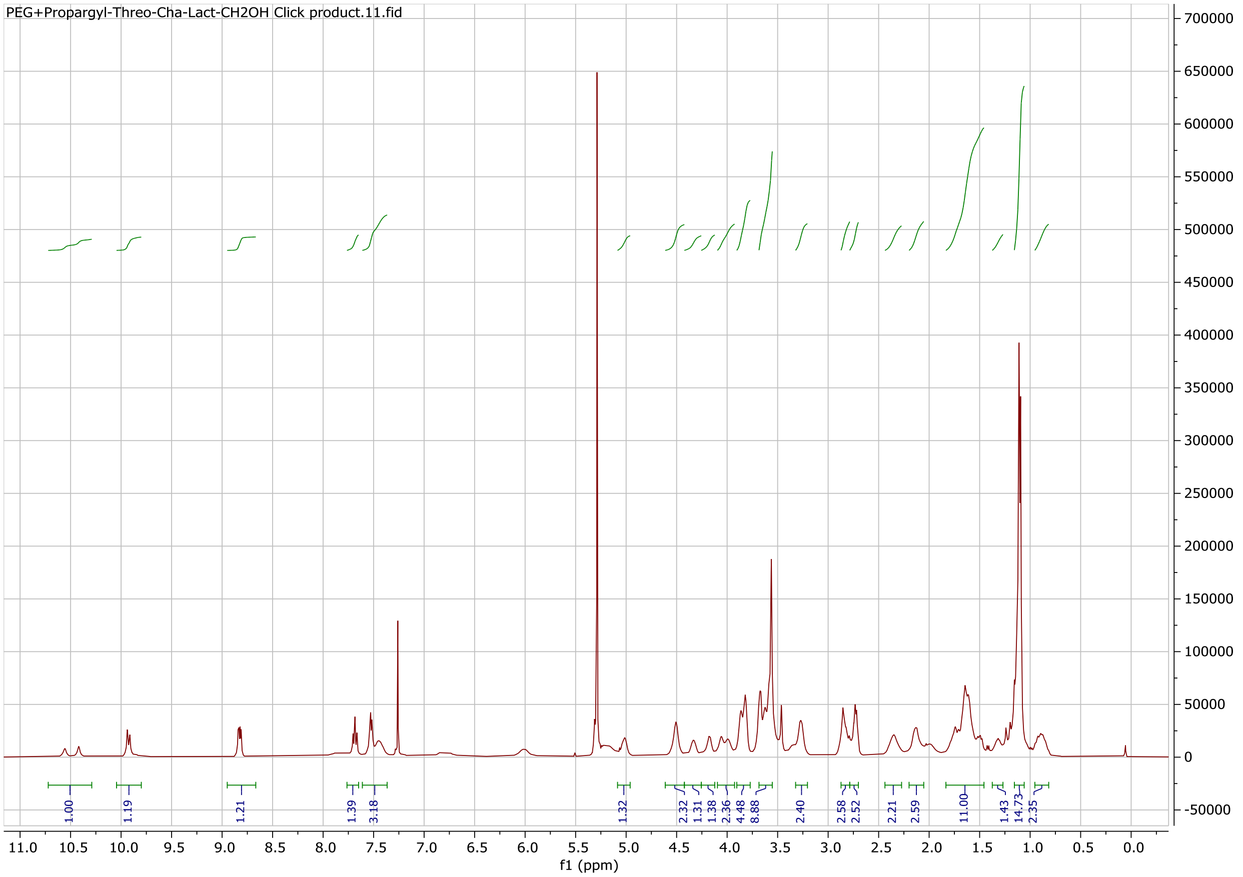

PEG+Propargyl-Threo-Cha-Lact-CH2OH Click product.11.fid
f1 (ppm)

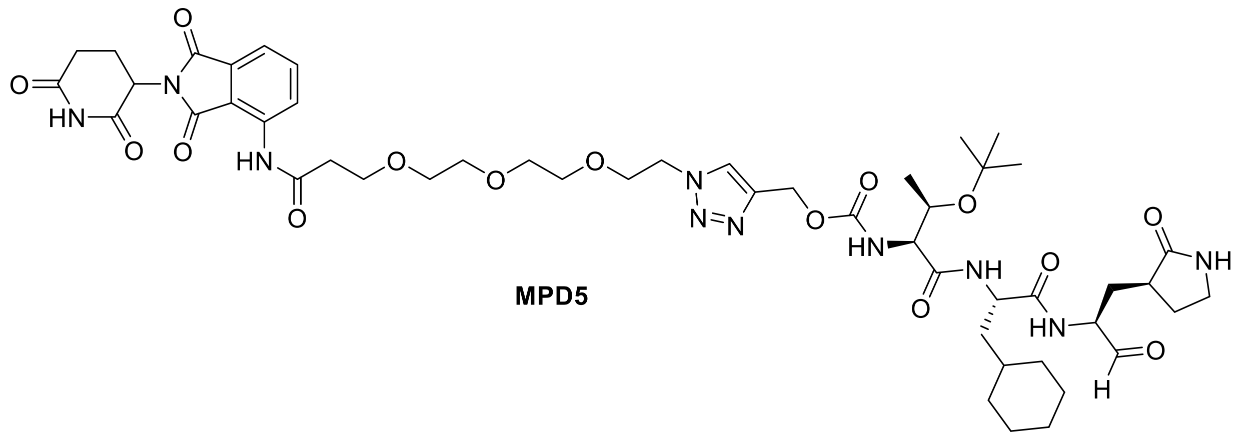

**MPD5**

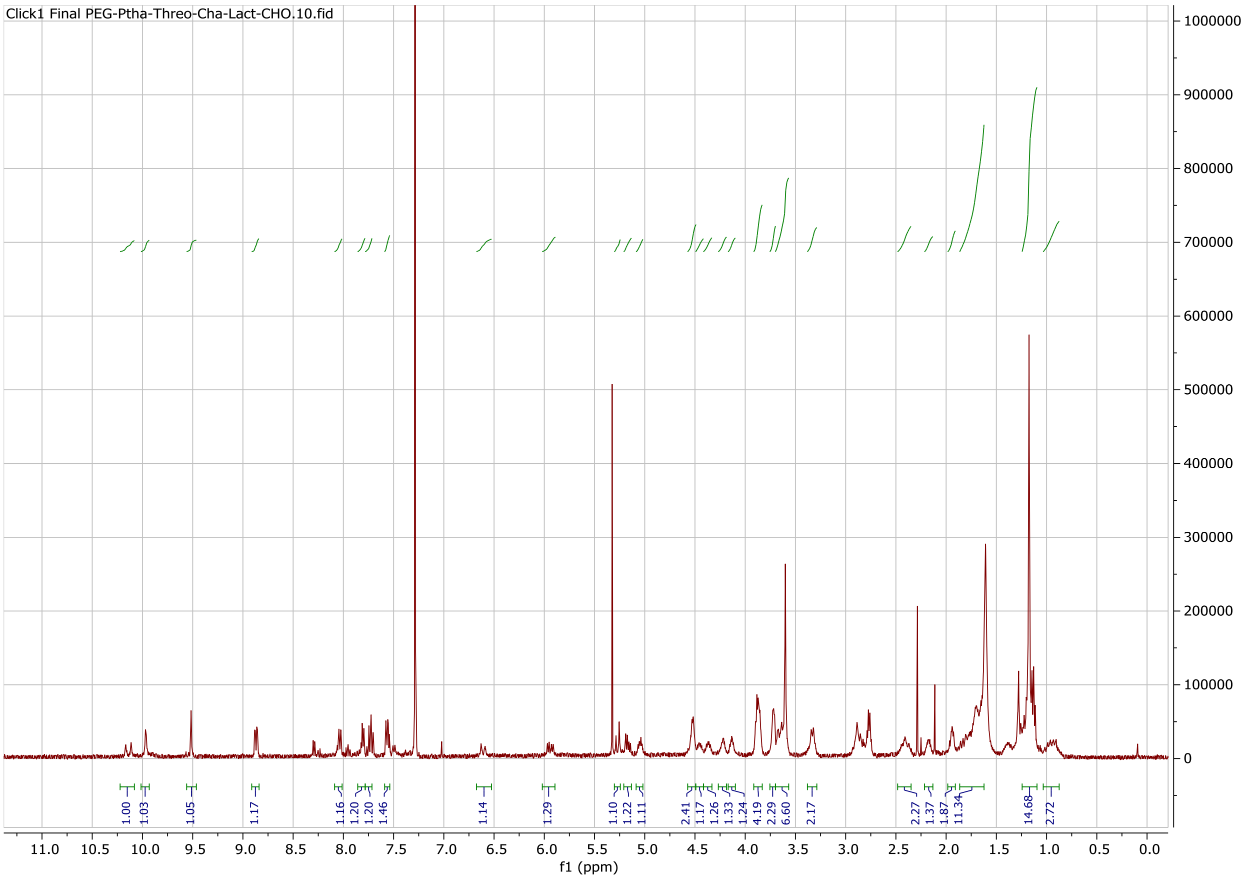
Click1 Final PEG-Ptha-Threo-Cha-Lact-CHO.10.fid
1000000
900000
800000
700000
600000
500000
400000
300000
200000
100000
0
1.00
1.03
1.05
1.17
1.16
1.20
1.20
1.46
1.14
1.29
1.10
1.22
1.11
2.41
1.17
1.26
1.33
1.24
4.19
2.29
6.60
2.17
2.27
1.37
1.87
11.34
14.68
2.72
11.0 10.5 10.0 9.5 9.0 8.5 8.0 7.5 7.0 6.5 6.0 5.5 5.0 4.5 4.0 3.5 3.0 2.5 2.0 1.5 1.0 0.5 0.0
f1 (ppm)

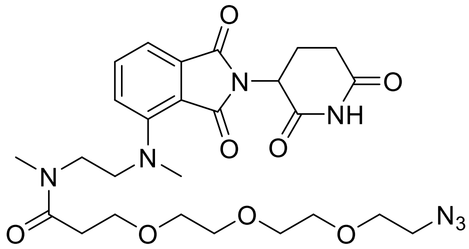

MPD6a

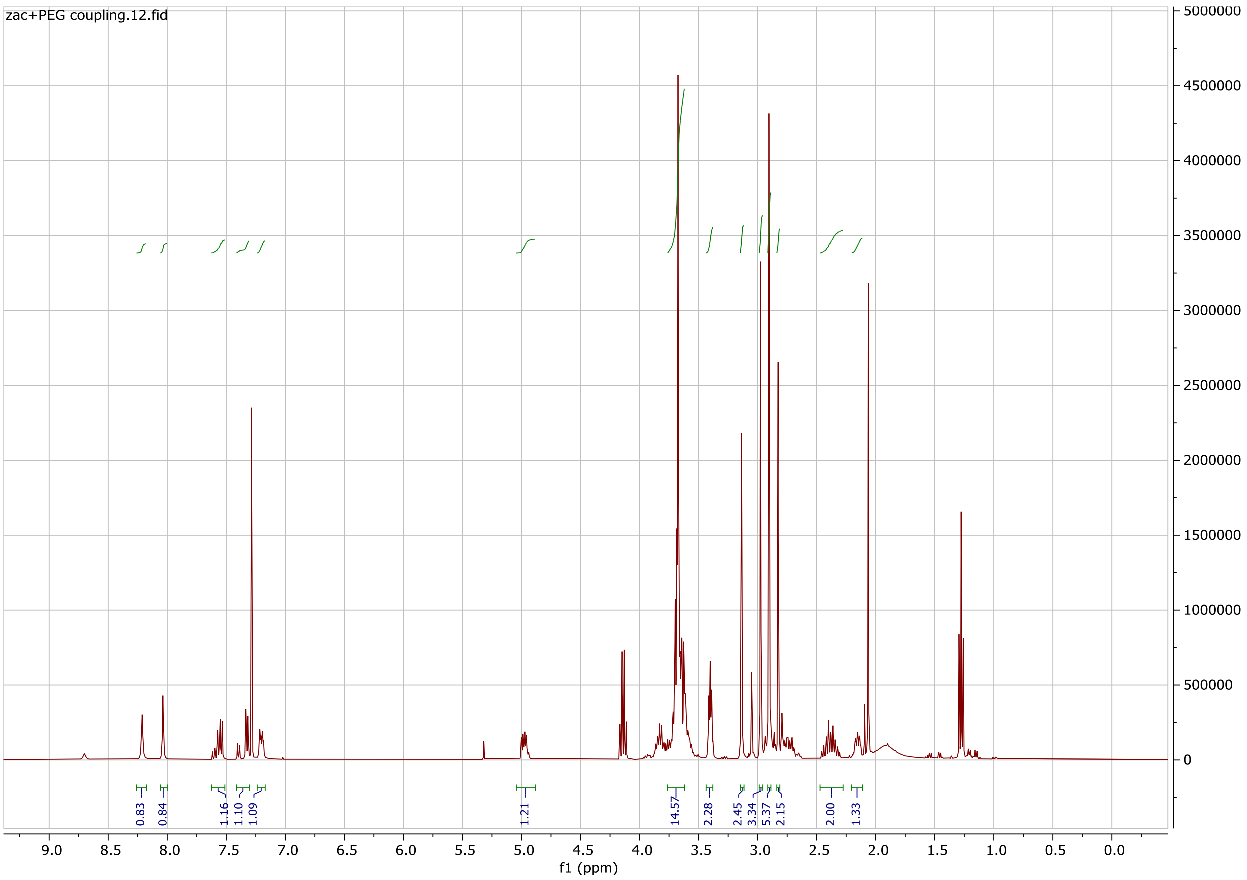
zac+PEG coupling.12.fid
f1 (ppm)
9.0 8.5 8.0 7.5 7.0 6.5 6.0 5.5 5.0 4.5 4.0 3.5 3.0 2.5 2.0 1.5 1.0 0.5 0.0
5000000 4500000 4000000 3500000 3000000 2500000 2000000 1500000 1000000 500000 0
0.83 0.84 1.16 1.10 1.09 1.21 14.57 2.28 2.45 3.34 5.37 2.15 2.00 1.33

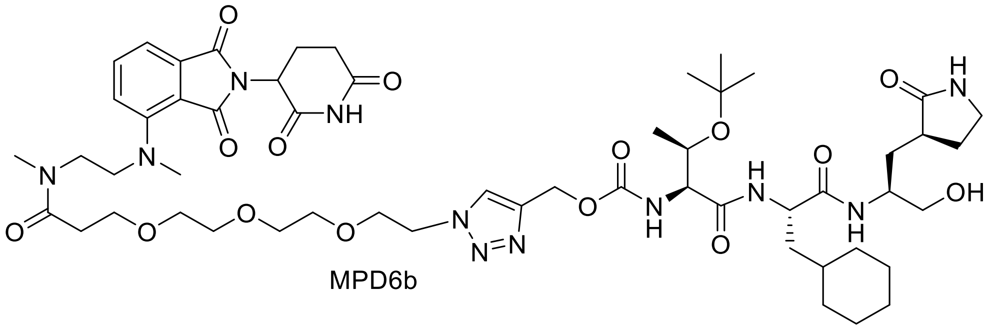

MPD6b

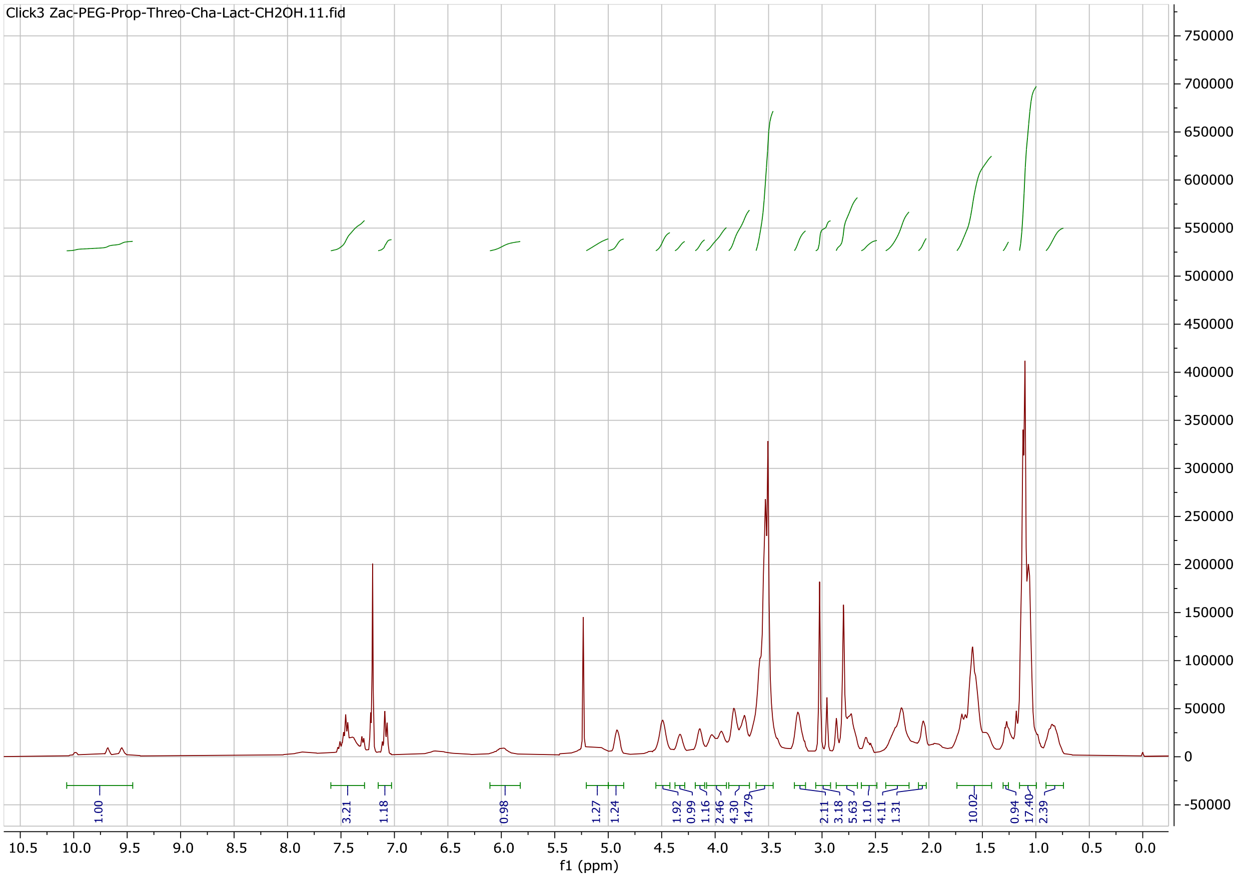

Click3 Zac-PEG-Prop-Threo-Cha-Lact-CH2OH.11.fid
1.00
3.21
1.18
0.98
1.27
1.24
1.92
0.99
1.16
2.46
4.30
14.79
2.11
3.18
5.63
1.10
4.11
1.31
10.02
0.94
17.40
2.39
f1 (ppm)

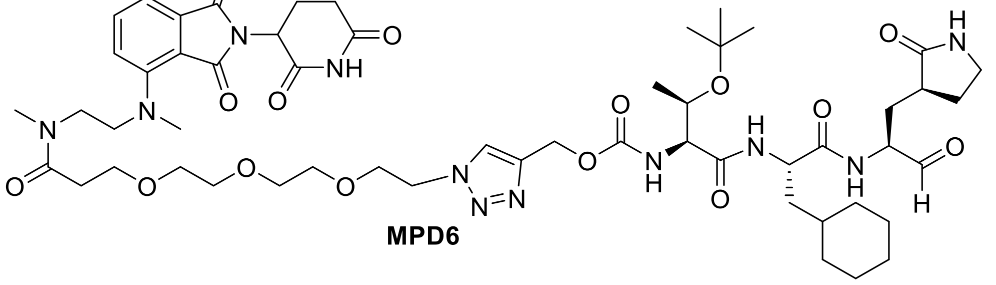

**MPD6**

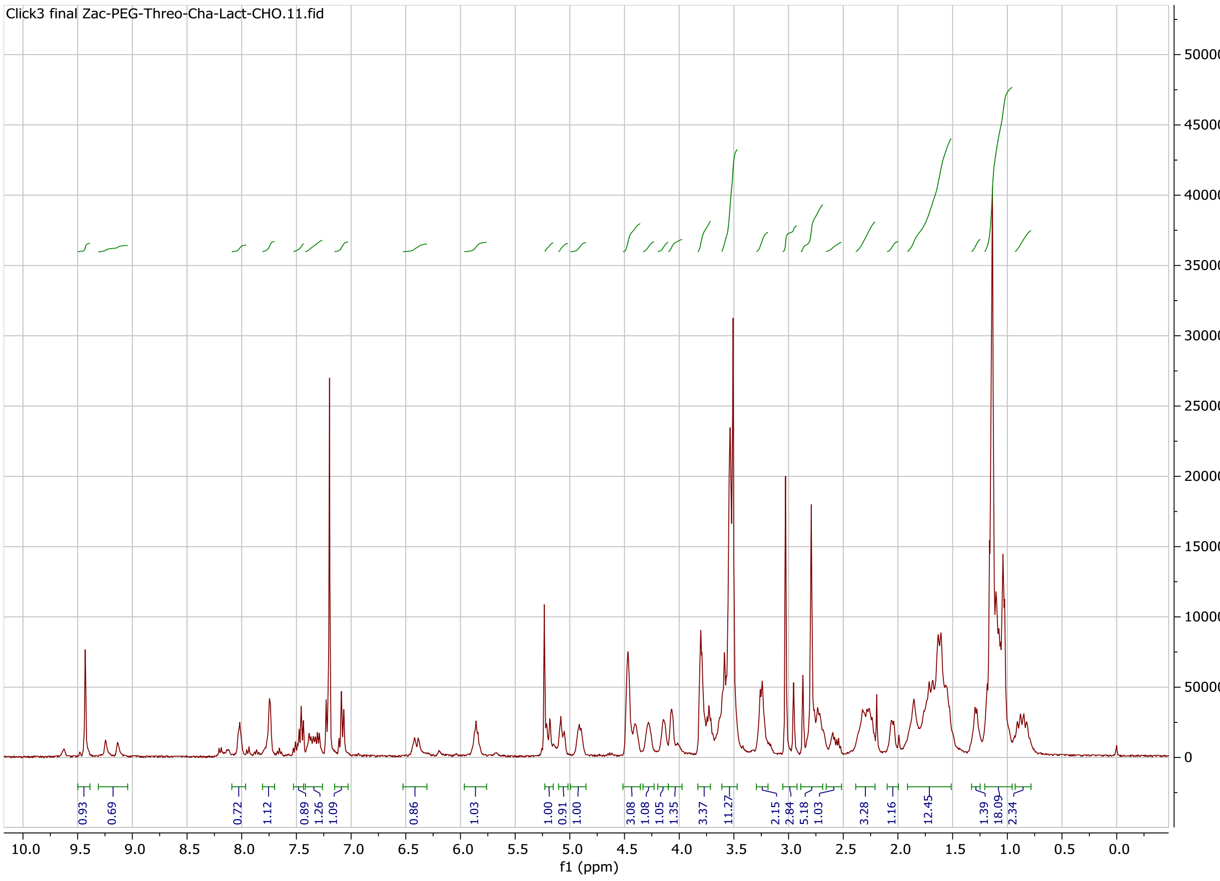

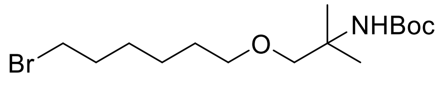
Br
O
NHBoc

MPD7b

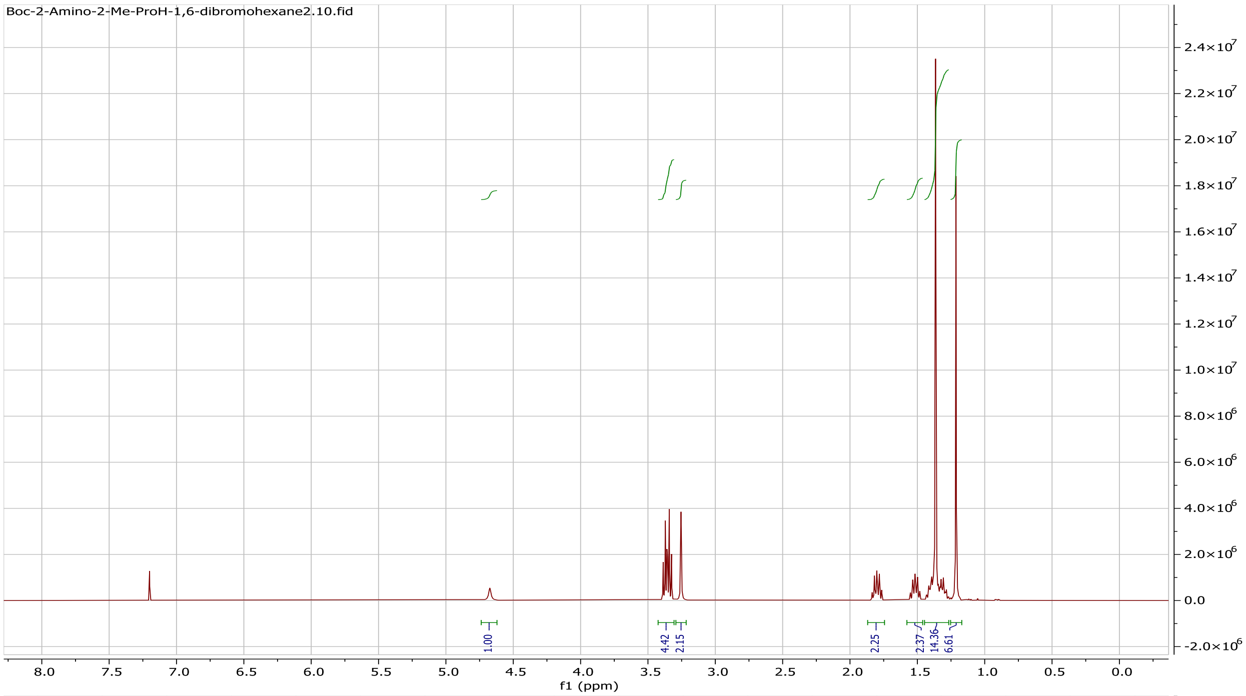
Boc-2-Amino-2-Me-ProH-1,6-dibromohexane2.10.fid
1.00
4.42
2.15
2.25
2.37
14.36
6.61
f1 (ppm)

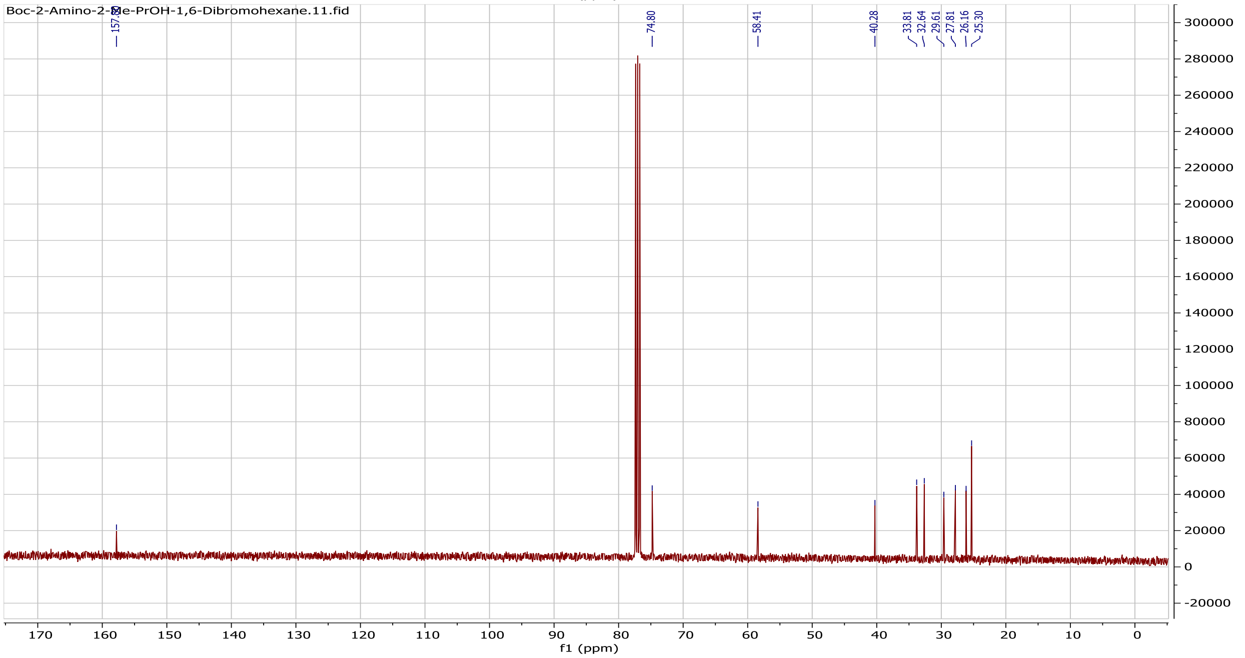
Boc-2-Amino-2-Me-PrOH-1,6-Dibromohexane.11.fid
74.80
58.41
40.28
33.81
32.64
29.61
27.81
26.16
25.30
f1 (ppm)

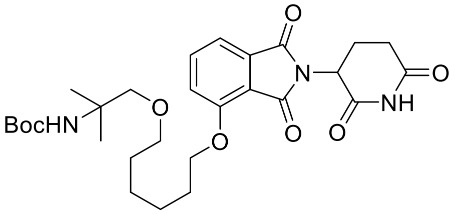

BocHN
O
O
O
N
O
O
NH
O


MPD7c

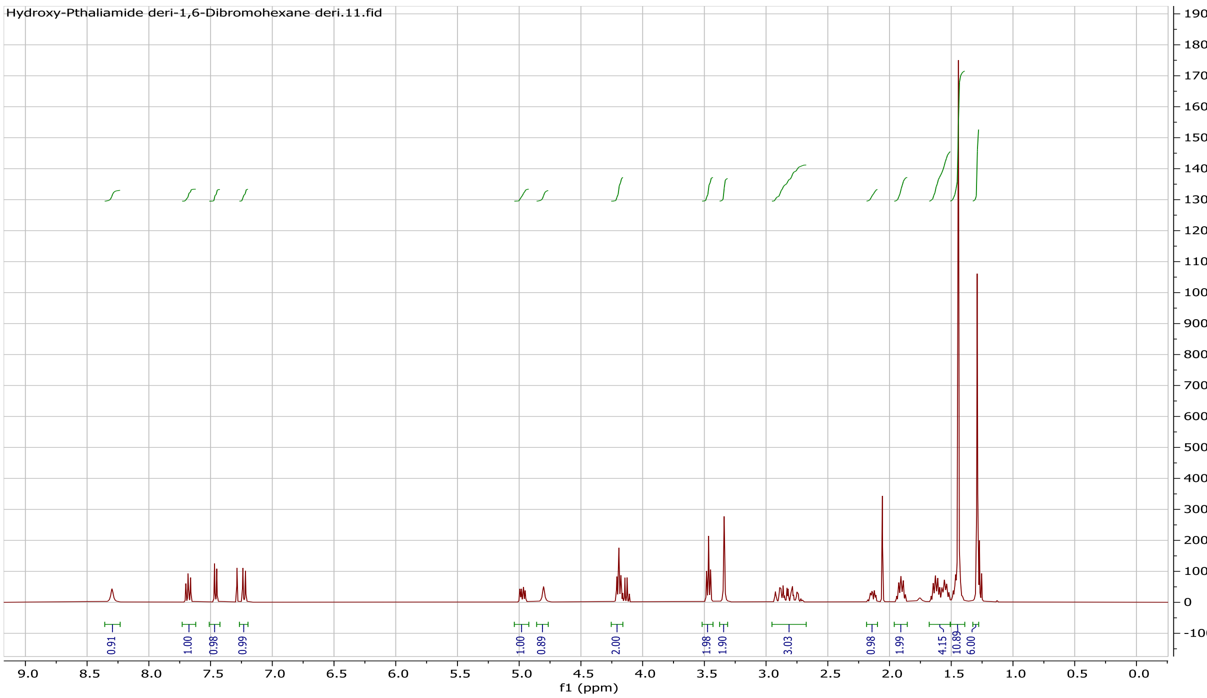

Hydroxy-Pthaliamide deri-1,6-Dibromohexane deri.11.fid
f1 (ppm)

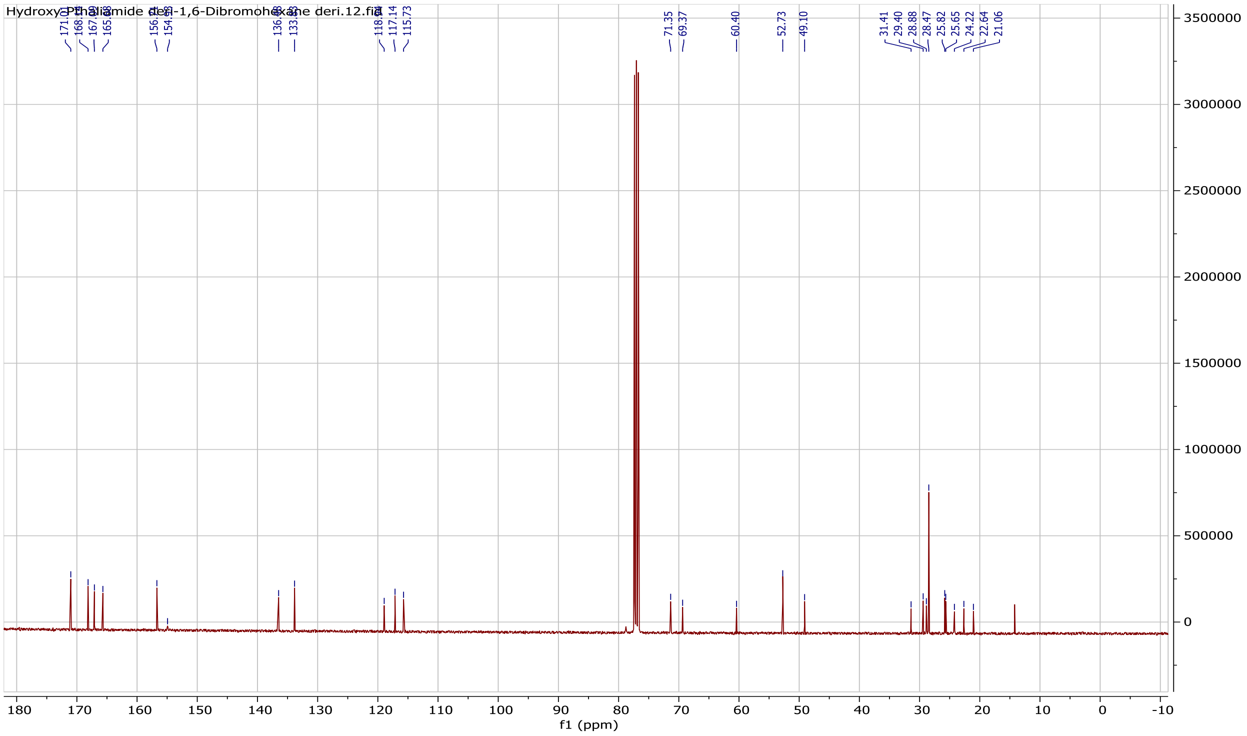
Hydroxy-Phthaliamide deri-1,6-Dibromohexane deri.12.fid
117.14
115.73
71.35
69.37
60.40
52.73
49.10
31.41
29.40
28.88
28.47
25.82
25.65
24.22
22.64
21.06
3500000
3000000
2500000
2000000
1500000
1000000
500000
0
180
170
160
150
140
130
120
110
100
90
80
70
60
50
40
30
20
10
0
-10
f1 (ppm)

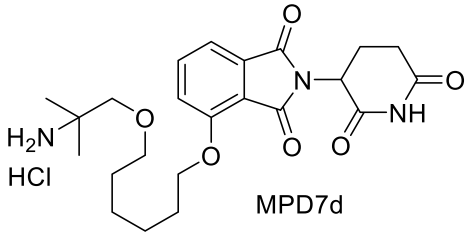
H2N
HCl
O
N
NH

MPD7d

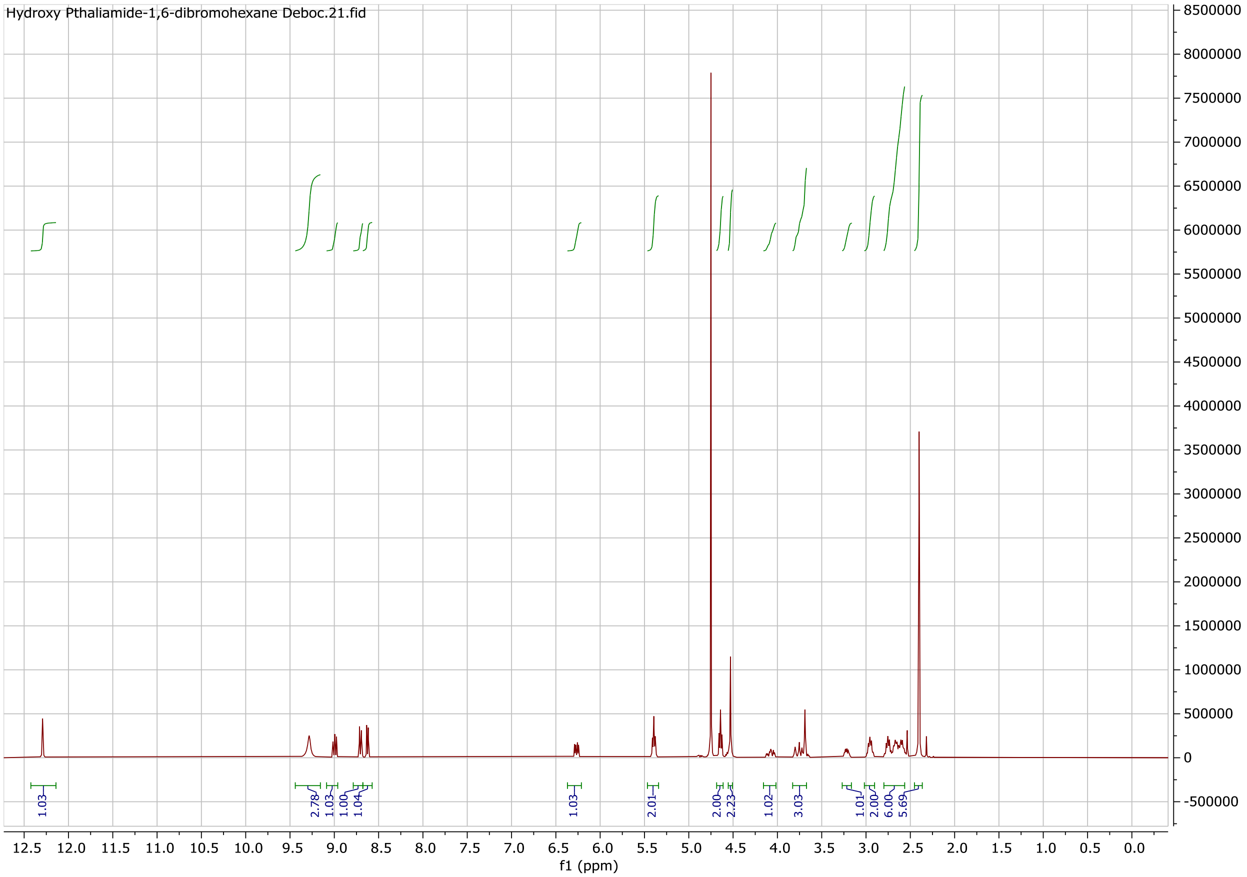
Hydroxy Pthaliamide-1,6-dibromohexane Deboc.21.fid
f1 (ppm)
1.03
2.78
1.03
1.00
1.04
1.03
2.01
2.00
2.23
1.02
3.03
1.01
2.00
6.00
5.69

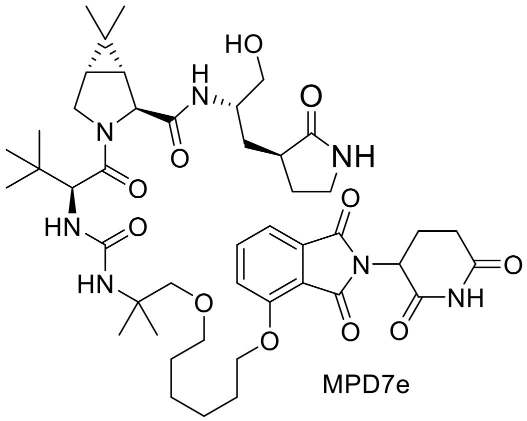

MPD7e

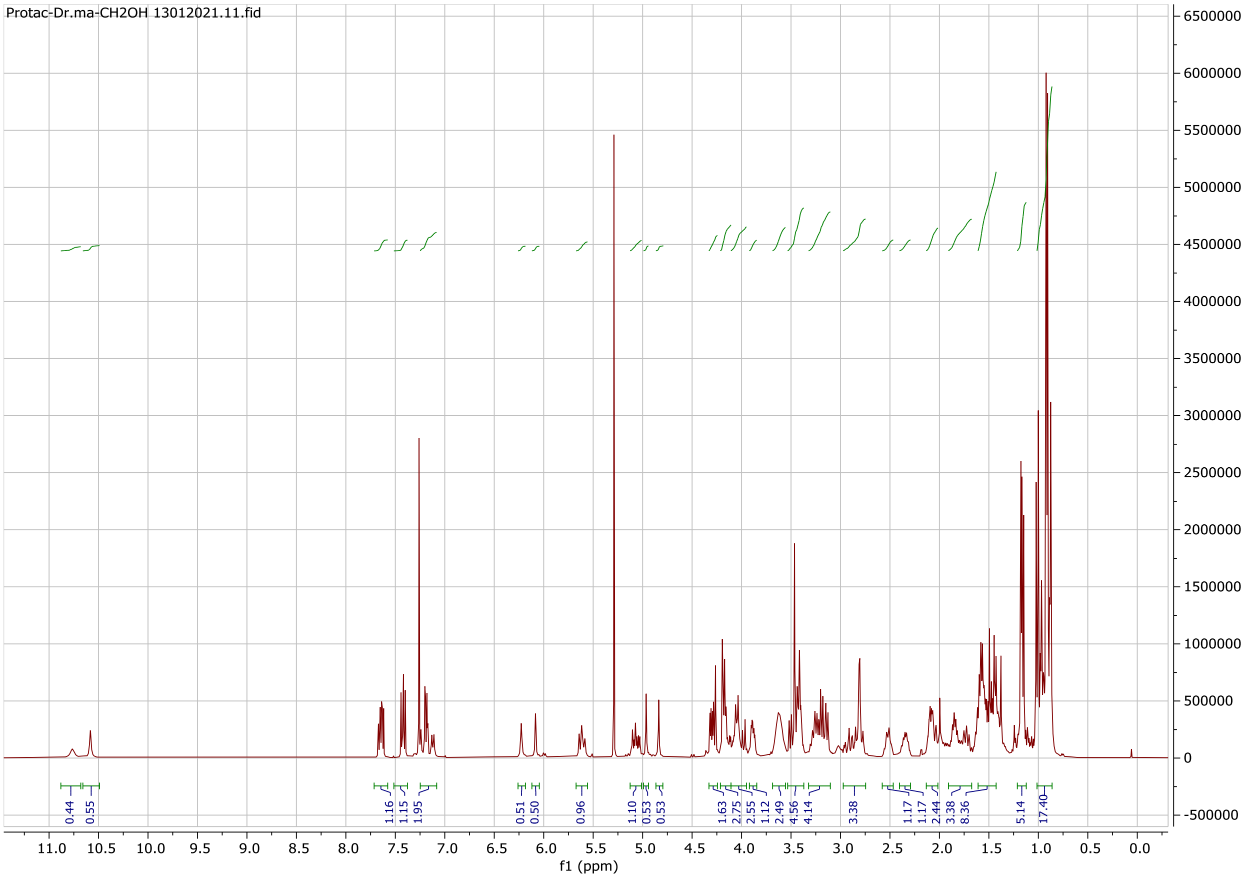

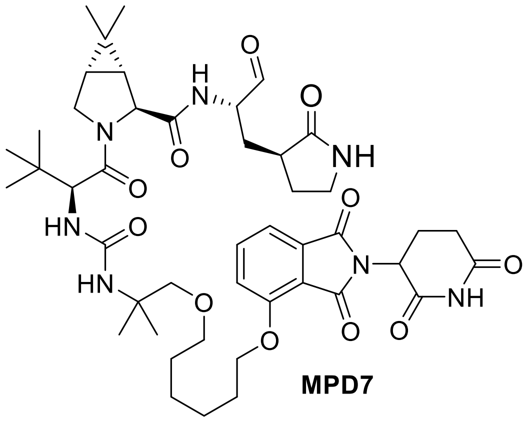

**MPD7**

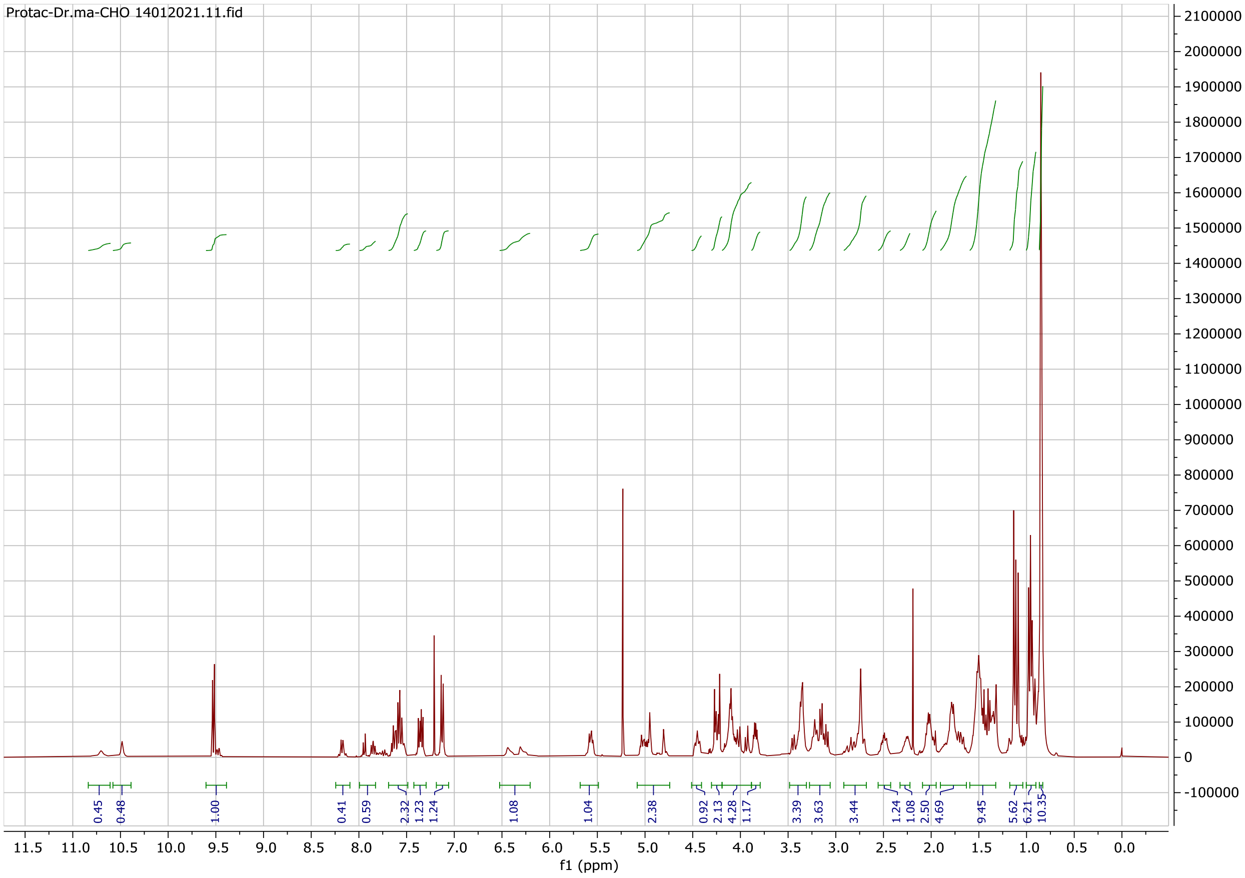

Protac-Dr.ma-CHO 14012021.11.fid
0.45
0.48
1.00
0.41
0.59
2.32
1.23
1.24
1.08
1.04
2.38
0.92
2.13
4.28
1.17
3.39
3.63
3.44
1.24
1.08
2.50
4.69
9.45
5.62
6.21
10.35
f1 (ppm)

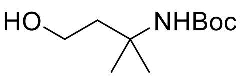

MPD8a

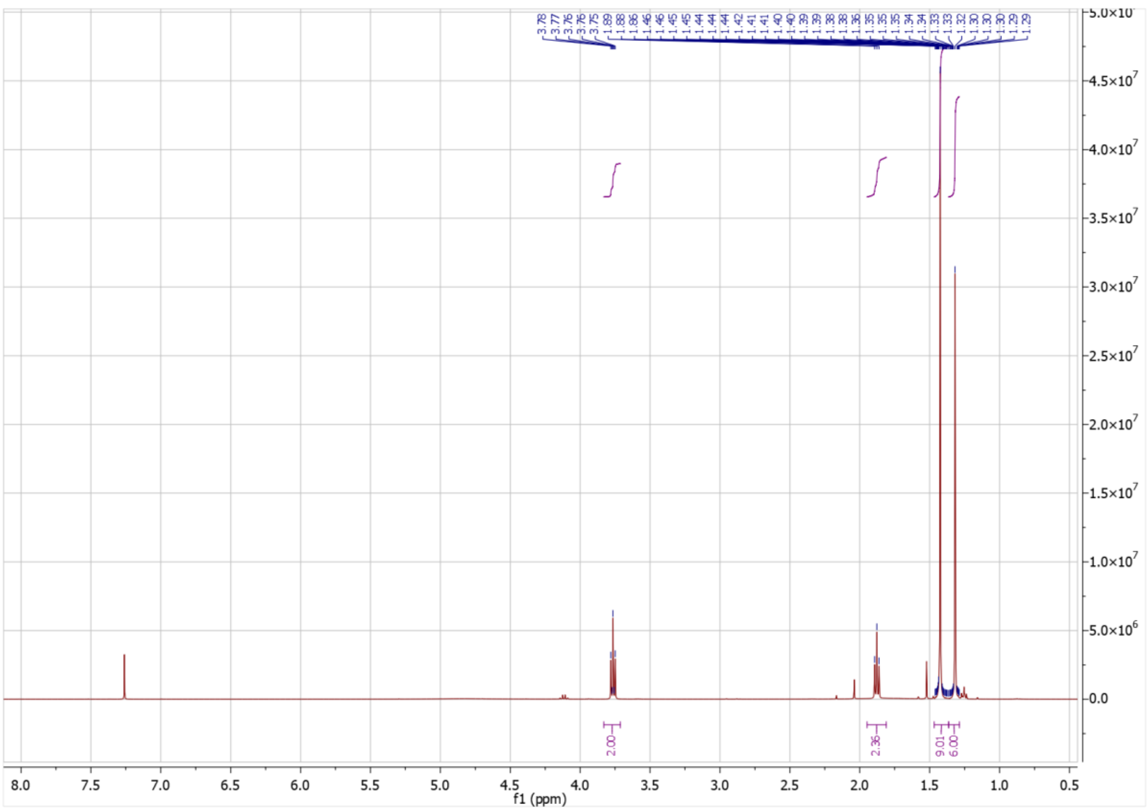

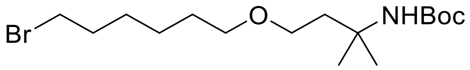

Br
O
NHBoc


MPD8b

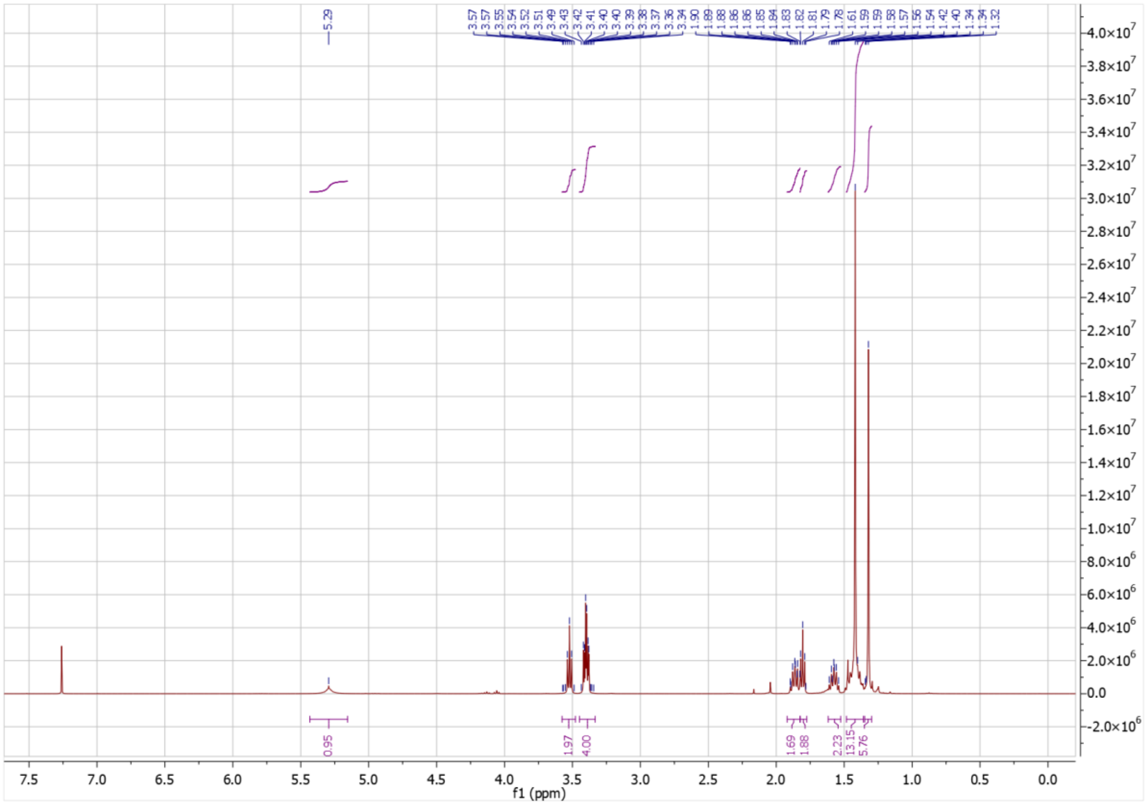

5.29
3.57 3.57 3.55 3.54 3.52 3.51 3.49 3.43 3.42 3.41 3.40 3.39 3.38 3.37 3.36 3.34
1.90 1.89 1.88 1.86 1.86 1.85 1.84 1.83 1.82 1.81 1.79 1.78 1.61 1.59 1.59 1.59 1.58 1.57 1.56 1.54 1.42 1.40 1.34 1.32
0.95
1.97
4.00
1.69
1.88
2.23
13.15
5.76
f1 (ppm)

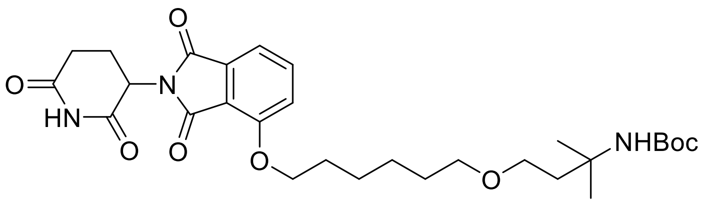


MPD8c

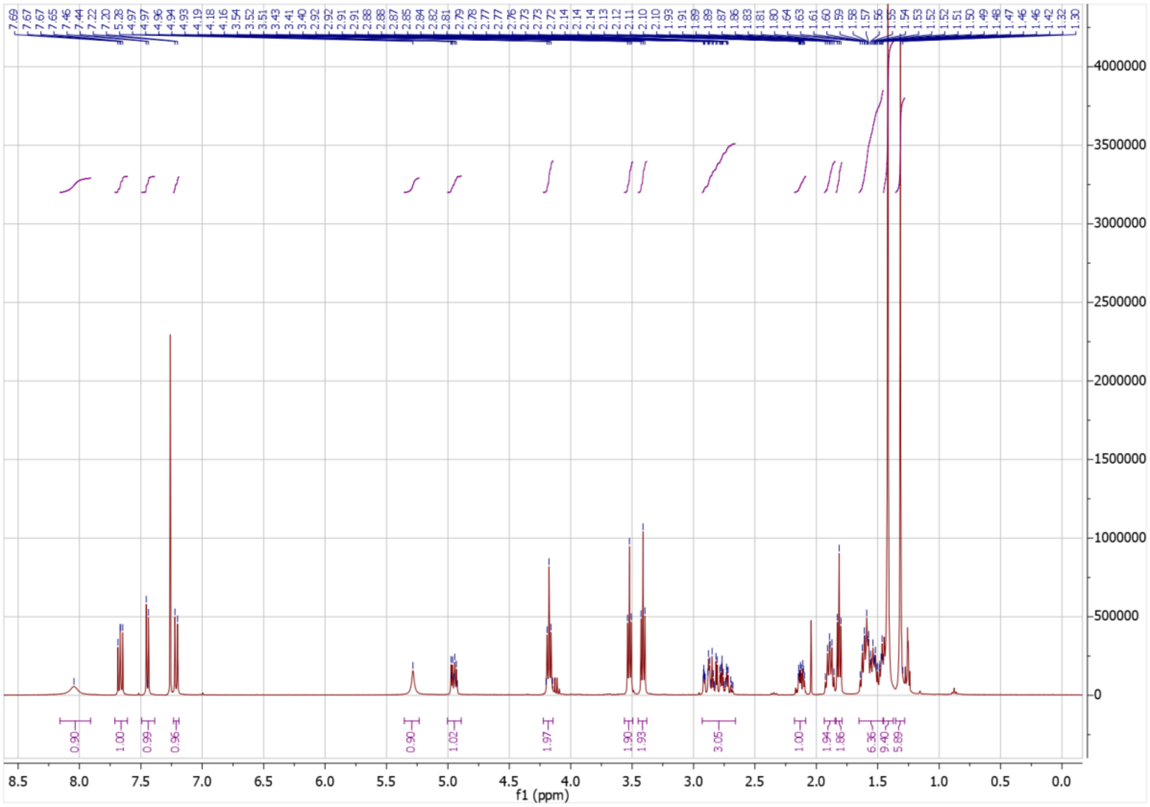

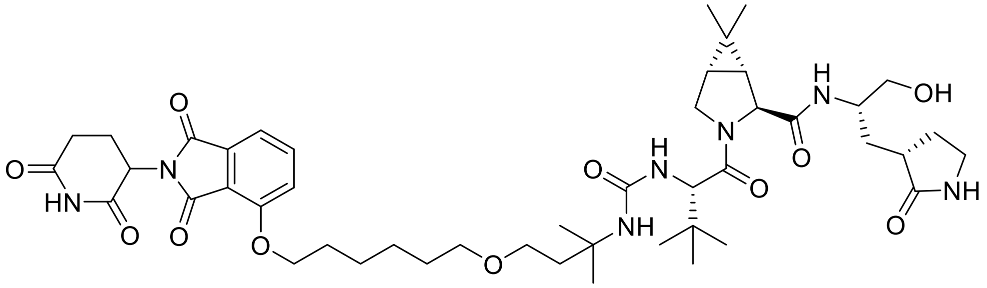

MPD8e

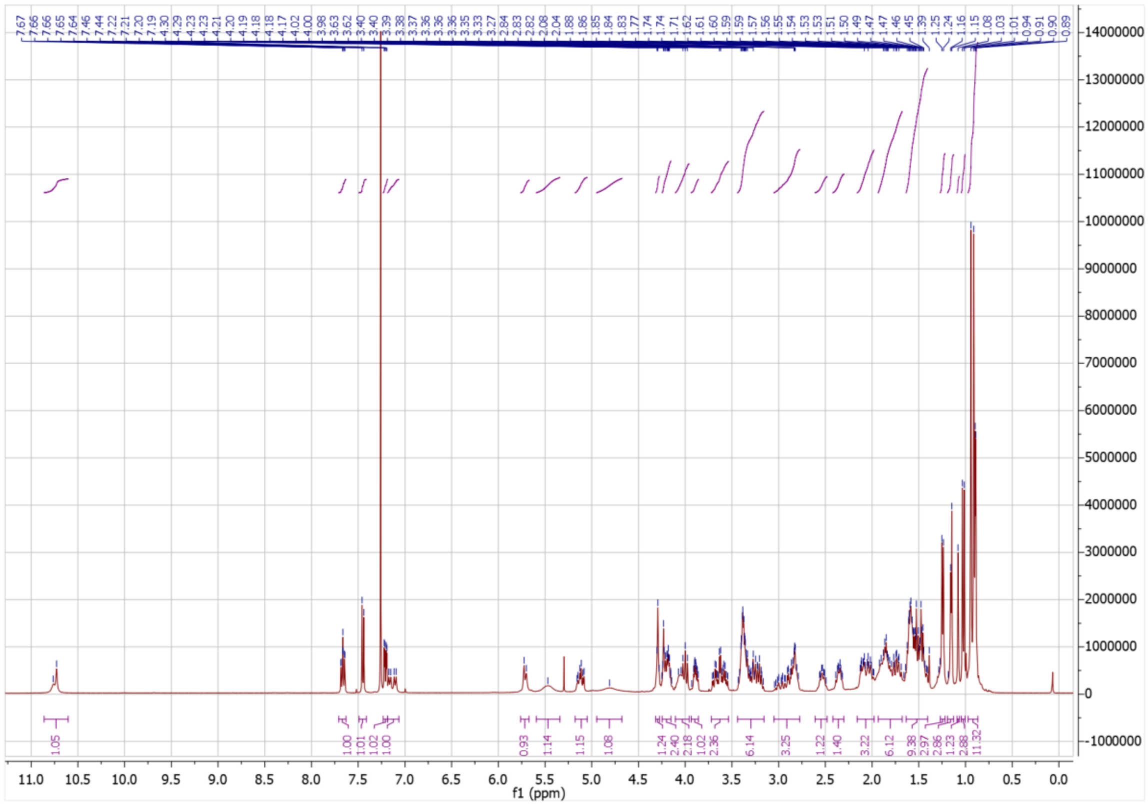

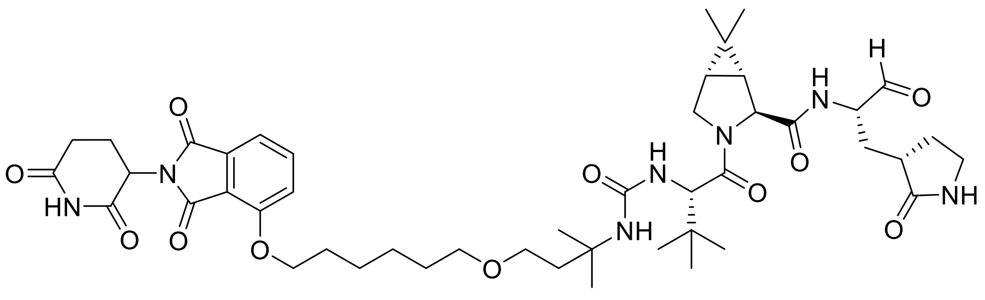

MPD8

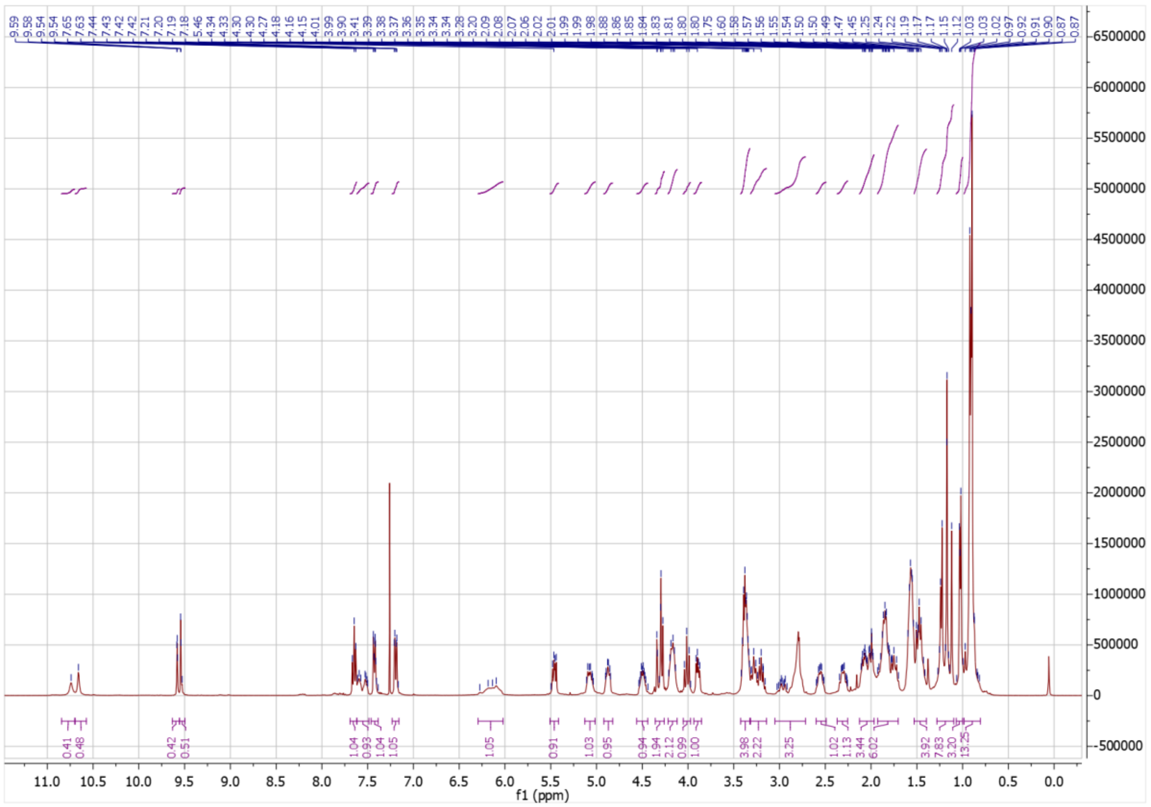

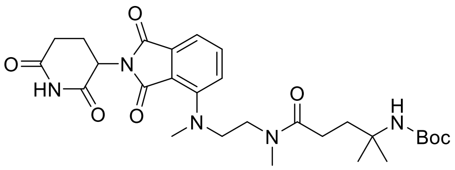

**MPD9b**

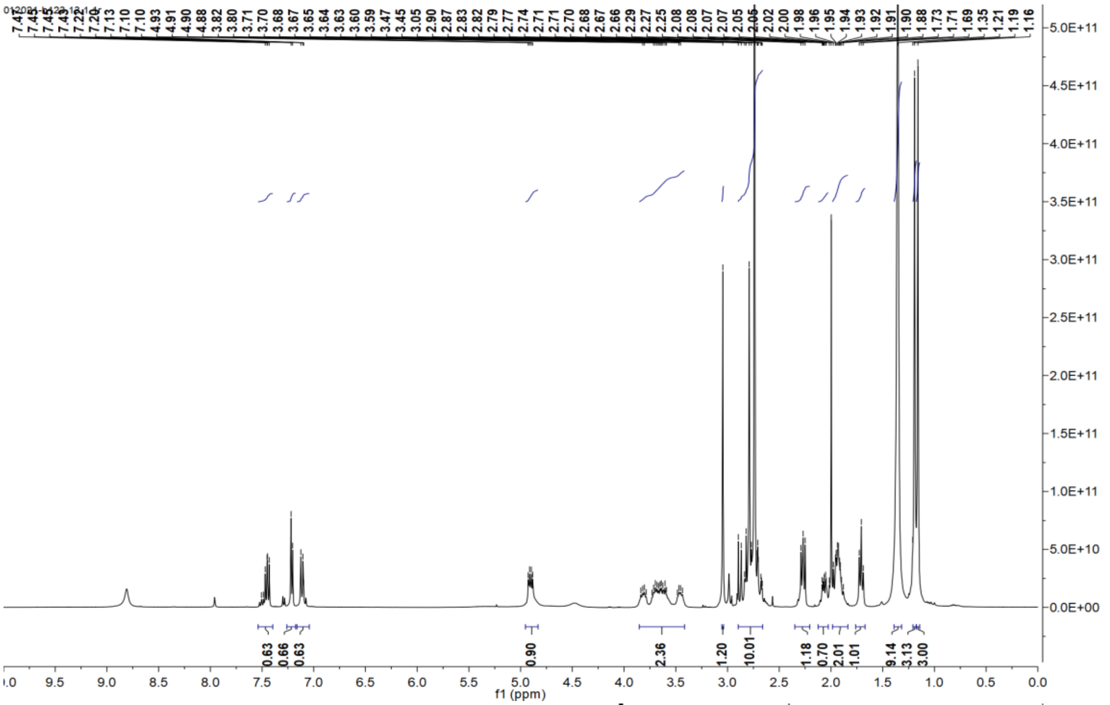

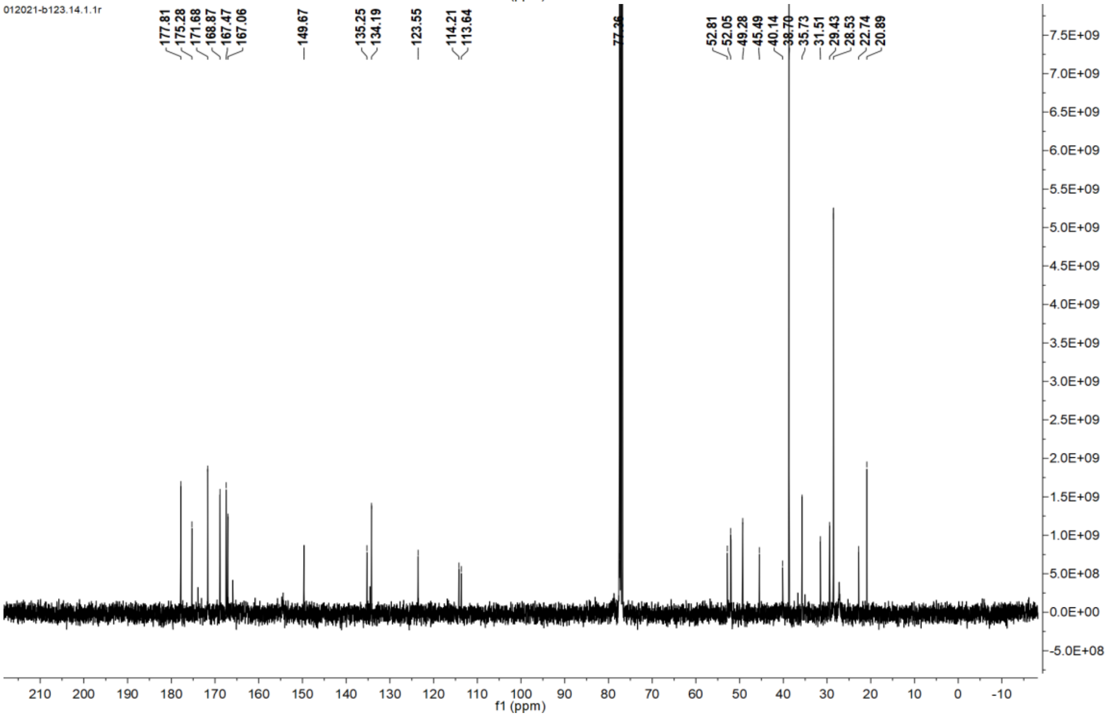

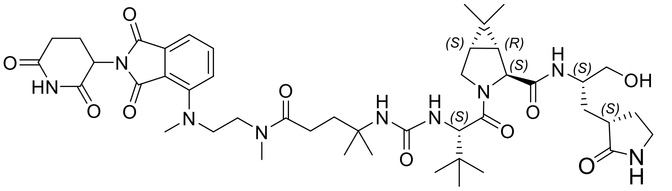
O
HN
O
N
O
O
N
N
O
H
N
H
N
O
(S)
O
(S)
(R)
(S)
N
O
H
N
(S)
OH
(S)
O
NH

**MPD9d**

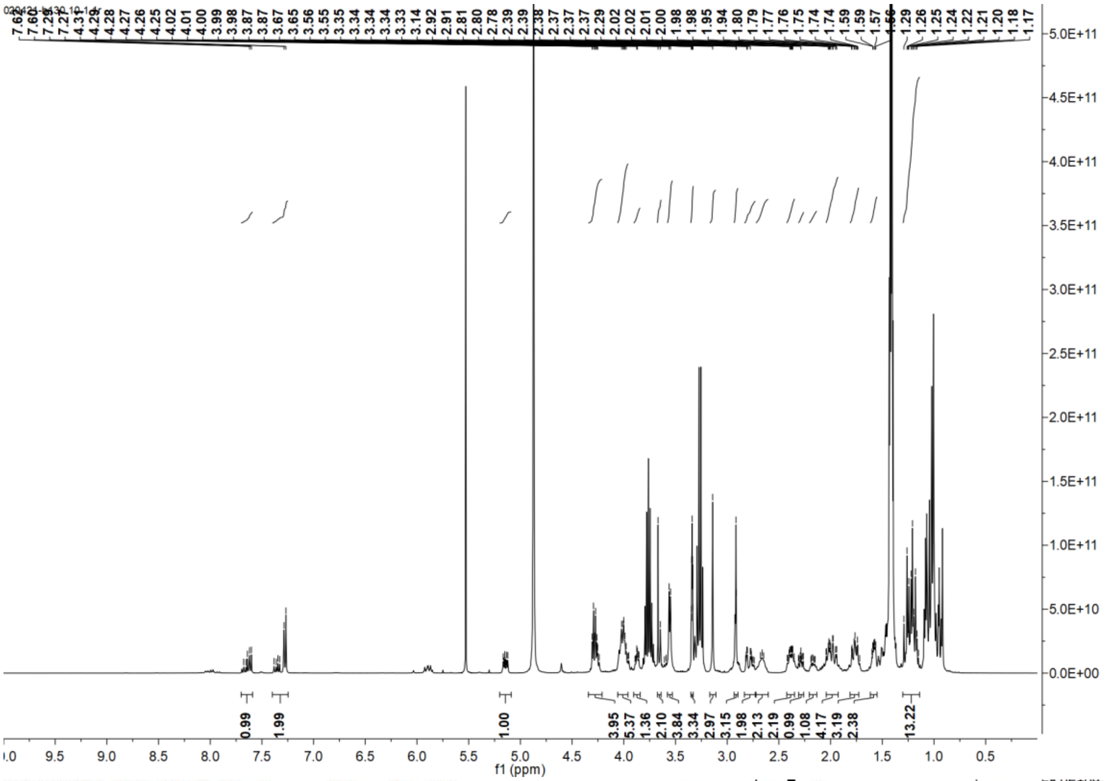
7.62 7.60 7.29 7.27 4.31 4.29 4.28 4.27 4.26 4.25 4.02 4.01 4.00 3.99 3.98 3.87 3.87 3.67 3.65 3.56 3.55 3.35 3.34 3.34 3.34 3.33 3.14 2.92 2.91 2.81 2.80 2.78 2.39 2.39 2.38 2.37 2.37 2.37 2.29 2.02 2.02 2.01 2.00 1.98 1.98 1.98 1.95 1.94 1.80 1.79 1.77 1.76 1.75 1.74 1.74 1.74 1.59 1.59 1.57 1.29 1.26 1.25 1.24 1.22 1.21 1.20 1.18 1.17
0.99 1.99 1.00 3.95 5.37 1.36 2.10 3.84 3.34 2.97 3.15 1.98 2.13 2.19 0.99 1.08 4.17 3.19 2.38 13.22
f1 (ppm)

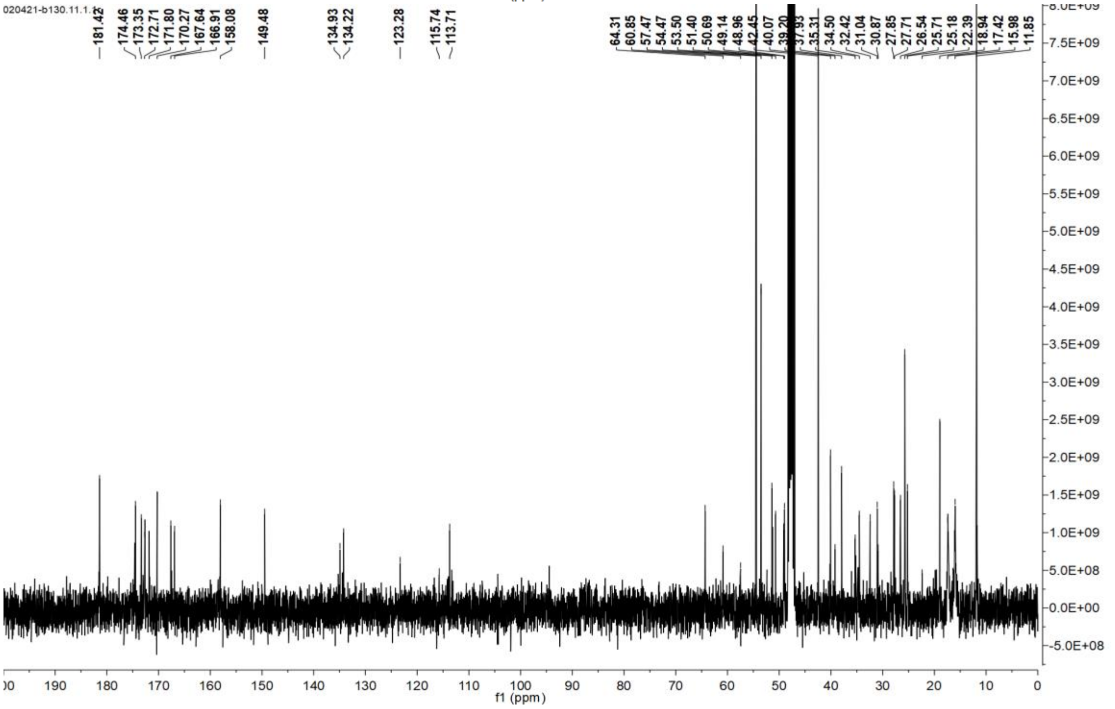
181.42 174.46 173.35 172.71 171.80 170.27 167.64 166.91 158.08 149.48 134.93 134.22 123.28 115.74 113.71
64.31 60.85 57.47 54.47 53.50 51.40 50.69 49.14 48.96 42.45 40.07 39.20 37.93 35.31 34.50 32.42 31.04 30.87 27.85 27.71 26.54 25.71 25.18 22.39 18.94 17.42 15.98 11.85
f1 (ppm)

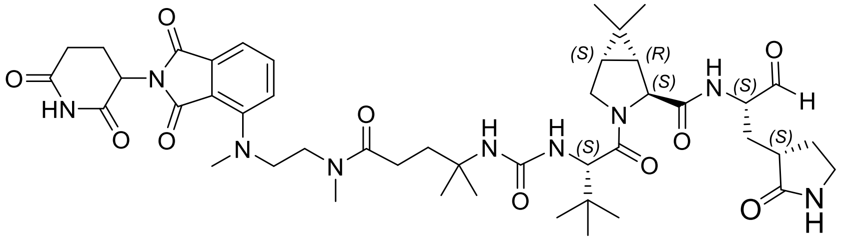

MPD9

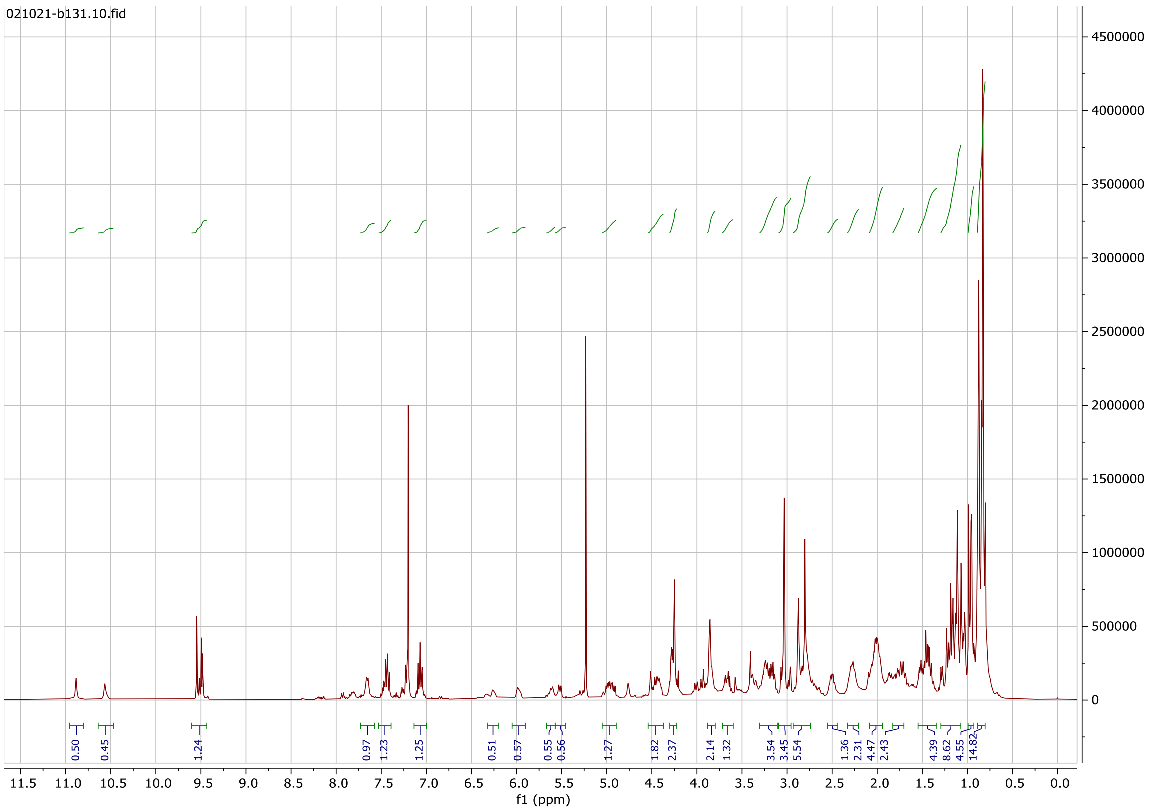

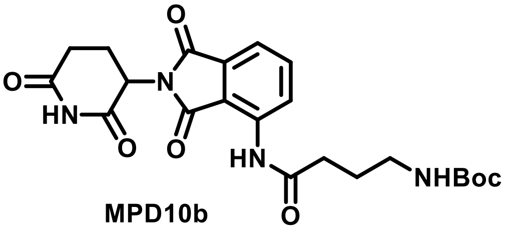

**MPD10b**

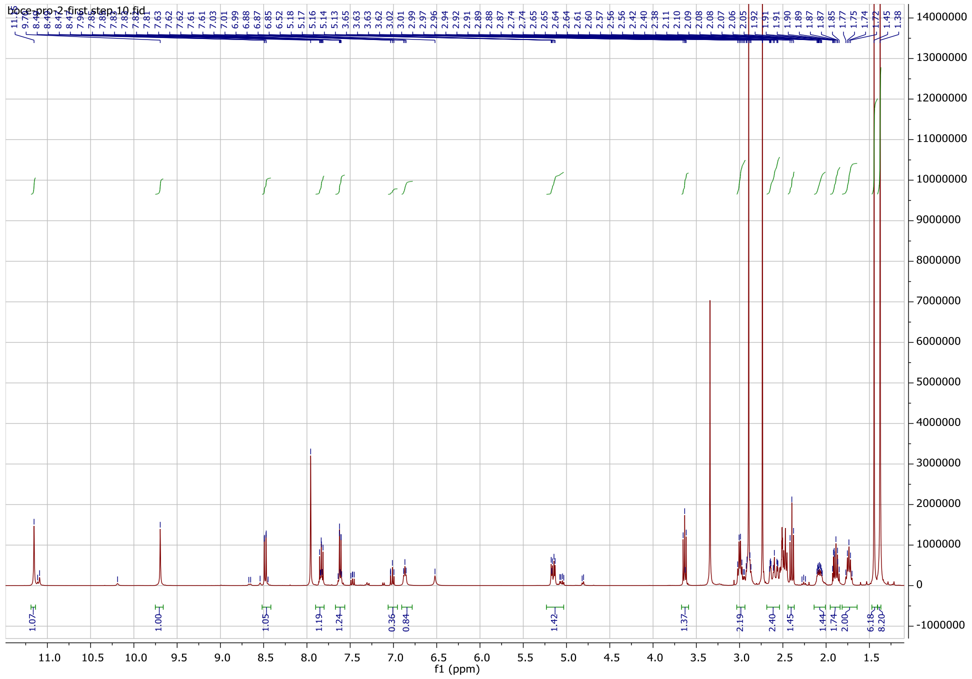

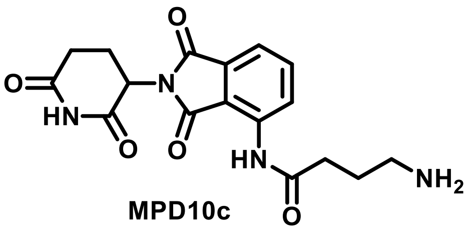
O
O
N
HN
O
O
HN
NH2
O

MPD10c

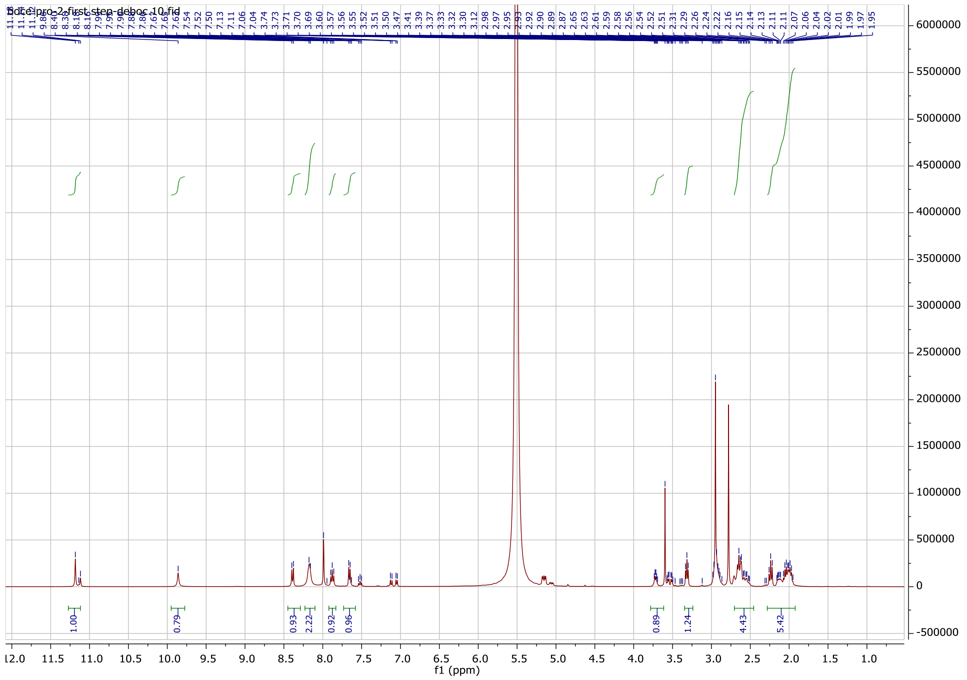
boce-pro-2-first-step-deboc.10.fid
f1 (ppm)

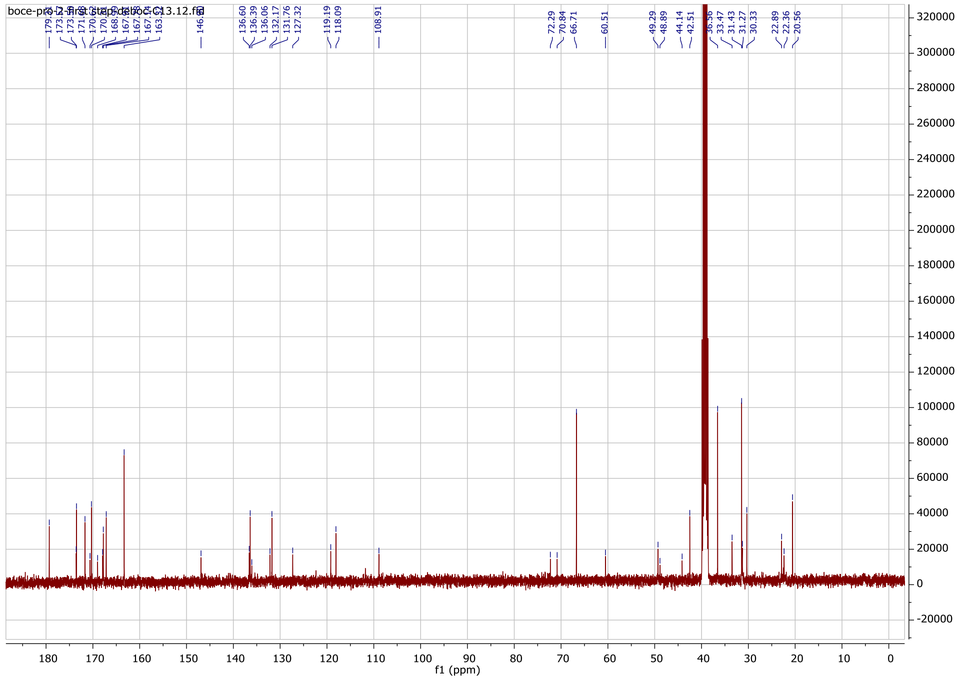
boce-pro-2-first-step-deboc-C13.12.fid
f1 (ppm)

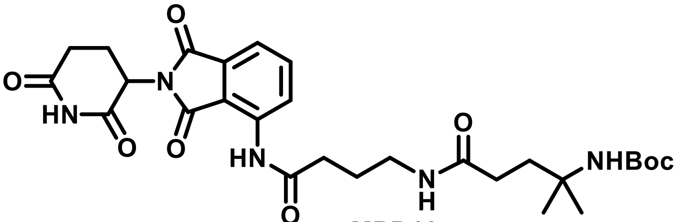

**MPD10e**

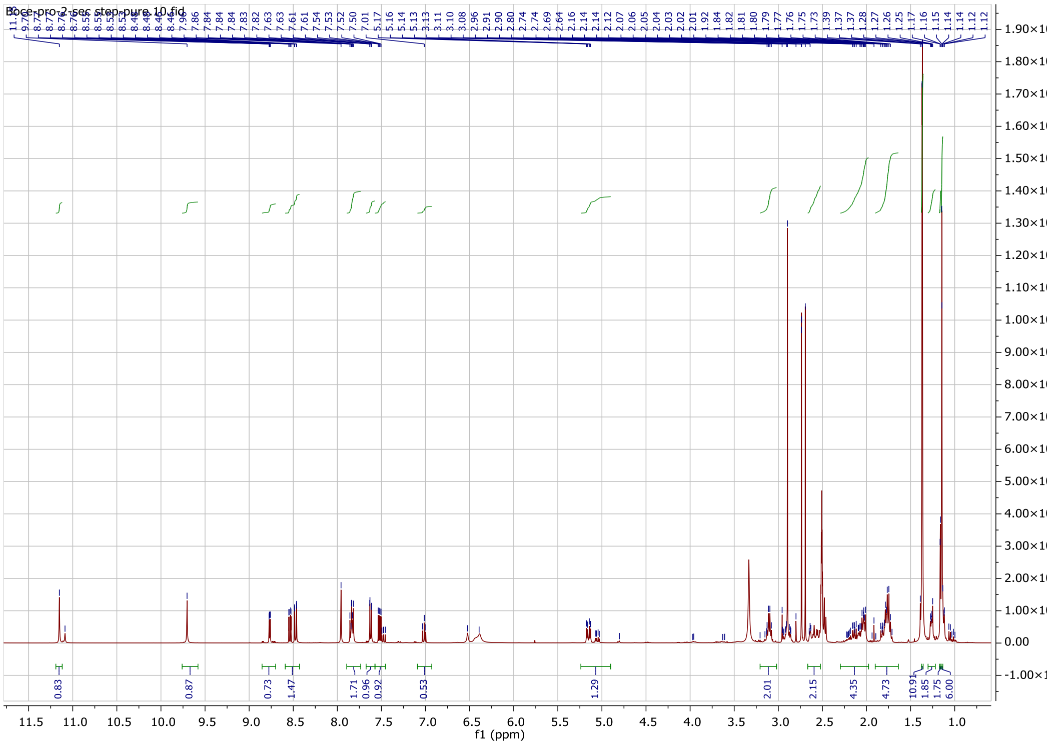

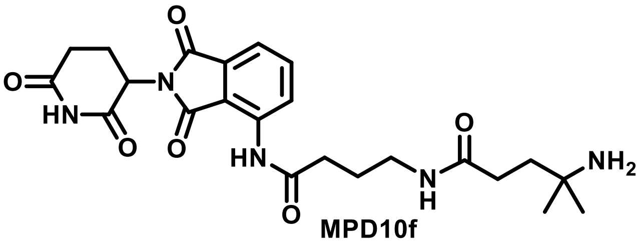

**MPD10f**

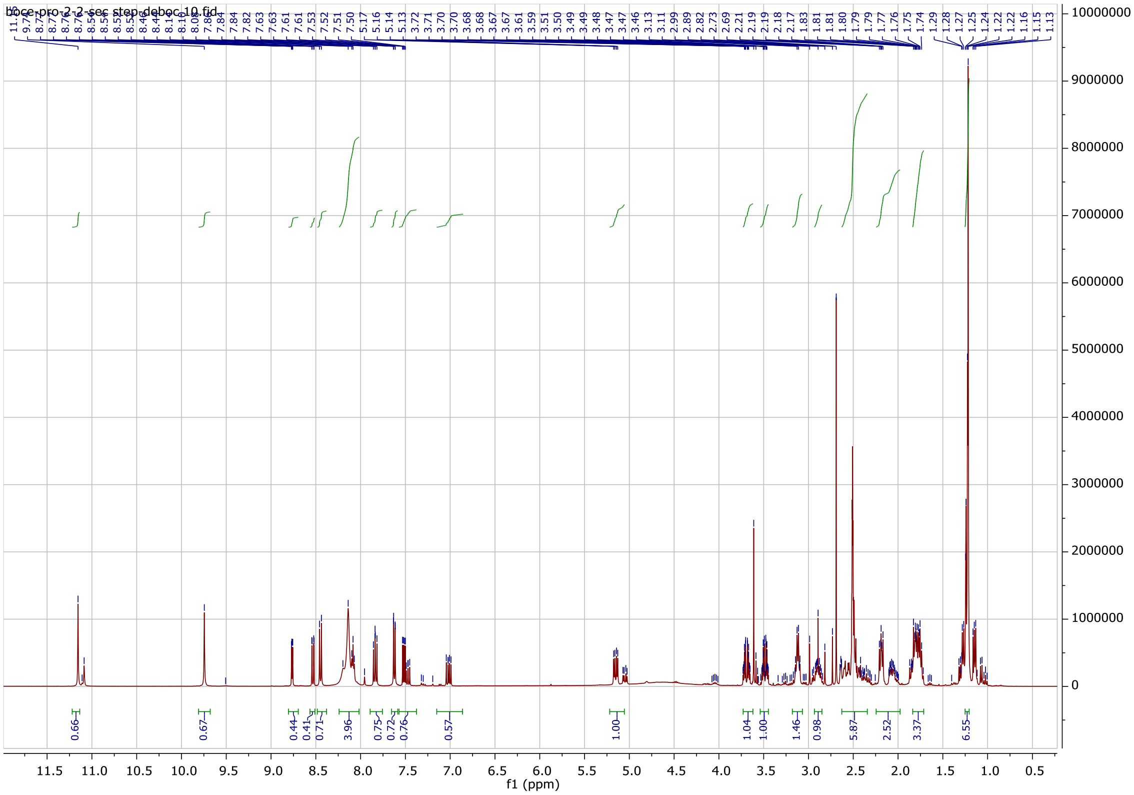

f1 (ppm)
11.5 11.0 10.5 10.0 9.5 9.0 8.5 8.0 7.5 7.0 6.5 6.0 5.5 5.0 4.5 4.0 3.5 3.0 2.5 2.0 1.5 1.0 0.5
10000000 9000000 8000000 7000000 6000000 5000000 4000000 3000000 2000000 1000000 0
0.66 0.67 0.44 0.41 0.71 3.96 0.75 0.72 0.76 0.57 1.00 1.04 1.00 1.46 0.98 5.87 2.52 3.37 6.55

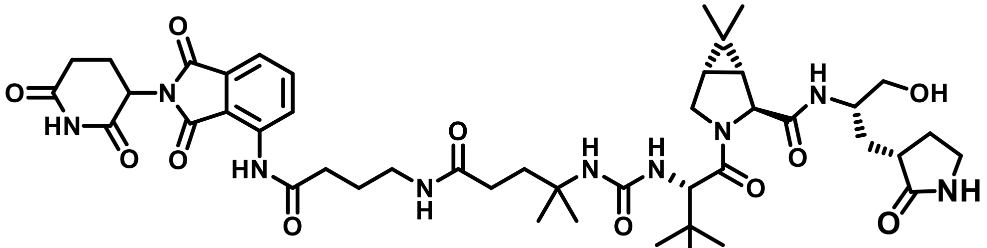

MPD10g

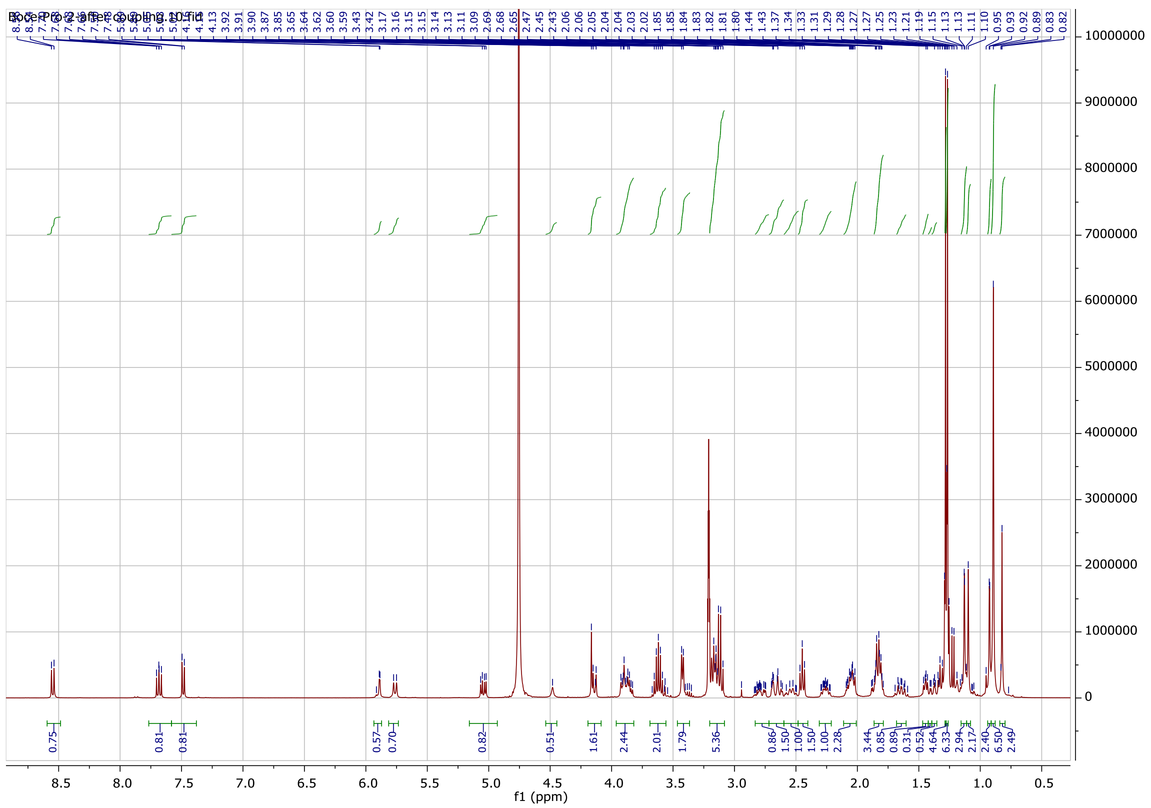

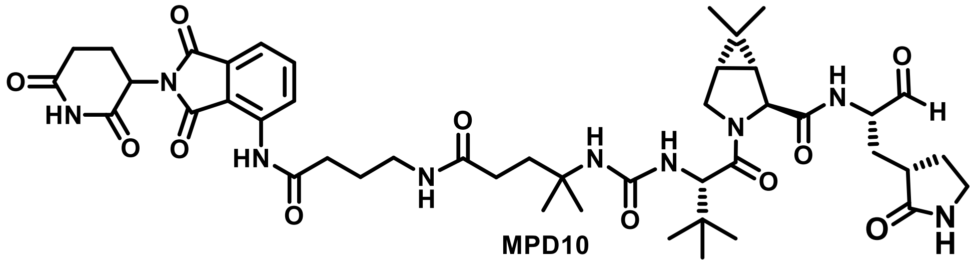

MPD10

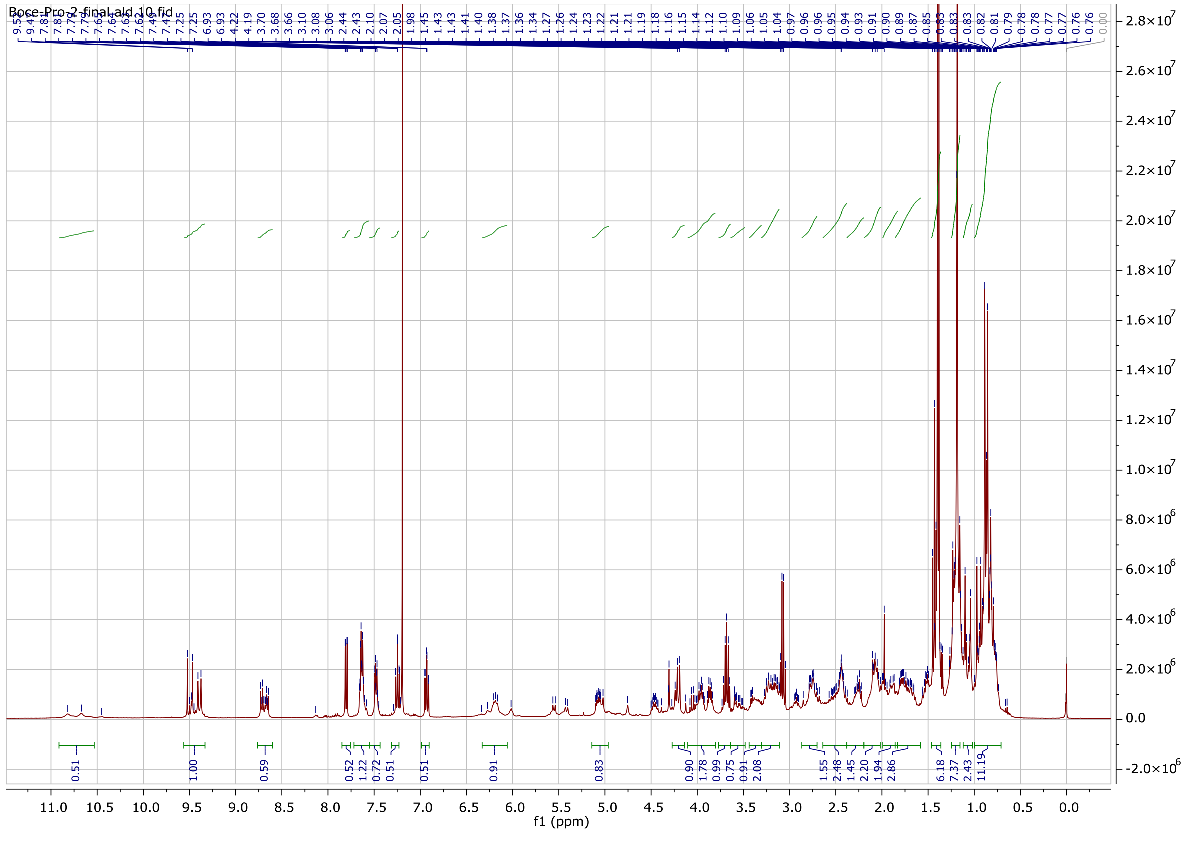

### LC-MS data of MPDs

### Control (1:1 DMSO: ACN by volume)

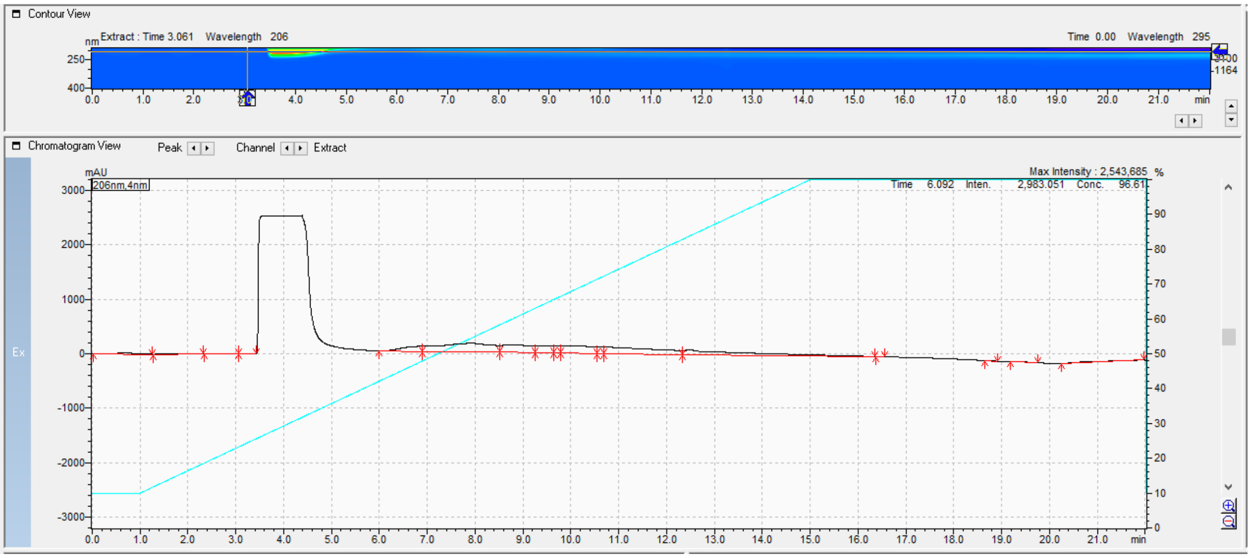

### MPD1

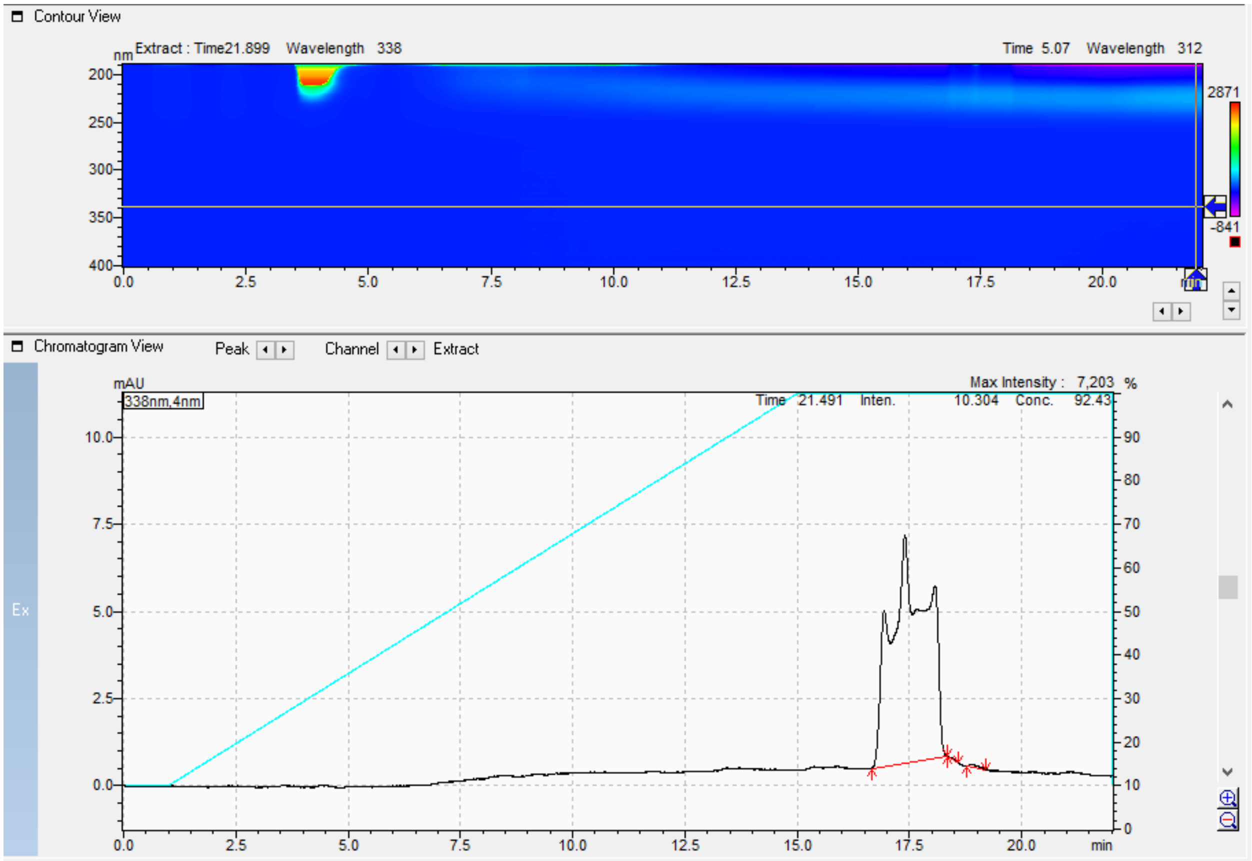


Results View - Peak Table

Peak Table | Compound | Group | Calibration Curve

| Peak# | Ret. Time | Area | Height | Mark | Conc. | Unit | ID# |
|---|---|---|---|---|---|---|---|
| 1 | 17.397 | 359198 | 6562 | M | 99.651 | | |
| 2 | 18.372 | 397 | 21 | M | 0.110 | | |
| 3 | 18.884 | 862 | 62 | M | 0.239 | | |
| Total | | 360458 | 6644 | | 100.000 | | |

### MPD2

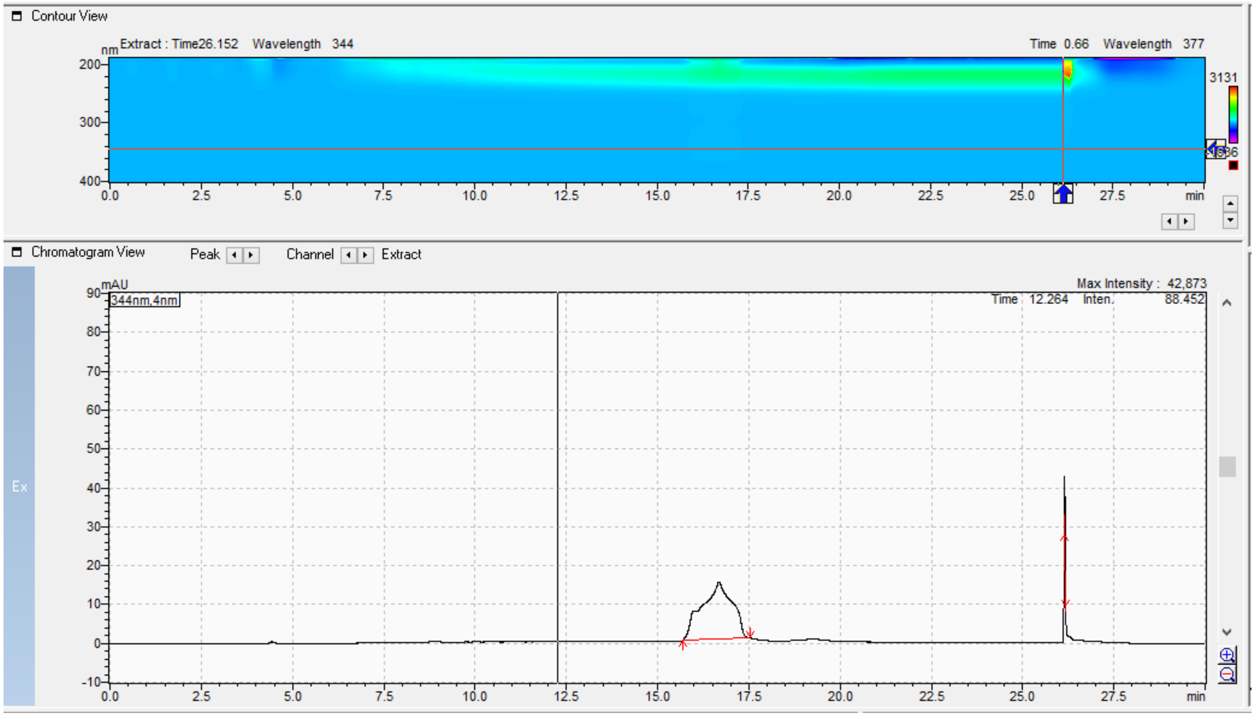


Results View - Peak Table

Peak Table | Compound | Group | Calibration Curve

| Peak# | Ret. Time | Area | Height | Mark | Conc. | Unit | ID# |
|---|---|---|---|---|---|---|---|
| 1 | 16.684 | 847865 | 14683 | M | 97.766 | | |
| 2 | 26.153 | 19370 | 16928 | M | 2.234 | | |
| Total | | 867235 | 31611 | | 100.000 | | |

**MPD3**

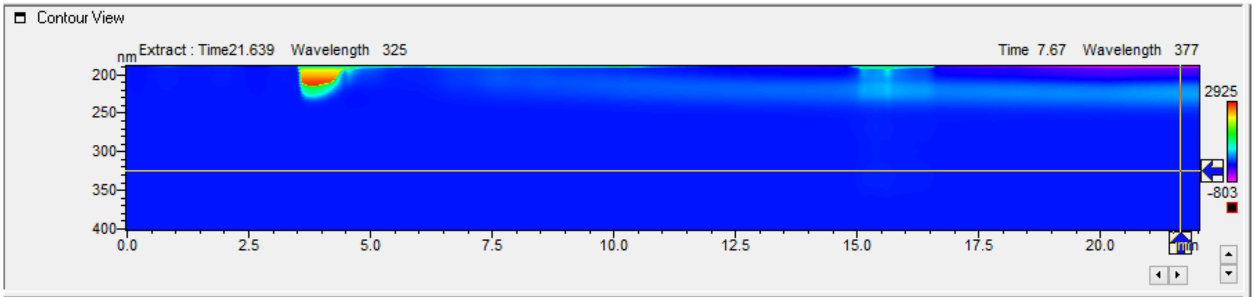


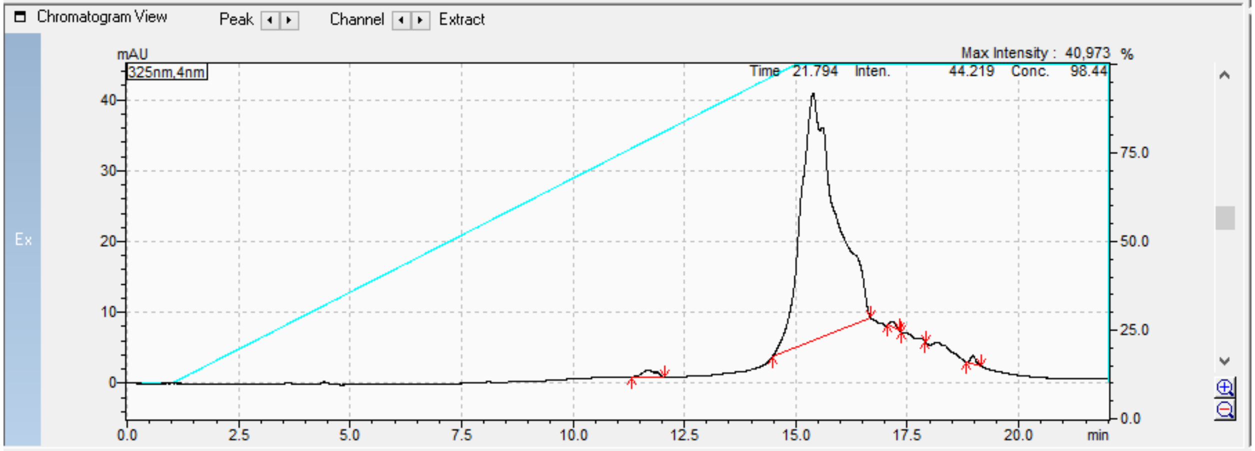


Results View - Peak Table

Peak Table | Compound | Group | Calibration Curve

| Peak# | Ret. Time | Area | Height | Mark | Conc. | Unit | ID# |
|---|---|---|---|---|---|---|---|
| 1 | 11.681 | 20846 | 945 | M | 1.074 | | |
| 2 | 15.385 | 1902312 | 34927 | M | 98.051 | | |
| 3 | 17.183 | 8017 | 863 | M | 0.413 | | |
| 4 | 17.369 | -15 | 3 | M | -0.001 | | |
| 5 | 17.897 | -45 | 2 | M | -0.002 | | |
| 6 | 18.972 | 9002 | 1154 | M | 0.464 | | |
| Total | | 1940119 | 37894 | | 100.000 | | |

### MPD4

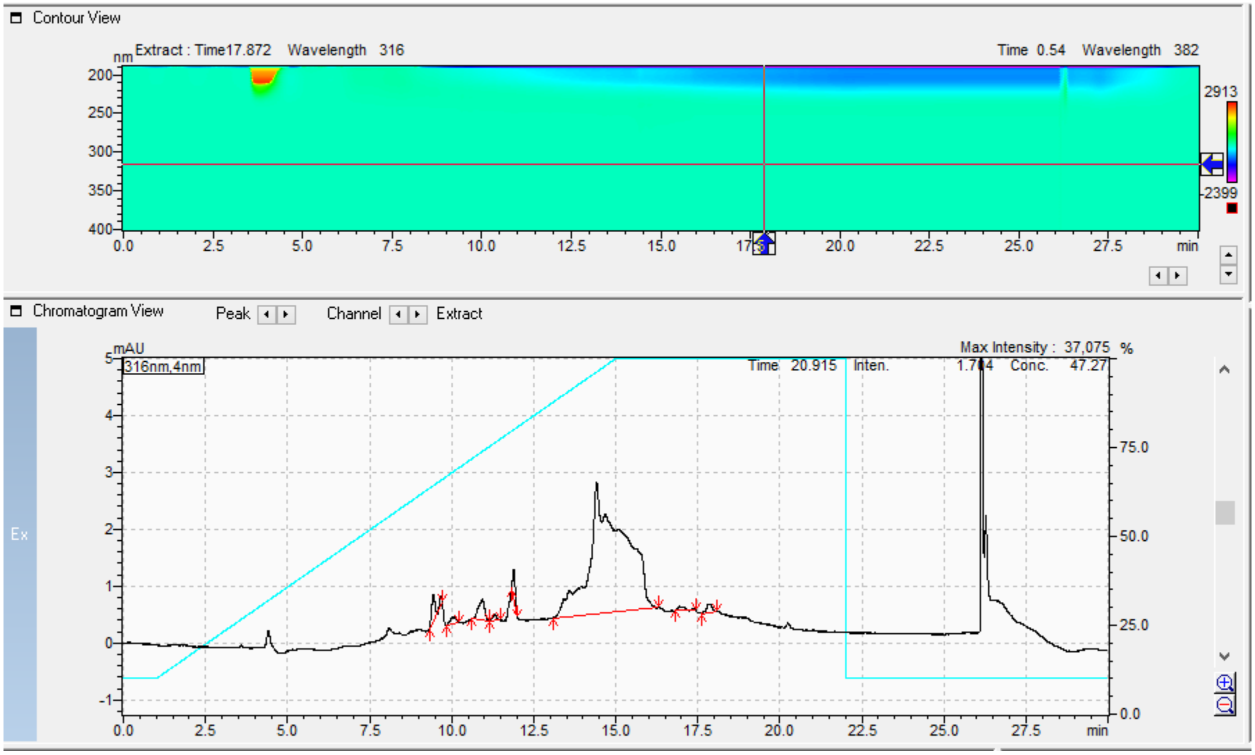


Results View - Peak Table

Peak Table | Compound | Group | Calibration Curve

| Peak# | Ret. Time | Area | Height | Mark | Conc. | Unit | ID# |
|---|---|---|---|---|---|---|---|
| 1 | 9.445 | 3819 | 456 | M | 2.098 | | |
| 2 | 10.046 | 1433 | 112 | M | 0.787 | | |
| 3 | 10.932 | 4854 | 366 | M | 2.667 | | |
| 4 | 11.311 | 951 | 106 | M | 0.522 | | |
| 5 | 11.894 | 2614 | 496 | M | 1.436 | | |
| 6 | 14.419 | 165492 | 2308 | M | 90.924 | | |
| 7 | 16.948 | 1156 | 70 | M | 0.635 | | |
| 8 | 17.872 | 1693 | 145 | M | 0.930 | | |
| Total | | 182012 | 4059 | | 100.000 | | |

### MPD5

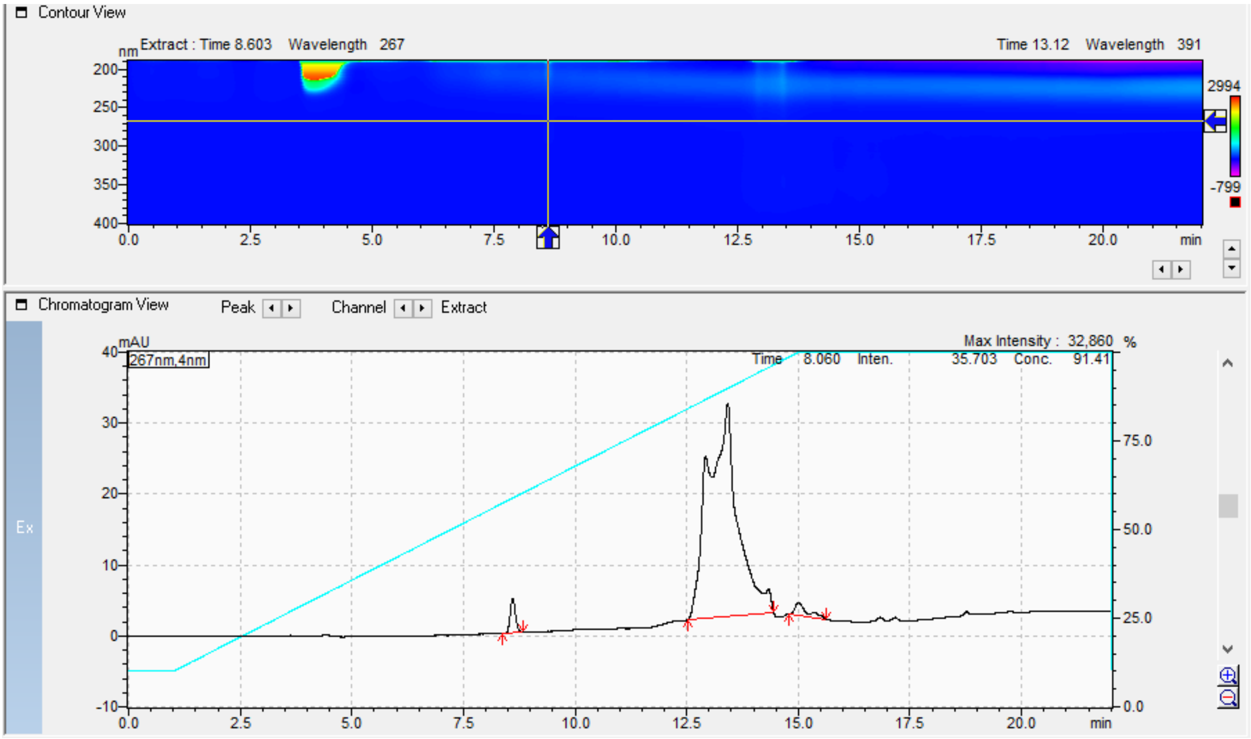


Results View - Peak Table

Peak Table | Compound | Group | Calibration Curve

| Peak# | Ret. Time | Area | Height | Mark | Conc. | Unit | ID# |
|---|---|---|---|---|---|---|---|
| 1 | 8.603 | 39498 | 4806 | M | 2.721 | | |
| 2 | 13.420 | 1378158 | 30085 | M | 94.958 | | |
| 3 | 15.006 | 33684 | 1799 | M | 2.321 | | |
| Total | | 1451339 | 36690 | | 100.000 | | |

**MPD6**

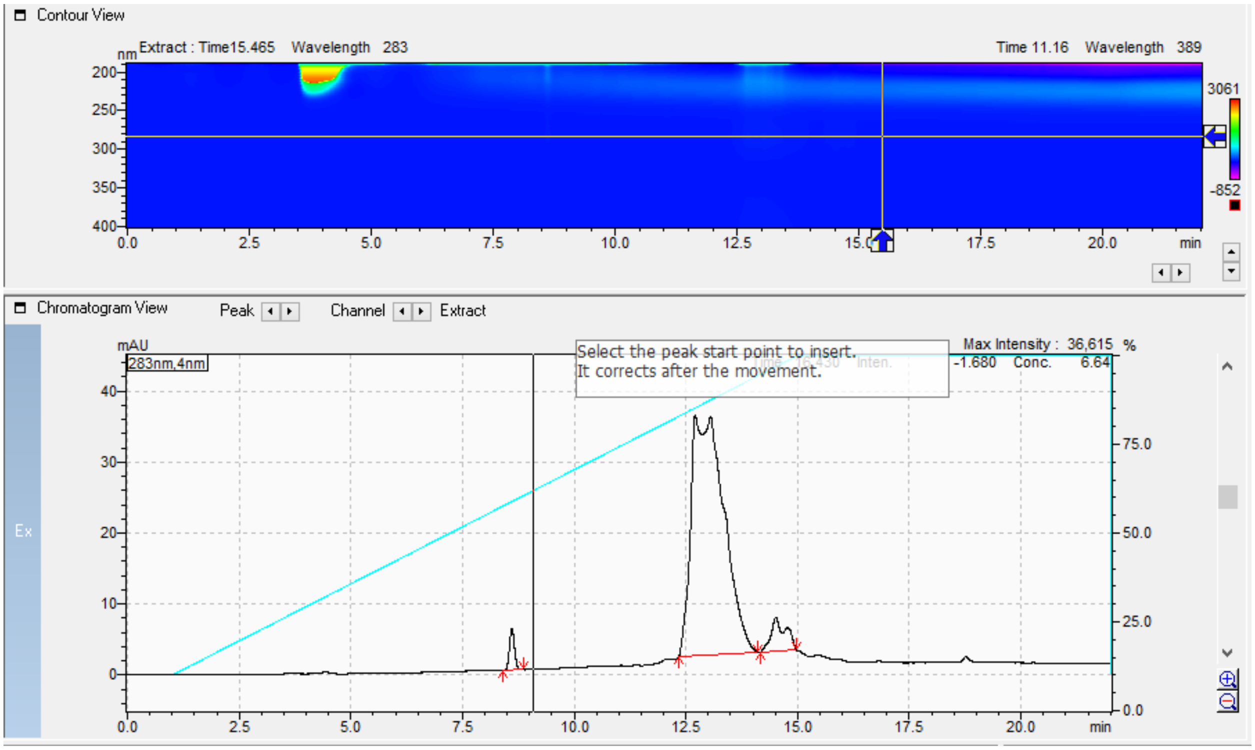


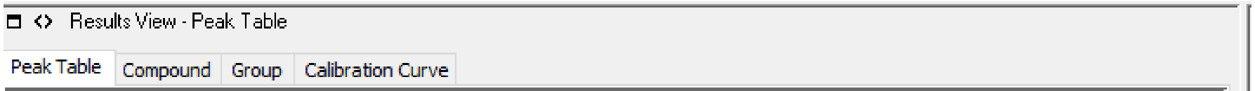


| Peak# | Ret. Time | Area | Height | Mark | Conc. | Unit | ID# |
|---|---|---|---|---|---|---|---|
| 1 | 8.605 | 44530 | 5711 | M | 2.350 | | |
| 2 | 12.706 | 1752405 | 34108 | M | 92.484 | | |
| 3 | 14.513 | 97884 | 4435 | M | 5.166 | | |
| Total | | 1894819 | 44255 | | 100.000 | | |

### MPD7

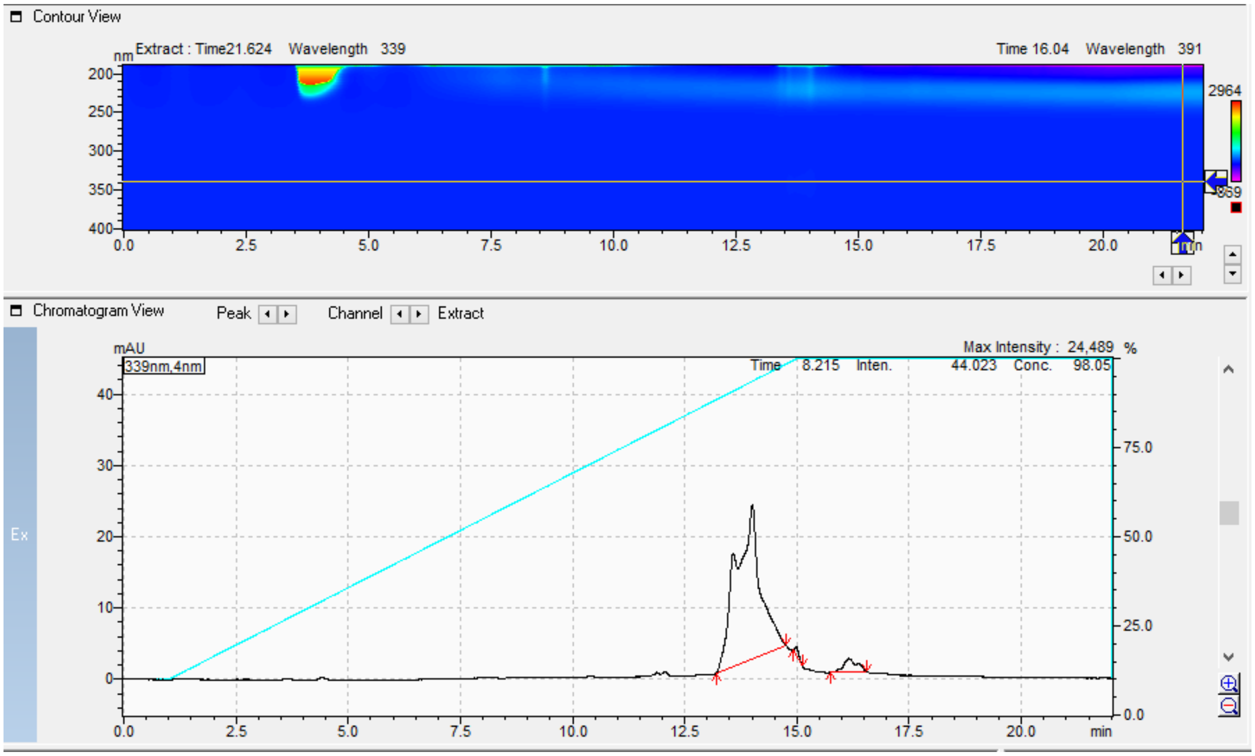


Results View - Peak Table

Peak Table | Compound | Group | Calibration Curve

| Peak# | Ret. Time | Area | Height | Mark | Conc. | Unit | ID# |
|---|---|---|---|---|---|---|---|
| 1 | 14.008 | 821465 | 21582 | M | 94.515 | | |
| 2 | 14.978 | 7437 | 1094 | M | 0.856 | | |
| 3 | 16.152 | 40236 | 1905 | M | 4.629 | | |
| Total | | 869139 | 24581 | | 100.000 | | |

### MPD8

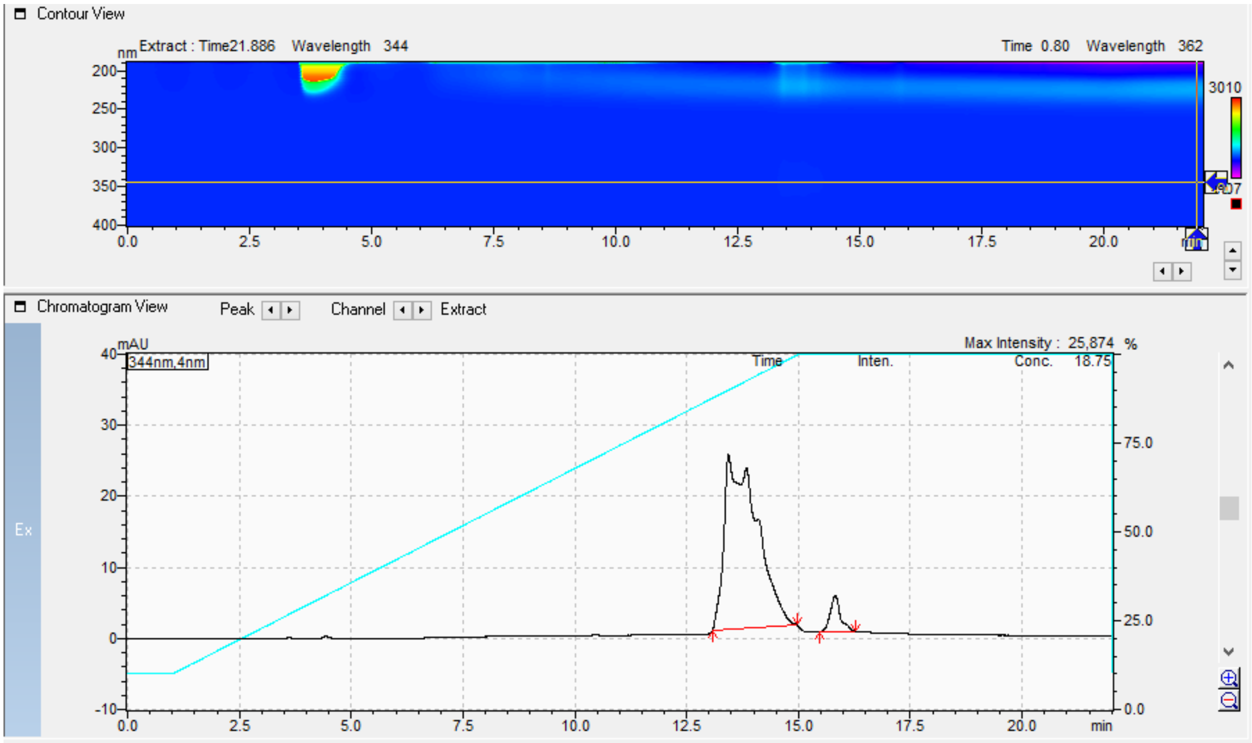


Results View - Peak Table

Peak Table | Compound | Group | Calibration Curve

| Peak# | Ret. Time | Area | Height | Mark | Conc. | Unit | ID# |
|---|---|---|---|---|---|---|---|
| 1 | 13.435 | 1247120 | 24610 | M | 93.911 | | |
| 2 | 15.822 | 80856 | 5057 | M | 6.089 | | |
| Total | | 1327976 | 29666 | | 100.000 | | |

**MPD9**

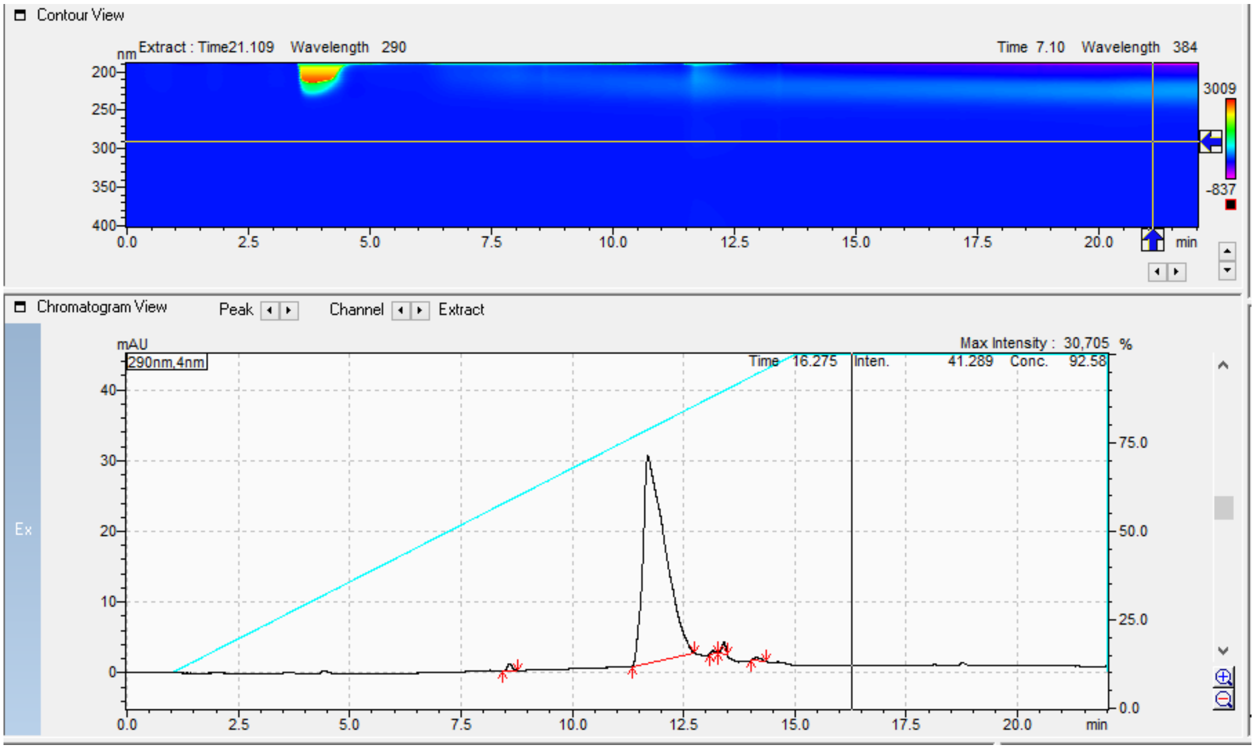


Results View - Peak Table

Peak Table | Compound | Group | Calibration Curve

| Peak# | Ret. Time | Area | Height | Mark | Conc. | Unit | ID# |
|---|---|---|---|---|---|---|---|
| 1 | 8.595 | 7314 | 947 | M | 0.708 | | |
| 2 | 11.696 | 1007580 | 29277 | M | 97.514 | | |
| 3 | 13.165 | 2693 | 448 | M | 0.261 | | |
| 4 | 13.406 | 10317 | 1661 | M | 0.998 | | |
| 5 | 14.125 | 5366 | 460 | M | 0.519 | | |
| Total | | 1033270 | 32793 | | 100.000 | | |

**MPD10**

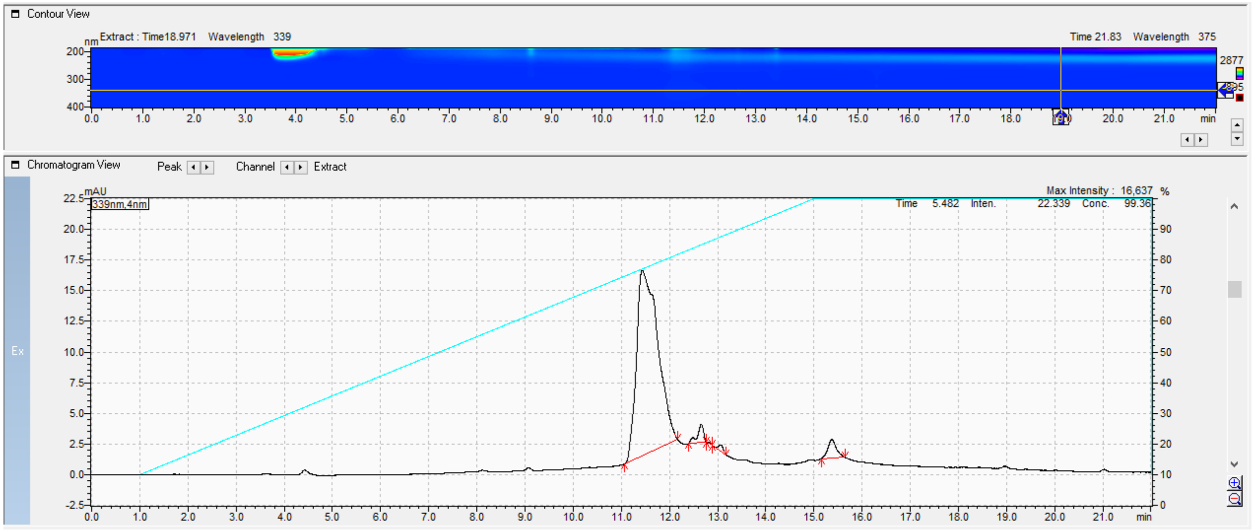

Results View - Peak Table

Peak Table | Compound | Group | Calibration Curve

| Peak# | Ret. Time | Area | Height | Mark | Conc. | Unit | ID# |
|---|---|---|---|---|---|---|---|
| 1 | 11.434 | 462891 | 15099 | M | 92.940 | | |
| 2 | 12.660 | 13380 | 1479 | M | 2.686 | | |
| 3 | 12.832 | 314 | 120 | M | 0.063 | | |
| 4 | 13.066 | 2727 | 496 | M | 0.548 | | |
| 5 | 15.371 | 18742 | 1538 | M | 3.763 | | |
| Total | | 498054 | 18731 | | 100.000 | | |
